# A near-full-length HIV-1 genome from 1966 recovered from formalin-fixed paraffin-embedded tissue

**DOI:** 10.1101/687863

**Authors:** Sophie Gryseels, Thomas D. Watts, Jean-Marie M. Kabongo, Brendan B. Larsen, Philippe Lemey, Jean-Jacques Muyembe-Tamfum, Dirk E. Teuwen, Michael Worobey

## Abstract

Although estimated to have emerged in humans in Central Africa in the early 1900s, HIV-1, the main causative agent of AIDS, was only discovered in 1983. With very little direct biological data of HIV-1 from before the 1980s, far-reaching evolutionary and epidemiological inferences regarding the long pre-discovery phase of this pandemic are based on extrapolations by phylodynamic models of HIV-1 genomic sequences gathered mostly over recent decades. Here, using a very sensitive multiplex RT-PCR assay, we screened 1,652 formalin-fixed paraffin-embedded tissue specimens collected for pathology diagnostics in Kinshasa, Democratic Republic of Congo (DRC), between 1959 and 1967. We report the near-complete genome of one positive from 1966 (“DRC66”)—a non-recombinant sister lineage to subtype C that constitutes the oldest HIV-1 near-full-length genome recovered to date. Root-to-tip plots showed the DRC66 sequence is not an outlier as would be expected if dating estimates from more recent genomes were systematically biased; and inclusion of DRC66 sequence in tip-dated BEAST analyses did not significantly alter root and internal node age estimates based on post-1978 HIV-1 sequences. There was larger variation in divergence time estimates among datasets that were subsamples of the available HIV-1 genomes from 1978-2015, showing the inherent phylogenetic stochasticity across subsets of the real HIV-1 diversity. In conclusion, this unique archival HIV-1 sequence provides direct genomic insight into HIV-1 in 1960s DRC, and, as an ancient-DNA calibrator, it validates our understanding of HIV-1 evolutionary history.

**Significance:** Inferring the precise timing of the origin of the HIV/AIDS pandemic is of great importance because it offers insights into which factors did—or did not—facilitate the emergence of the causal virus. Previous estimates have implicated rapid development during the early 20^th^ century in Central Africa, which wove once-isolated populations into a more continuous fabric. We recovered the first HIV-1 genome from the 1960s, and it provides direct evidence that HIV-1 molecular clock estimates spanning the last half-century are remarkably reliable. And, because this genome itself was sampled only about a half-century after the estimated origin of the pandemic, it empirically anchors this crucial inference with high confidence.

## Introduction

The human immunodeficiency virus (HIV) pandemic is one of the most devastating in all human history: more than 70 million people have so far been infected with the virus and over 35 million have died (1). Though AIDS and its main causative agent HIV-1 were discovered almost four decades ago, the virus has likely been circulating for about a century. In the time before its discovery, several strains of the virus had already spread from Central Africa to far corners of the world and diversified into the different subtypes we classify the virus into today. These strains belong to HIV-1 group M (2), which is set apart from the other HIV strains causing AIDS in humans—HIV-1 group N, O, P and the eleven HIV-2 groups—by its pandemic scale and much higher prevalence, accounting for >98% of all cases (1).

As a retrovirus with an RNA genome, HIV-1 evolves so rapidly that sequence data collected at various time points can calibrate molecular clock models used to infer the timing of historical, unsampled events in the virus’s past. However, rates of molecular evolution appear to vary depending on the time frame considered, so rates estimated over a recent time frame can not necessarily be extrapolated to a deeper time frame (3, 4). For example, molecular clock estimates based on the time that endogenous lentiviruses integrated in hosts’ genomes are several orders of magnitude slower than clock rates estimated from time-stamped extant samples. Clock rates measured with recent HIV and simian immunodeficiency virus (SIV) samples are also not reconcilable with the limited divergence between SIV strains in monkeys on an island established >10,000 years ago and SIV strains in the mainland relatives (5).

While time-dependent bias appears to grow in a continuous fashion with larger time frames (3), different time frames reflect different biological processes (4). For RNA viruses, these include differences in host-induced selection pressures (6) and differences in population dynamics, for example when comparing intra-host to inter-host dynamics (7) or newly-emerged outbreaks versus endemically circulating viruses (8). As such, it should be possible to demarcate time-frames with consistent epidemic behavior and therefore likely also consistent average molecular clock rates.

A crucial aspect in estimating molecular clock rates across such longer time frames is calibration by known events in the past (9). While ancient genomic DNA or RNA sequenced from archival samples or hosts remains are only available for relatively recent evolutionary times, they represent powerful calibrators as they offer direct evidence of an organism’s presence and identity at a point in the past. Short HIV-1 RNA fragments belonging to two different subtypes isolated from archival tissue specimens from 1959 and 1960 provided a upper limit for how long HIV-1 had circulated in people and when the diversification of major lineages occurred (10).

The wealth of genetic sequences of HIV-1 sampled throughout the decades since its discovery plus the few pre-discovery archival sequences led to the phylogenetic inference of the time of the most recent common ancestor (TMRCA) of HIV-1 group M around 1920 in Central Africa (10–13). The original spillover event of HIV-1 group M from central chimpanzees to humans must have occurred in rural south-east Cameroon (14) and, assuming the estimates of the TMRCA are correct, it must have occurred at or before this point in time. Early divergence events that resulted in the diversity of HIV-1 group M we see today likely took place in the major—and at that time massively growing—cities on the banks of the Congo river in what is now Republic of Congo (ROC) and Democratic Republic of Congo (DRC) (10, 11). This is also apparent from the much higher diversity of HIV-1 group M in DRC, ROC and Cameroon, with divergent lineages and circulating recombinant forms (CRFs) still being described from that region (15–18).

Genetic sequences from before HIV-1’s discovery in 1983 are scarce because 1) few archival specimens exist from that time period, 2) for those specimens that exist, odds are strongly against retrospectively finding a patient’s samples with an unspecific syndrome, 3) while screening by PCR amplification may be the most sensitive technique, sensitive existing primer sets might miss unknown divergent strains, 4) the further back in time one looks, the lower the expected prevalence of the virus, and 5) genetic molecules, especially viral RNA, tend to degrade during long-term storage or degrade during the original preservation of the specimen. Indeed, most long-term stored specimens are initially intended for histology, and while afterwards remaining fairly stable at room temperature, the original formalin-fixation process degrades RNA and DNA tremendously. However, short viral DNA or RNA sequences around 50-200 bases in length are still recoverable from such formalin-fixed-paraffin-embedded (FFPE) samples when using dedicated ultra-sensitive amplification techniques (10, 19–22).

Our lab recently described a particularly sensitive procedure, the “jackhammer RT-PCR” to detect and obtain high-quality, potentially divergent viral genomic sequences, yielding 8 of the 9 oldest HIV-1 group M genomes characterized to date (23).

Here, we screened 1,652 FFPE samples from central Africa using a jackhammer RT-PCR procedure modified to accommodate the small size of RNA fragments present in such specimens. We determined the near-full-length genome of an HIV-1 positive sample from Kinshasa, DRC, from 1966, the earliest HIV-1 genome assembled to date. We further investigated the effects of including or excluding this genome as a calibration point from deep within the pre-discovery phase of HIV/AIDS on the emergence and divergence times estimates of the HIV pandemic.

## Methods

See supplement for a more detailed description of the methods.

We screened a total of 1,652 FFPE samples from various tissues, originating from the DRC from 1959-67 Data recorded were unlinked to individual identifiers and the work was approved by the Human Subjects Protection Program at the University of Arizona. Paraffin was dissolved with xylene and total RNA extracted using the High Pure FFPE RNA Micro Kit (Roche, Indianapolis, IN) or Qiagen miRNeasy FFPE kit. The eight HIV-1 screening primer pairs listed in Supplementary Materials, Table S1, target regions 72 and 112 nucleotides (nt) in length of the *gag*, *pol* and *env* genes plus a human *beta actin* positive control. Reverse transcription was carried out for each RNA sample in 8-fold multiplex and a ‘pre-amplification’ step for each resulting cDNA sample was then carried out in 8-fold multiplex by adding a pool of the forward primers so that extremely rare templates molecules are not lost during aliquoting (23). In the final amplification step, each of the reactions was carried out as a single-plex PCR reaction with each primer pair. Products were cloned and then sequenced at the University of Arizona Genetics Core facility.

FFPE tissue from a patient sampled in 1966 was HIV-1 positive. We subjected this HIV-1 sample, henceforth called “DRC66”, to jackhammer-PCR attempts to sequence its coding genome. We designed 124 pan-HIV-1 group M primer pairs. Of these, 23 yielded HIV sequences, which were indicative of a subtype C-like genome. We then designed 220 primer pairs for subtype C-like genomes, of which 120 yielded HIV sequences. We closed most remaining gaps using a primer-walking approach for which a total of 446 primers were designed. All PCRs were organized in a jackhammer approach. Sequences of all used primers are available in the Supplementary Materials, Table S4. All DRC66 amplicon sequences were assembled and a consensus sequence is deposited in GenBank with Accession Number MN082768.

We searched for pre-existing drug resistance in DRC66 using the HIV Drug Resistance Database Program (24).

To compare our jackhammer PCR approach with a deep sequencing approach, paired-end sequencing was carried out on an Illumina HiSeq (see Supplementary Material). None of the reads mapped to HIV-1 but several bacterial, viral and fungal reads were recovered.

We searched extensively for both partial and full genomic sequences deposited in GenBank that had close similarity to the DRC66 genomic sequence. We built an alignment of the *pol* region sequences between 2485-4274 (HXB2 notation) for all found divergent subtype C sequences together with our subsampled dataset A (see below). We verified in RDP4 (25) that there had been no recombination among lineages within this alignment.

We then downloaded all non-recombinant near-complete (>7000 nt) HIV-1 group M genomes sampled in Africa from the LANL database on November 19 2018 (N=2342) and all HIV-1 group M genomes from other parts of the world that were sampled before 1985 (N=78). The REGA HIV subtyping tool v3.41 identified seven significant inter-subtype recombinants in this dataset, which we removed. Fourteen hypermutated sequences identified by Hypermut (26), were further removed. The final dataset contained 830 sequences, which was codon-aligned based on the translated gene sequences using the LANL HIVAlign tool (27) using the Hidden Markov Model (28). Intra-subtype recombination was investigated for sequences from each subtype separately in RDP4 (25). Sequence regions larger than 300 nt identified by at least six methods as involved in recombination were masked from the alignment. This alignment was further down-sampled in five distinct sub-alignments (named subsampled datasets A to E), for the purpose of efficient computation, as well as to assess the inherent uncertainty through phylogenetic variation among random samples of actual HIV-1 diversity.

Maximum-likelihood trees of the complete dataset and subsampled datasets were estimated with RAxML v8.2 and, to explore clock-like signal, root-to-tip distance was plotted against sampling time with R’s stats package. Tip-date calibrated phylogenetic trees were estimated using BEAST v. 10.4 (29) for each subsampled dataset A-E. All BEAST xml files are available in the supplement. Convergence and ESS values of all parameters were evaluated in Tracer v1.6. The first 20% of the Markov chain Monte Carlo runs were removed as burn-in and the three parameter log and trees log output files for each dataset were combined in R v3.4.2 and Logcombiner (29). These analyses were repeated but with 1) the DRC66 sequence removed from the alignment; and 2) without providing the sampling year of DRC66 but instead having its sampling date estimated from the rest of data using BEAST’s “tip date sampling” option and a uniform prior bound between 0 and 1000 (in years since youngest sampling date) (30). To explore how well tip dates of other samples could be estimated, we repeated the BEAST analyses of one of the subsampled alignments (A) for each of five different sequences in the dataset using BEAST’s “tip date sampling” option.

## Results

### HIV-1 genomic sequence from 1966 characterized via jackhammer PCR is the earliest known near-complete HIV genome

Out of 1,652 archival FFPE specimens from DRC dated between 1959 and 1967, lymph node tissue of a 38-year-old male biopsied in the year 1966 in Kinshasa, DRC, was found to be HIV-1 positive. No pathological annotations were associated with the specimens. We henceforth call this sample “DRC66”.

A large part of the genomic sequence was determined via PCR products yielding overlapping stretches of 54-110 nt genomic RNA, for which the RT and a pre-amplification step were efficiently carried out in pools of no more than 8 reactions while a final amplification step was carried out individually for each primer pair (collectively called the jackhammer PCR procedure). All final amplification products were cloned and at least 10 clones were Sanger sequenced. A final stretch covering 8347 bases of the genome (HXB2 positions 816 to 9175) was characterized, containing approximately 1959 undetermined sites (by comparison with HXB2 genome length), leaving a total of 6388 characterized nucleotide sites.

Analysis in REGA HIV subtyping tool and building RaXML phylogenies of 200-400nt alignments across manually determined sliding windows (results not shown) revealed that the entire DRC66 genome forms a sister lineage to the subtype C clade. There was no evidence for recombination with other HIV-1 group M subtypes or clades.

### Phenotypic characteristics

A comparison of V3 amino acid sequences between DRC66 and subtype C samples with known co-receptor usage (31) suggests that DRC66 would have been an R5 virus utilizing CCR5 co-receptor at the time of sampling.

Two or three known residues in the DRC66 of Integrase have been previously determined to confer resistance to integrase inhibitor drugs: H51Y, T66I and P145S. While H51Y and T66I mutations mildly reduce elvitegravir (EVG) susceptibility in patients, P145 induces high level resistance to EVG in vitro, though is rarely selected for in patients (24, 32, 33). The specific nucleotides that coded for the 51Y and 66I residues were present in 2/2 and 3/3 of successfully sequenced clones for those sites, but 145S was only present in 1/11 sequenced clones. A similar analysis of 568 *integrase* subtype C sequences sampled in Africa did not indicate any other strain harboring these specific three residues. No residues that confer resistance to Protease or Reverse Transcriptase inhibitor drugs were detected in the DRC66 sequence of those corresponding genes.

### Deep Illumina sequencing unsuccessful in uncovering HIV-1 sequence data

Illumina sequencing of this same sample yielded 68,847,162 paired reads, none of which mapped to HIV-1’s HXB2 genome, a subtype C genome (U46016) nor to the DRC66 consensus sequence generated by Sanger sequencing.

De novo assembly of reads into contigs >200 nt led to identification of mostly fungi and bacteria (see details in Supplementary Materials Table S2). These were most likely environmental organisms that we speculate entered the FFPE lymph node tissue sample during sample preparation in 1966 or during its long storage time since. Nevertheless, organisms that often represent opportunistic infections in AIDS patients were also detected, such as *Bartonella*, *Mycoplasma* and *Candida* (Table S2), though it cannot be excluded that these were derived from later-invading environmental species of these genera rather than being present in the patient’s original biopsy material. Identification of a fungus-borne virus (related to the known yeast virus *Saccharomyces cerevisiae* virus L-A) provides evidence that our deep sequencing approach was able to detect RNA viruses, although this virus contained a double-stranded RNA genome instead of the less stable ssRNA genomes of lentiviruses.

### DRC66 is a sister lineage to the subtype C clade and has extant relatives

The maximum likelihood tree of the complete dataset (830 non-recombinant genomes: 799 from Africa sampled between years 1983-2015; 31 from Europe and America sampled between years 1978-1985) reveals that DRC66 represents a sister lineage to the clade conventionally denoted as subtype C (Figure 1A). Extensive BLAST searches and searches via neighbor-joining trees of downloaded HIV-1 sequences revealed that DRC66 is the only reported near-complete genome from this lineage. However, the subtype-C like portions of the genomes of three recently described circulating-recombinant-forms (designated CRF93_CPX) from Kinshasa and Mbuji-Mayi, DRC, sampled in 2008 (15), formed a monophyletic group with DRC66 in a maximum likelihood tree of a partial *pol* alignment (Figure 1C). Partial *pol* sequences of other so-called “divergent C lineages” from CRF92_CU, described in Villabona *et al*. (15) and Rodgers *et al.* (17), were also included in this tree. The DRC66-CRF93_CPX clade is well supported, and as shown in Figure 1C, constitutes a sister clade to the rest of subtype C-like sequences. While the clade containing CRF92_CU is also well supported, the relationship between this clade, other “divergent C lineages” and the conventional subtype C cannot be recovered with confidence in this tree of partial *pol* sequences (Figure 1C).

**Figure 1.**
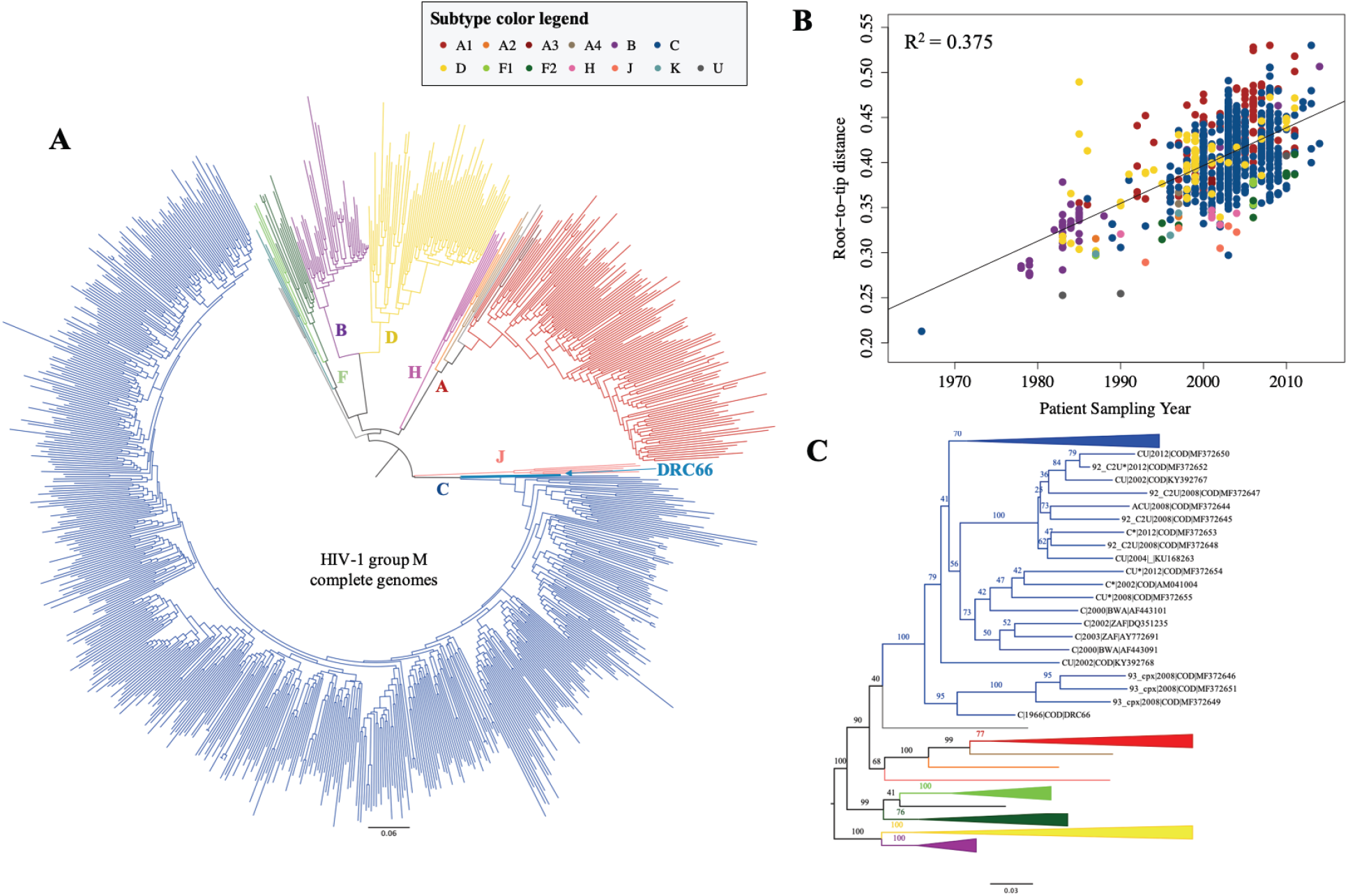
**A.** Mid-point rooted maximum likelihood tree of the complete dataset of 830 HIV-1 group M genomes. **B.** Evolutionary distances between the root and all tips of the tree shown in panel A plotted against year the sequence was sampled. **C.** Mid-point rooted maximum likelihood tree of a 1789 nt *pol* alignment that includes subsampled dataset A plus multiple “divergent subtype C-like” sequences, as reported in refs. (15) and (18) that are either derived from recombinant genomes (e.g. ‘CU’) or of which only partial sequence is available. Tips of the divergent subtype C-related sequences, including DRC66, are labeled by subtype (marked with * if determined based on partial sequence only, e.g. ‘C*’), sampling year, sampling country and GenBank accession number. For sampling country, COD=Democratic Republic of the Congo, BWA=Botwana, ZAF=Zambia.

### The root-to-DRC66-tip distance in ML tree is consistent with its sampling time

There is a good correlation between sampling years and root-to-tip distances in the ML tree of the complete, full genome dataset (R^2^=0.375) (Figure 1B). Although the R^2^ values of these regressions are difficult to interpret statistically, because of non-independence of the data points, they are indicative of the information held by the sampling dates on evolutionary rates. The root-to-tip distance of DRC66 is the shortest of all samples, as expected from the oldest sample in the dataset. The residual of −0.0413 of the DRC66 sample in this regression is small. Importantly, it is almost exactly the same as its residual (−0.0432) when calculating the regression without using the DRC66 data point but using all other root-to-tip data from the same ML tree (which yields a similar R^2^ of 0.361; data not shown). The DRC66 sequence is thus not an outlier in the root-to-tip regression plots. This indicates that clock-like signal from more-recently sampled HIV-1 genomes can reliably estimate dates of events from decades earlier.

Correlations between root-to-tip divergences and sampling times were very comparable between the full dataset of 830 genomes and the subsampled datasets of 176 or 177 genomes, or even improved in the subsampled datasets, indicating that the temporal signal is maintained when subsampling the dataset (Figure S1). R^2^ values were also very similar when estimating root-to-tip vs time correlations for phylogenies built from alignments that either included or excluded the DRC66 sequence (Figure S1).

### Inclusion of the DRC66 genome in BEAST time-stamped phylogenies indicates robust clock inferences using recent genomes

The five subsampled datasets were each subjected to three types of time-stamped analysis in BEAST (see Methods for details): 1) subsampled dataset including DRC66 and its sampling time; 2) subsampled dataset including DRC66 but leaving its sampling time unknown and to be estimated; 3) subsampled dataset excluding DRC66 sequence. Figure 2A shows a time-scaled phylogenetic BEAST tree of subsampled dataset A that includes DRC66 and its date; the time-scaled trees of subsamples B-E are displayed in Fig. S2 in the Supplementary Materials. Mean evolutionary rates and dating estimates are summarized in Table S3, Figure 2B, and Figure 3.

**Figure 2.**
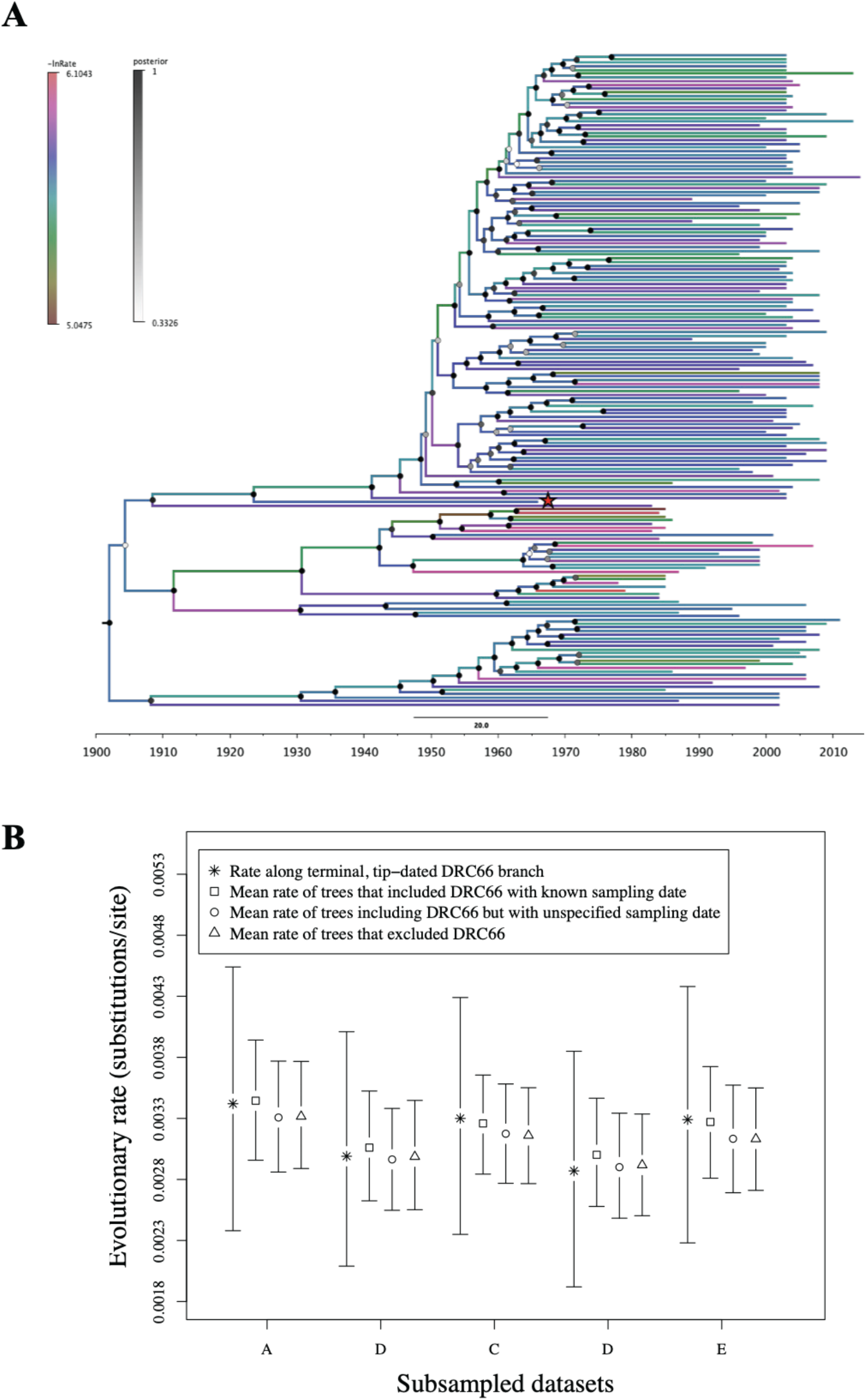
**A**. Time scaled phylogenetic BEAST tree of subsampled dataset A (trees of the other four subsampled dataset are displayed in Figure S2) estimated under a model that includes the sampling date of DRC66. Branches are color coded by log of the estimated evolutionary rate for that branch, drawn from a lognormal distribution using the uncorrelated relaxed clock model (34). Node labels are coded on a grey scale by the posterior probability of monophyly of the corresponding clade. The DRC66 sample is marked with a red star. **B**. For each of the five different subsamples, estimate of the evolutionary rate along the terminal branch leading to DRC66 in BEAST analyses that included DRC66’s sampling date (stars) and mean evolutionary rates across entire phylogenies for analyses that included DRC66 and its sampling date (squares), that included DRC66 but not its sampling date (circles), and that excluded DRC66 (triangles). Error bars represent 95% HPD intervals.

**Figure 3.**
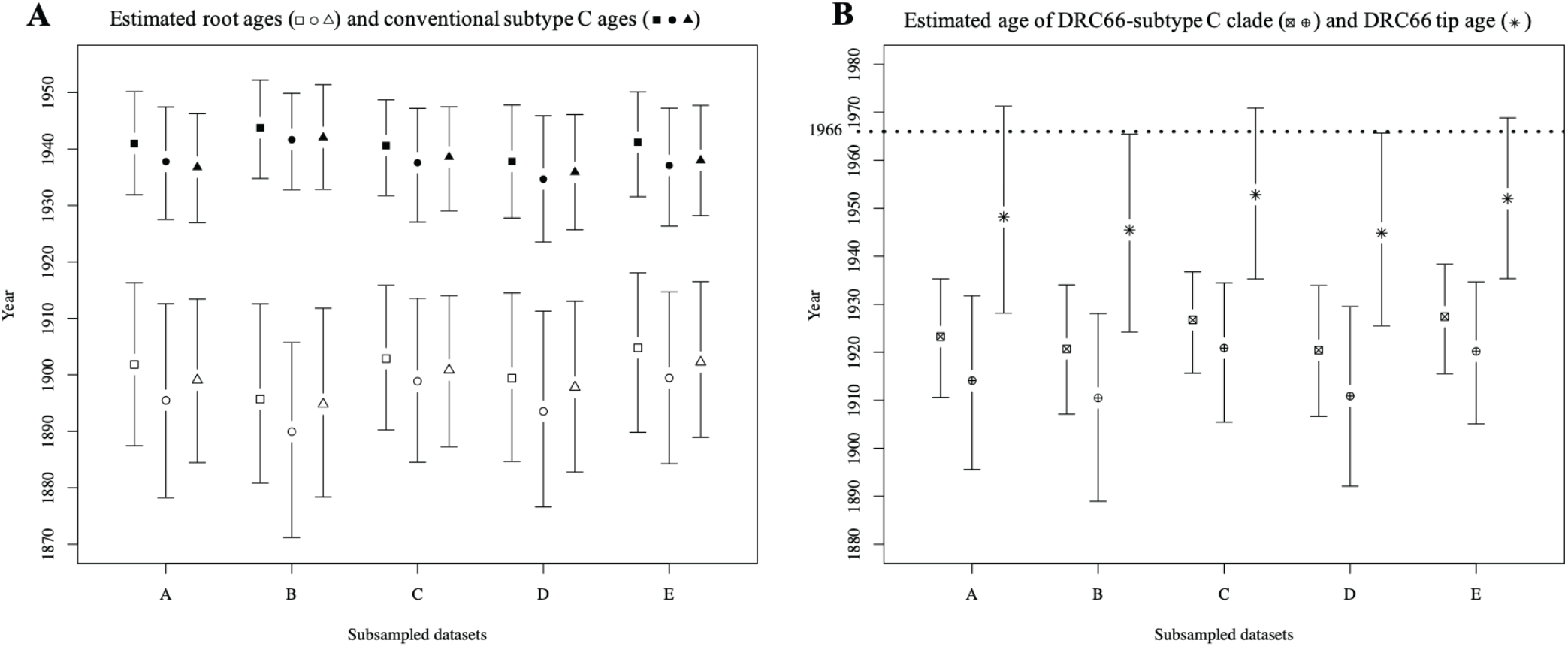
The means of different node ages and their 95% HPD intervals for time-scaled phylogenies of the five different subsampled datasets (A-E), which were each analysed in BEAST in three different ways: including DRC66 and its tip age (squares), including DRC66 but with its sampling date unknown and to be estimated (circles), and excluding DRC66 sequence (triangles). See also Table S3 in the supplement. **A**: age estimates of the root (open characters) and the node representing the common ancestor of conventional subtype C (filled characters). **B**: age estimates of the clade that encompasses both conventional subtype C and DRC66 (characters with crosses) and the estimated sample ages of DRC66 for those analyses in which this was left to be estimated (stars).

The posterior TMRCA estimates for the five different subsampled datasets A-E of HIV-1 group M genomes (that included DRC66 sequence and its age) ranged between 1896 (95% HPD 1881-1913) and 1905 (95% HPD 1890-1918) (Figure 3A, Table S3). The 95% highest probability distribution (HPD) intervals overlapped in all pairwise comparisons between subsampled datasets (Figure 3A, Table S3). While for each subsampled dataset the posterior mean estimates for the TMRCA were always slightly older when the DRC66 date was not specified and left estimated (between 4 and 6 years older) and when the DRC66 sequence was not included in the BEAST analyses (between 1 and 3 years older), this variation was smaller than the variation among some of the different subsampled datasets (Figure 3A, Table S3).

A similar pattern of little difference in age estimates between datasets including or excluding the DRC66 sample was observed for the node that represents MRCA of conventional subtype C (the subtype C clade that does not include DRC66), as well as for the node that represents the MRCA of the clade that incorporates both conventional subtype C and DRC66 (Figure 3B, Table S3).

In the BEAST analyses that included the DRC66 sample but left its sampling date to be estimated, the posterior means for the DRC66 tip date estimates were between 13 and 21 years older than the actual sampling year, and for 2 of 5 subsampled datasets the upper 95% HPD interval only just included the year 1966 (Figure 3B, Table S3). In subsequent BEAST analyses in which we estimated tip dates for five other HIV-1 sequences downloaded from GenBank, the posterior means for the tip date estimates were between 7 years younger and 39 years older than the real sampling dates (Table S3). The latter, with a mean tip date of 1954 (95% HPD 1933-1980), was the estimate for a divergent sequence of undetermined subtype (U) from 1983 and indicates that dates of the tips of long external branches are particularly difficult to estimate under a relaxed clock model.

## Discussion

Here we present what is currently the oldest near-complete HIV genome, from 1966 in Kinshasa, DRC. This DRC66 sample is 10 years older than the previously earliest characterized full genome, an 01A1G strain that was isolated from blood in 1976, also in DRC, but which underwent multiple cell culture passages before sequencing (35). There are only nine other HIV-1 genomes available from the pre-discovery phase of AIDS (1978-1982), all subtype B from the U.S. (36). The oldest HIV-1 genomic fragments are derived from plasma and FFPE samples from 1959 and 1960, again both from Kinshasa, DRC (10, 37). While these provided undisputable evidence of the presence and major diversification of HIV-1 group M two decades before its discovery, the short sequences that were recovered do not allow complete characterization of the HIV-1 strain involved and contain only a fraction of the phylogenetic information that is present in complete genomes.

To achieve sequence coverage across the DRC66 archival genome, labor-intensive amplification of overlapping short fragments between 54nt and 110nt in a highly sensitive jackhammer PCR procedure proved necessary. In comparison, none of the >65 million reads of an Illumina HiSeq run without prior amplification on the same sample contained HIV-1 sequence data. The latter approach had provided a full genome 3000x coverage of an influenza A H1N1 flu strain in an FFPE sample from 1918, however (22). Perhaps the difference in success resulted from different storage conditions in a humid tropical versus a temperate region, as evidenced by the majority of our reads being derived from environmental organisms that could have invaded the sample during preparation or storage, or, more likely, from a comparatively low viral titer in the FFPE lymph node specimen.

Globally, more HIV-1 group M cases are caused by strains that belong to the subtype C clade than any other clade, largely because southern Africa holds the highest HIV-1 burden and subtype C predominates there (38). Estimated to have originated in south-eastern DRC, phylodynamic analyses indicated subtype C strains have spread from there to southern Africa via connections between mining cities (11). Currently about 19% of HIV-1 infections in DRC are classified as subtype C (mostly documented from partial gene sequences). The DRC66 sequence represents a sister lineage to the subtype C clade, and quite divergent: we estimate it shared a common ancestor with subtype C some 20 to 30 years before the time of the common ancestor of conventional subtype C. Parts of *gag* and *pol* from three recently described recombinant genomes from Kinshasa and Mbuji-Mayi sampled in 2008 are the only contemporary sequences that also belong to this lineage in part of their genomes. Villabona *et al*. and Rodgers *et al*. describe two additional so-called “divergent C lineages” sampled between 2001-2012 that are monophyletic with conventional C with respect to the DRC66 lineage, yet form separate sister lineages to subtype C (15, 17). Similarly, for most other HIV-1 subtypes more divergent lineages can be found in DRC (in particular Kinshasa) and other central African countries than in other regions where the more restricted within-subtype diversity arose in a relatively short time after founder events. The DRC66 genome provides a unique insight into how the current conventional subtype C clade represents a fraction of the C-like diversity that was present in DRC in the 1960s. The fact that particular residues of the translated Integrase protein of DRC66 are known to induce resistance to integrase inhibitor drugs, which were obviously developed long after DRC66 was sampled, highlights that the natural 1960s diversity already harbored some genetic basis for anti-HIV therapy failure.

We further investigated whether the phylogenetic information in the suite of HIV-1 genomes sampled across the past decades, almost all after the discovery of HIV-1, reliably captures HIV-1’s evolutionary rates over the longer time frame that includes HIV-1’s long pre-discovery phase in humans. Few calibration points from direct biological observation are typically available to test such conclusions for real-world analyses, especially for such a medically important pathogen. Crucially, such ancient DNA calibration points can lead to dramatic changes in evolutionary histories once thought to be definitively established. For example, recently reported hepatitis B virus sequences from the Bronze age and Neolithic suggested a 100-fold slower evolutionary rate for this double-stranded DNA virus than previously thought (39–41), and these data are prompting updates to evolutionary clock models to better accommodate time-dependent rate variation (9). Because it is impossible to completely rule out such biases without complete genomic information from an early evolutionary time-point, we believe it is important to attempt to recover such information from surviving HIV-1 specimens.

Reassuringly, in the context of HIV-1 group M we do not observe that an ‘ancient’ HIV-1 genome significantly changes evolutionary inferences based on phylogenies built from more-recent genomes. Indeed, there is remarkably little difference in key estimates—including the overall age of the pandemic lineage of HIV—when this sequence is included in phylogenomic analyses. Given that it is more than 50 years older than currently circulating HIV-1 strains, this sequence provides direct evidence for the reliability of dating estimates over the last half-century of HIV-1 circulation. This stands in contrast to the disconnection between short-term rates observed in simian immunodeficiency viruses (SIV) and the rates at which SIV strains evolve when averaged across centuries or millennia of evolution in natural populations of different primate species, where molecular clock dating theory has difficulties accommodating the rate differences (5, 42).

Interestingly, our analysis highlights an often-overlooked source of uncertainty in evolutionary divergence dating based on any sample of genomes. The suite of HIV-1 genomes sampled from patients and available in public databases is inevitably a very limited subsample of the true diversity of HIV-1 group M. To investigate the degree of variation such an unavoidable sampling process induces, we subsampled the available GenBank sample of non-recombinant HIV-1 group M genomes from Africa. While credible intervals of all dating and rate estimates overlapped substantially, the overall variation between subsamples was larger than that induced in each subsample when DRC66 was either included or excluded. Besides variation in the underlying evolutionary models used in different studies, usage of different HIV-1 genome samples could also explain why our HIV-1 group M TMRCA estimates are somewhat older here than previously reported: 1920 (95% HPD 1909-1930) (11), 1930 (1911-1945) (43), 1932 (1905-1954) (13), 1920 (1902–1939) (13), and 1908 (1884–1924) (10). Variability in these TMRCA estimates among datasets indicates that an optimal dataset that fully captures HIV-1 group M’s natural evolutionary history over the past century has yet to be retrieved.

In conclusion, using a highly sensitive amplification protocol for degraded archival samples, we here present the oldest HIV-1 near-complete genome available to date. While we are careful not to extrapolate to other pathogen-host systems and much deeper time scales evident in SIV, our study indicates that evolutionary rates calibrated from HIV-1 group M sequences sampled across the decades after its discovery can be used reliably to infer the timing of events that occurred during the pre-discovery era. We note that the inherent stochasticity associated with a sample of the true viral diversity in nature inevitably introduces uncertainty to phylogenetic dating estimates, which is addressable by purposely subsampling datasets.

## Supporting information

Supplementary folder with BEAST xml files

## Acknowledgements

We would like to thank Tatenda Mangurenje and Ryan Ruboyianes for their excellent technical help.

This work was supported by NIH/NIAID R01AI084691 and the David and Lucile Packard Foundation (MW). SG was supported by an EMBO long term postdoctoral fellowship (ALTF-328) and an OUTGOING [Pegasus]^2^ Marie Skłodowska-Curie Fellowship of the Research Foundation - Flanders (“Fonds voor Wetenschappelijk Onderzoek – Vlaanderen”) (12T1117N) during this work. PL acknowledges funding from the European Research Council under the European Union's Horizon 2020 research and innovation programme (grant agreement no. 725422-ReservoirDOCS) and the Research Foundation - Flanders (G066215N, G0D5117N and G0B9317N).

