## Supplementary Materials for "A near-full-length HIV-1 genome from 1966 recovered from formalin-fixed paraffin-embedded tissue"

### Supplementary methods

#### Sample collection

We gathered a collection of 3,344 FFPE samples from DRC between 1959 and 1967. For 2,380 of these the tissue type was legibly recorded, and for 1,142 specimens a brief diagnosis was recorded. We selected 671 samples for which either the recorded diagnosis was consistent with immune deficiency and/or the tissue type was known to be HIV-1-tropic. These "high priority samples" and an additional, randomly chosen set of 981 samples were tested for the presence of HIV-1 RNA. This included 27 samples previously described in (1), but tested with a different PCR procedure. The total of 1,652 screened FFPE samples were mostly biopsies (N=998) rather than necropsies (N=72) or unknown (N=582). When known, tissues were most commonly derived from uterus (N=220), tumors (N=113), liver (N=72), lymph nodes (N=167), skin (N=60) and fallopian tubes (N=35). Data recorded were unlinked to individual identifiers and the work was approved by the Human Subjects Protection Program at the University of Arizona.

#### Screening FFPE samples

Small pieces of tissue were removed from the center of the FFPE block using a sterile scalpel. Paraffin was dissolved by adding 400µL of xylene, which was afterwards washed three times with ethanol. Samples were dried at 55°C, and subsequently total RNA was extracted using the High Pure FFPE RNA Micro Kit

(Roche, Indianapolis, IN) or Qiagen miRNeasy FFPE kit, according to manufacturer's instructions, though without the addition of carrier RNA.

The eight screening primer pairs are listed in Table S1. Seven primer pairs target regions between 72 and 112 nucleotides (nt) in length of *gag*, *pol* and *env* gene of HIV-1 and one primer pair targets human *beta actin*, which served as a positive control to indicate survival of amplifiable host RNA in the sample.

Reverse transcription was carried out for each RNA sample in 8-fold multiplex. A mixture of 3 μL of a pool of reverse primers at 10μM, 1μL of 10mM dNTP and 6μL of RNA extract was heated to 70°C for 5 minutes and chilled at 4°C. 10μL of the RNA extract was mixed with a mastermix consisting of 1μL GoScriptTM Reverse Transcriptase, 4μL GoScript buffer, 4 μL 25 mM Mg and 1μL of RNasin® Plus RNase Inhibitor (all Promega, Fitchburg, WI) and incubated for 30 min at 50°C followed by 30 min at 55°C for 4 cycles, then an incubation at 85°C for 10 minutes.

A 'pre-amplification' step for each resulting cDNA sample was then carried out in 8-fold multiplex by adding a pool of the forward primers. The purpose of pre-amplification is to increase the titer of target molecules in the RT reaction prior to aliquoting small amounts of material into individual PCR reactions, so that extremely rare templates molecules are not lost during aliquoting. 10μL of the RT product was added to a 40μL mastermix containing ingredients as specified for Accustart Taq DNA Polymerase HiFi (Quantabio), and run for 40 cycles using an annealing temperature of 52°C.

In the final amplification step, each of the reactions was carried out as a single-plex PCR reaction with each primer pair. First, 18μL of pre-amplified product was mixed with a 190μL mastermix containing ingredients as specified for Accustart Taq DNA Polymerase HiFi (Quantabio), and for each primer pair 23 μL of the previous was mixed with 2μL of both forward and reverse primers at 10μM. This was run for 40 cycles using an annealing temperature of 52°C.

We ran the final amplification products on a 2% agarose gel. Products that yielded bands close to expected heights were cloned with the pGEM-T Easy cloning kit (Promega, Fitchburg, WI) and Sanger sequenced at the University of Arizona Genetics Core facility.

**Table S1.** List of primers used in Jackhammer screening PCRs.

| Primer pair # | Name | Direction | Sequence | Approximate Tm | Amplicon length |
|---|---|---|---|---|---|
| 1 | gagA | F | 5'- TGCGAGAGCGTCAGTATTAAG-3' | 60 | 72nt |
| | | R | 5'- TTTCCCCCTGGCCKTAAC-3' | 56-58 | |
| 2 | gagB | F | 5'- TGGGAGAAAATTCGGTTAAGG-3' | 57 | 75nt |
| | | R | 5'- CTCCCTGCTTGCCCATACTA -3' | 60.3 | |
| 3 | envH | F | 5'- GTGTACCCACAGACCCCAAC -3' | 63.2 | 97nt |
| | | R | 5'- CTCATGCATTTGTTCTACCATGT -3' | 55.5 | |
| 4 | polFt | F | 5'- AGCTGGACTGTCAATGAYATACAGA-3' | 63-64 | 112nt |
| | | R | 5'- GCTTTGGYTCCCCTAAGG-3' | 56-58 | |
| 5 | B2M* | F | 5'- TGCTGTCTCCATGTTTGATGTATCT-3' | 60 | 85nt |
| | | R | 5'- TCTCTGCTCCCCACCTCTAAGT-3' | 60 | |

| | | | | | |
|---|---|---|---|---|---|
| 6 | pol45 | F | 5'- AGAAACATGGGARACATGG-3' | 57 | 78nt |
| | | R | 5'- GGAGGGGTATTRACAAAYTC-3' | 55 | |
| 7 | pol57 | F | 5'- CCAGCACAYAAAGGRATTGG-3' | 57 | 100nt |
| | | R | 5'- CCTGRGCYTTATCTATTCCA-3' | 57 | |
| 8 | pol60 | F | 5'- ATTGGAGARCAATGGCTAG-3' | 56 | 81nt |
| | | R | 5'- GCTGACAYTKATCACAGCT-3' | 55 | |

*human beta actin marker

#### Genome sequencing

In addition to the DRC60 FFPE sample we previously reported to contain HIV-1 (1), FFPE tissue from a patient sampled in 1966 was HIV-1 positive in the current study. We subjected this HIV-1 sample, henceforth called "DRC66", to jackhammer-PCR attempts to sequence its coding genome. The HIV-1 positive sample from 1960 detailed in (1) was mostly depleted and did not allow us to recover a complete genome from that sample in this study.

An initial HIV-1 genomic sequencing procedure with primers designed to work across the entire diversity of HIV-1 group M included a total of 124 primer pairs to be used in a jackhammer PCR approach. Forward and reverse primers were organized in separate pools of no more than eight that targeted non-overlapping regions of either *gag*, *env* or concatenated *vif*, *vpr* and *vpu*. We carried out the RT in multiplex (with a pool of reverse primers) and the pre-amplification in multiplex (with a pool of forward primers), as described above for the initial screening phase. A final amplification PCR was carried out for each primer pair separately, as described above. Gel-based detection, cloning and sequencing were carried out as described above. High quality Sanger sequences were obtained of an average of 10 clones per successful PCR product. Of these initial 124 PCRs, 23 yielded HIV sequences, which were indicative of a subtype C-like genome. A similar procedure was then carried out with a set of 220 primers designed to target the diversity of subtype C-like genomes. 120 of these 220 PCRs yielded HIV sequences.

We further attempted to close remaining uncharacterized gaps using a primer-walking approach for which a total of 446 primers were designed. These PCRs were again organized in a jackhammer approach.

Sequences of all used primers are listed in Table S4 at the bottom of this document.

All DRC66 amplicon sequences were assembled and aligned in MEGA v6 (2). A 95% base-matching consensus sequence was determined, in which variable sites were marked with ambiguity codes, and is deposited in Genbank with Accession Number MN082768.

#### Illumina sequencing and assembly

To compare our jackhammer PCR approach with a deep sequencing approach, the HIV-1 positive sample was also subjected to Illumina sequencing, using the protocol described in Xiao *et al.*, 2013, where it was successfully applied to yield 3000x genome coverage of pandemic H1N1 Influenza A from an FFPE sample from 1918 (3). In brief, a total RNA library was constructed using the mRNA-seq sample kit (Illumina, San Diego, CA) but without the poly-A selection and RNA fragmentation steps. Human rRNA presence in this library was reduced by duplex-specific nuclease digestion (4). Paired-end sequencing was carried out on an Illumina HiSeq.

Low-quality end of reads were trimmed with Trimmomatic, using default settings. None of the reads mapped to a HIV-1 subtype C reference genome or a consensus genome sequence of DRC66 sample generated by PCR, cloning and Sanger sequencing as detailed above.

Trimmed reads were further *de novo* assembled using Trinity v2.3.2 (5), retaining contigs of minimum 200 nt length. These contigs were compared with BLASTn to *Homo sapiens* reference genome (GRCh38.p10) as well as to a database compiled for IMSA+A (6) containing bacterial, viral and fungal genomes. A maximum of 200 hits with a minimum e-value of $1e^{-15}$ were retained. Contigs with no hits to these databases were subsequently compared to NCBI's nucleotide database using the same BLASTn procedure.

BLASTn outputs were further processed in R v3.4. For each contig, only BLASTn hits with the highest bit score were retained, with possible ties.

If a contig blasted to different taxa (which were gathered with taxize package (7)) with equal bit score, the lowest taxonomic level (e.g. genus, family, order etc) that the different taxa had in common was retained. To avoid false positive BLAST hits, we removed sequences for which the identified taxon level was only present once in the dataset in case the identified taxon level was higher than the species level and the alignment was shorter than 200 bases. The number of reads contributing to each contig were numerated by Trinity.

#### Finding similar sequences to DRC66

We ran partial DRC66 sequences of window sizes ranging from 200-1000 nt through NCBI BLAST and searched extensively for both partial and full genomic sequences deposited in GenBank that had close similarity to the DRC66 genomic sequence. To ensure we did not miss any partial HIV-1 gene sequence that would be phylogenetically closely related to DRC66 but would have a relatively low percentage nucleotide similarity to DRC66 sequence, we downloaded all sequences (>200 nt) marked as subtype C, as unknown subtype (U) and as inter-subtype recombinants from the Los Alamos National Laboratories (LANL) HIV database. For 14 sequence stretches of DRC66 that were largely free of unknown nucleotide sites and that ranged from 231 to 548 nt in length, we built alignments with corresponding LANL database sequences that overlapped for at least 150 nt, using the LANL codon-based HIVAlign tool (8) and choosing the Hidden Markov Model (9). Alignments contained between 830 and 5721 sequences. We inferred neighbor-joining trees in Geneious on the full alignments. Sequences that clustered with DRC66 were extracted together with reference subtype C sequences and were more closely examined for their relationship with DRC66 by building trees with RAxML v8.2 using the GTR-CAT model for rate heterogeneity, and a rapid bootstrap analysis with 200 replicates.

We found that the subtype-C like portions of the genomes of three circulating recombinant forms (designated CRF93_CPX) from Kinshasa and Mbuji-Mayi, DRC, sampled in 2008 (10), formed a monophyletic group with DRC66 in a maximum likelihood tree of a partial *pol* alignment (see also Figure 1C). Villabona *et al*. (10) and Rodgers *et al*. (11) describe two more "divergent C lineages" from recombinant genomes, CRF92_CU and single CU recombinant genome that clusters with an available partial *pol* sequence. We therefore built an alignment of the *pol* region sequences between 2485-4274 (HXB2 notation), for all these divergent C sequences together with the subsampled dataset A (see below). We verified in RDP4 (12) that there had been no recombination among lineages within this alignment.

#### Datasets compiled for full-genome phylogenetic analyses

We downloaded all non-recombinant near-complete (>7000 nt) HIV-1 group M genomes, that were sampled in Africa, from the LANL database on November 19 2018 (N=2342). We excluded subtype G genomes as subtype G originated as a recombinant between subtypes A and J and a putative subtype G parent (13). To maximize deep temporal coverage, we also downloaded all HIV-1 group M genomes from other parts of the world that were sampled before 1985 (N=78).

We filtered those sequences for which the year of sampling was not reported and randomly selected one sequence per patient. The REGA HIV subtyping tool v3.41 further identified seven significant inter-subtype recombinants in this dataset, which we removed. Fourteen hypermutated sequences identified by the LANL Quality Control tool (https://www.hiv.lanl.gov/content/sequence/QC/index.html), specifically Hypermut (14), were further removed. The final dataset contained 830 sequences, which was codon-aligned based on the translated gene sequences using the LANL HIVAlign tool (8) using the Hidden Markov Model (9). Alignments were manually checked and edited in Geneious v10.2.6.

Intra-subtype recombination was investigated for sequences from each subtype separately in RDP4 (12). Sequence regions larger than 300 nt identified by at least six methods as involved in recombination were masked from the alignment.

This alignment was further down-sampled in five distinct sub-alignments, for the purpose of efficient computation, as well as to assess the inherent uncertainty through phylogenetic variation among random samples of actual HIV-1 diversity. For those HIV-1 genomes sampled after 1990, a random sample of 150 genomes was drawn five times without replacement. Because of the scarcity of non-subtype B genomes sampled before 1990, all of these were retained in each subsampled dataset. For subtype B genomes sampled before 1990, seven or six were randomly sampled without replacement. The first four subsampled datasets (A-D) contained 177 HIV-1 genomes, the fifth (E) 176 HIV-1 genomes.

#### Exploring time and root-to-tip distance correlations

Maximum-likelihood trees of the complete dataset and subsampled datasets (named A to E) were estimated with RAxML v8.2 using the GTR-CAT substitution model that accommodates among-site rate heterogeneity. In R 3.4.2, trees were midpoint rooted with phytools package (15) and the patristic distance between root to all tips determined with adephylo package (16). Root-to-tip distance was plotted and regressed with time of tip sampling with R's stats package.

#### Divergence time dating in BEAST

Tip-date calibrated phylogenetic trees were estimated using BEAST v. 10.4 (17) for each subsampled dataset A-E. We used a GTR substitution model with 6 categories in a gamma rate heterogeneity model, and an uncorrelated relaxed clock to model evolutionary rates along the tree (18). We used the skygrid coalescent model with 50 time slots going back 120 years as a tree prior (19). Three independent Markov chain Monte Carlo (MCMC) chains were run for ~300,000,000 generations each logging every 10000 generations. All BEAST xml files are available in a supplementary zip folder. Convergence and ESS values of all parameters were evaluated in Tracer v1.6. The first 20% of the sampled MCMC runs were removed as burn-in and the three parameter log and trees log output files for each dataset were combined in R v3.4.2 and Logcombiner (17).

These analyses were repeated but with 1) the DRC66 sequence removed from the alignment; and 2) without providing the sampling year of DRC66 but instead having its sampling date estimated from the rest of data using BEAST's "tip date sampling" option and a uniform prior bound between 0 and 1000 (in years since youngest sampling date) (20). For one of the subsampled datasets (B) when using the model with DRC66's tip-date sampling option, convergence of the three MCMC chains for node height of the subtype C +

DRC66 clade was not readily reached, so we re-ran BEAST for this dataset but enforcing the monophyly of that clade.

To explore how well tip dates of other samples could be estimated, we repeated the BEAST analyses of one of the subsampled alignments (A) for each of five different sequences using BEAST's "tip date sampling" option. Accession numbers of these tip-date estimated sequences were M62320 (a subtype A from 1985 Uganda), KJ704795 (a subtype B from 1978 USA), DQ369980 (a subtype C from 2003 South Africa), and AF286236 (an unknown subtype from 1983 DRC). For the latter, representing a divergent singleton lineage in this dataset, the position in the phylogeny was not well determined without the information with its tip date. Based on RAxML analyses and BEAST analyses in which the tip date was included, AF286236 should form a monophyletic group with the subtype C clade – so we subsequently enforced this monophyly in the BEAST analyses.

##### Signatures of drug resistance

We examined whether the DRC66 sample could be resistant to drugs currently used to treat HIV infection using the HIV Drug Resistance Database Program (21), which specifically searches for amino acid signatures of inhibitor-drug resistance in *pol* sequences coding for Integrase, Protease and Reverse transcriptase.

#### Supplementary Results

##### Results Illumina sequencing after de novo assembly

We *de novo* assembled about 20% of the reads in contigs >200 nt using Trinity. Preliminary scans revealed that shorter contigs mostly lead to unreliable organism identification, even at the family level or higher; therefore the set of contigs >200nt were used for an approximation of the relative contribution of different groups of organisms.

Identification of a fungus-borne virus (related to the known yeast virus *Saccharomyces cerevisiae* virus L-A) provides evidence that our deep sequencing approach was able to detect RNA viruses, although this virus contained a double-stranded RNA genome instead of the less stable ssRNA genomes of lentiviruses. The largest group of reads in the *de novo* assembled contig group were of fungal origin (46%), followed by bacteria (35%) (Table S2). Most of these fungi and bacteria were likely environmental organisms that we speculate entered the FFPE ganglion tissue sample during sample preparation in 1966 or during its long storage time since. Human reads only constituted 0.04% of the total Illumina reads, implying either that our duplex-specific nuclease digestion procedure was highly effective, or that the relative contribution of RNA from environmental organisms that invaded during or after the formalin fixation process was so high that they swamped RNA intrinsic to the sample. Nevertheless, organisms that often represent opportunistic infections in AIDS patients were also detected, such as *Bartonella*, *Mycoplasma* and *Candida* (Table S2), though it cannot be excluded that these were derived from later-invading environmental variants of these genera rather than being present in the patient's original biopsy material.

**Table S2.** Taxonomy results of Illumina sequencing, see methods in main text. Identified genera are organized by ecological and higher taxonomical properties. Number of de novo assembled contigs (>200nt) belonging to that genus and the total number of reads from which these contigs were assembled are given.

For those contigs for which it was not possible to assign a genus level, contig and read counts for either families or phyla are given.

| | Number of contigs | Number of reads |
|---|---|---|
| **Determinable at genus level** | **total: 2014** | **total: 4774189** |
| **Common lab DNA/RNA** | **total: 32** | **total: 1212861** |
| *Corynebacterium* | 3 | 57238 |
| *Micrococcus* | 2 | 45280 |
| *Microlunatus* | 1 | 8621 |
| *Ochrobactrum* | 19 | 406194 |
| *Pedobacter* | 1 | 11188 |
| *Phix174microvirus* | 2 | 647311 |
| *Sphingobium* | 2 | 24657 |
| *Streptococcus* | 2 | 12372 |
| **Environmental arthropod** | **total: 3** | **total: 140257** |
| *Suidasia* | 3 | 140257 |
| **Environmental bacteria** | **total: 8** | **total: 127366** |
| *Bdellovibrio* | 1 | 4710 |
| *Cyclobacterium* | 1 | 15494 |
| *Halanaerobium* | 1 | 15604 |
| *Prochlorococcus* | 1 | 26904 |
| *Sphingomonas* | 4 | 64654 |
| **Environmental fungi / fungi-like** | **total: 109** | **total: 2124908** |
| *Allomyces* | 2 | 26738 |
| *Aspergillus* | 80 | 1368943 |
| *Batrachochytrium* | 2 | 54830 |
| *Coprinopsis* | 1 | 31009 |
| *Debaryomyces* | 3 | 34123 |
| *Hyaloperonospora* | 1 | 17894 |
| *Mucor* | 1 | 24294 |
| *Naumovozyma* | 2 | 19843 |
| *Spizellomyces* | 3 | 73575 |
| *Talaromyces* | 10 | 422886 |
| *Torulaspora* | 1 | 33169 |
| *Wallemia* | 3 | 17604 |
| **Environmental fungus virus** | **total: 1** | **total: 43590** |
| *Totivirus* | 1 | 43590 |
| **Human** | **total: 21** | **total: 241550** |
| *Homo* | 21 | 241550 |
| **Potential normal human microbiome** | **total: 10** | **total: 296514** |
| *Cutibacterium* | 1 | 13034 |
| *Escherichia* | 6 | 244140 |
| *Malassezia* | 2 | 21342 |
| *Pseudopropionibacterium* | 1 | 17998 |
| **Potential AIDS-associated bacteria** | **total: 13** | **total: 528218** |
| *Bartonella* | 8 | 403212 |
| *Brucella* | 1 | 40245 |
| *Mycoplasma* | 1 | 714 |
| *Salmonella* | 2 | 83396 |
| *Veillonella* | 1 | 651 |
| **Potential AIDS-associated fungi** | **total: 7** | **total: 58925** |
| *Candida* | 4 | 23927 |
| *Coccidioides* | 2 | 30262 |
| *Nakaseomyces* | 1 | 4736 |

| | | |
|---|---|---|
| **Only determinable at family level** | **total: 13** | **total: 393336** |
| Brucellaceae (Bacteria) | 2 | 77666 |
| Enterobacteriaceae (Bacteria) | 6 | 167762 |
| Saccharomycetaceae (Fungi) | 1 | 1053 |
| Sclerotiniaceae (Fungi) | 2 | 140093 |
| Sordariaceae (Fungi) | 2 | 6762 |
| **Only determinable at phylum level** | **total: 40** | **total: 727600** |
| Actinobacteria (Bacteria) | 1 | 9466 |
| Apicomplexa (Protozoa) | 2 | 29739 |
| Arthropoda | 1 | 32760 |
| Ascomycota (Fungi) | 15 | 321816 |
| Bacteroidetes (Bacteria) | 2 | 27376 |
| Basidiomycota (Fungi) | 3 | 23932 |
| Chordata | 3 | 6011 |
| Cyanobacteria (Bacteria) | 1 | 1698 |
| Firmicutes (Bacteria) | 3 | 47826 |
| Mucoromycota (Fungi) | 1 | 7368 |
| Proteobacteria (Bacteria) | 8 | 219608 |
| **Taxonomy not determinable** | **total: 5** | **total: 76668** |
| **Grand Total** | **262** | **5971793** |

**Table S3.** From the time-scaled BEAST phylogenies: estimated mean evolutionary rates, estimated dates of the root and two internal nodes, estimated dates of various single tips whose dates were unspecified and left to be estimated.

| Dataset | Mean evolutionary rate across tree | Root date | Conventional subtype C clade date | Conventional C + DRC66 clade date | Date of single, to be estimated tip (specified in first column) |
|---|---|---|---|---|---|
| Subsample A | 0.00344 (0.00296-0.00394) | 1902 (1887-1916) | 1941 (1932-1950) | 1923 (1911-1935) | NA |
| Subsample B | 0.00306 (0.00262-0.00352) | 1896 (1881-1913) | 1944 (1935-1952) | 1921 (1907-1934) | NA |
| Subsample C | 0.00326 (0.00284-0.00365) | 1903 (1890-1916) | 1941 (1932-1949) | 1927 (1916-1937) | NA |
| Subsample D | 0.003 (0.00258-0.00346) | 1900 (1885-1915) | 1938 (1928-1948) | 1921 (1907-1935) | NA |
| Subsample E | 0.00327 (0.00281-0.00372) | 1905 (1890-1918) | 1941 (1932-1950) | 1927 (1915-1938) | NA |
| Subsample A - DRC66 sequence not included | 0.00332 (0.00289-0.00377) | 1899 (1884-1913) | 1937 (1927-1946) | NA | NA |
| Subsample B - DRC66 sequence not included | 0.00299 (0.00255-0.00345) | 1895 (1878-1912) | 1942 (1933-1951) | NA | NA |
| Subsample C - DRC66 sequence not included | 0.00316 (0.00277-0.00355) | 1901 (1887-1914) | 1939 (1929-1947) | NA | NA |
| Subsample D - DRC66 sequence not included | 0.00292 (0.0025-0.00334) | 1898 (1883-1913) | 1936 (1926-1946) | NA | NA |
| Subsample E - DRC66 sequence not included | 0.00313 (0.00271-0.00355) | 1902 (1888-1888) | 1938 (1928-1928) | NA | NA |
| Subsample A - estimating DRC66 tip date | 0.00331 (0.00286-0.00377) | 1896 (1878-1913) | 1938 (1928-1947) | 1914 (1896-1932) | 1948 (1928-1971) |
| Subsample B - estimating DRC66 tip date | 0.00296 (0.00255-0.00338) | 1890 (1872-1907) | 1942 (1933-1950) | 1911 (1889-1928) | 1946 (1924-1966) |
| Subsample C - estimating DRC66 tip date | 0.00317 (0.00277-0.00358) | 1899 (1885-1914) | 1938 (1927-1947) | 1921 (1905-1934) | 1953 (1935-1971) |
| Subsample D - estimating DRC66 tip date | 0.0029 (0.00248-0.00334) | 1894 (1877-1911) | 1935 (1924-1946) | 1911 (1892-1930) | 1945 (1926-1966) |
| Subsample E - estimating DRC66 tip date | 0.00313 (0.00269-0.00357) | 1899 (1884-1915) | 1937 (1926-1947) | 1920 (1905-1935) | 1952 (1935-1969) |
| Subsample A - estimating A1\|1985\|UGA\|M62320 tip date | 0.00345 (0.00297-0.00399) | 1902 (1888-1918) | 1941 (1932-1950) | 1923 (1910-1936) | 1992 (1976-2010) |
| Subsample A - estimating B\|1978\|USA\|KJ704795 tip date | 0.00328 (0.00283-0.00377) | 1897 (1883-1913) | 1938 (1928-1948) | 1920 (1907-1933) | 1973 (1969-1978) |
| Subsample A - estimating C\|2003\|ZAF\|DQ369980 tip date | 0.00341 (0.00295-0.00391) | 1901 (1887-1915) | 1940 (1931-1949) | 1922 (1910-1934) | 1995 (1978-2014) |
| Subsample A - estimating D\|1983\|COD\|M27323 tip date | 0.00343 (0.00296-0.00397) | 1901 (1886-1916) | 1941 (1932-1950) | 1923 (1910-1936) | 1974 (1964-1985) |
| Subsample A - estimating U\|1983\|COD\|AF286236 tip date | 0.00345 (0.00298-0.00393) | 1900 (1887-1915) | 1941 (1932-1950) | 1922 (1909-1934) | 1954 (1933-1980) |

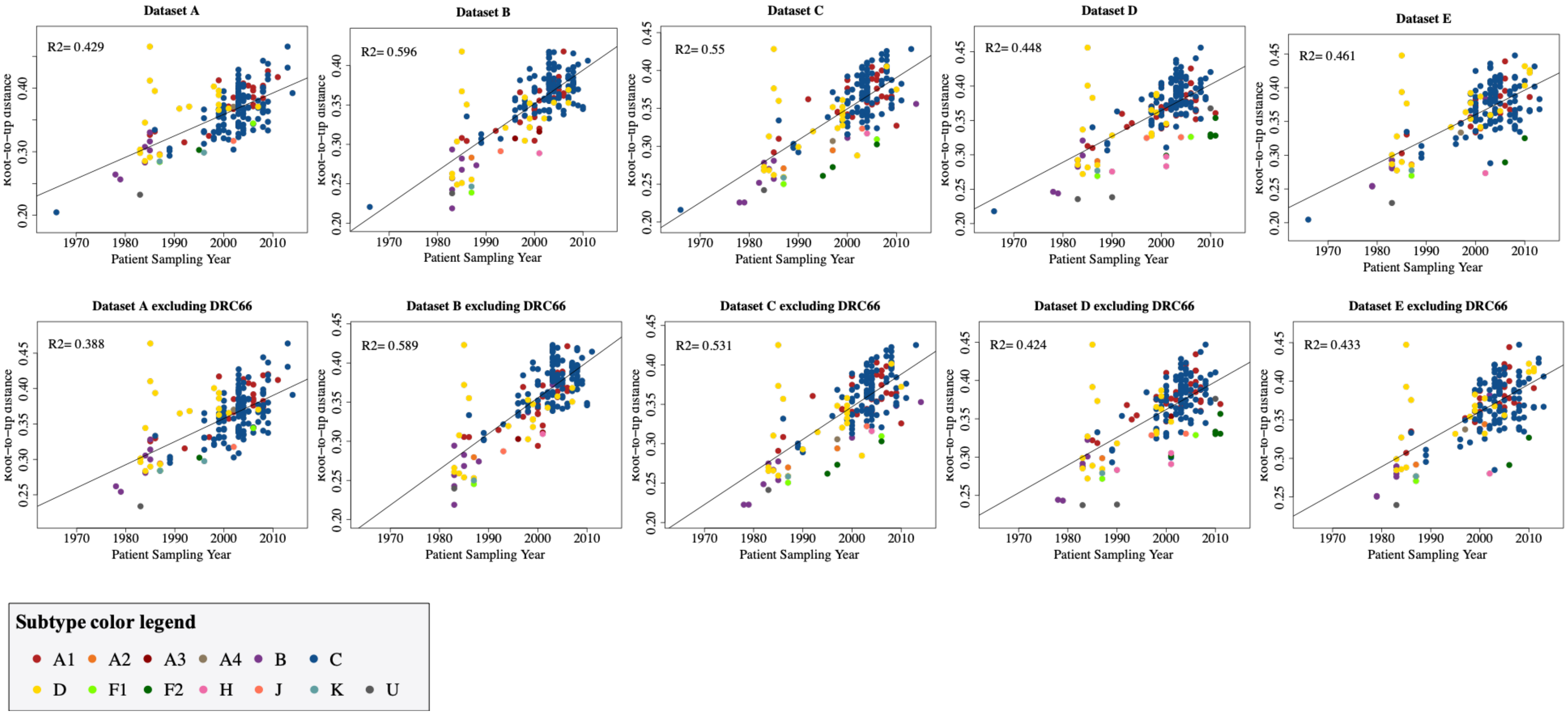

**Fig. S1.** Root-to-tip plots of ten maximum likelihood trees constructed in RAxML v8.2: five trees based on subsampled datasets A-E that included the DRC66 sequence, and five trees based on those sample subsampled datasets A-E but from which DRC66 was excluded.

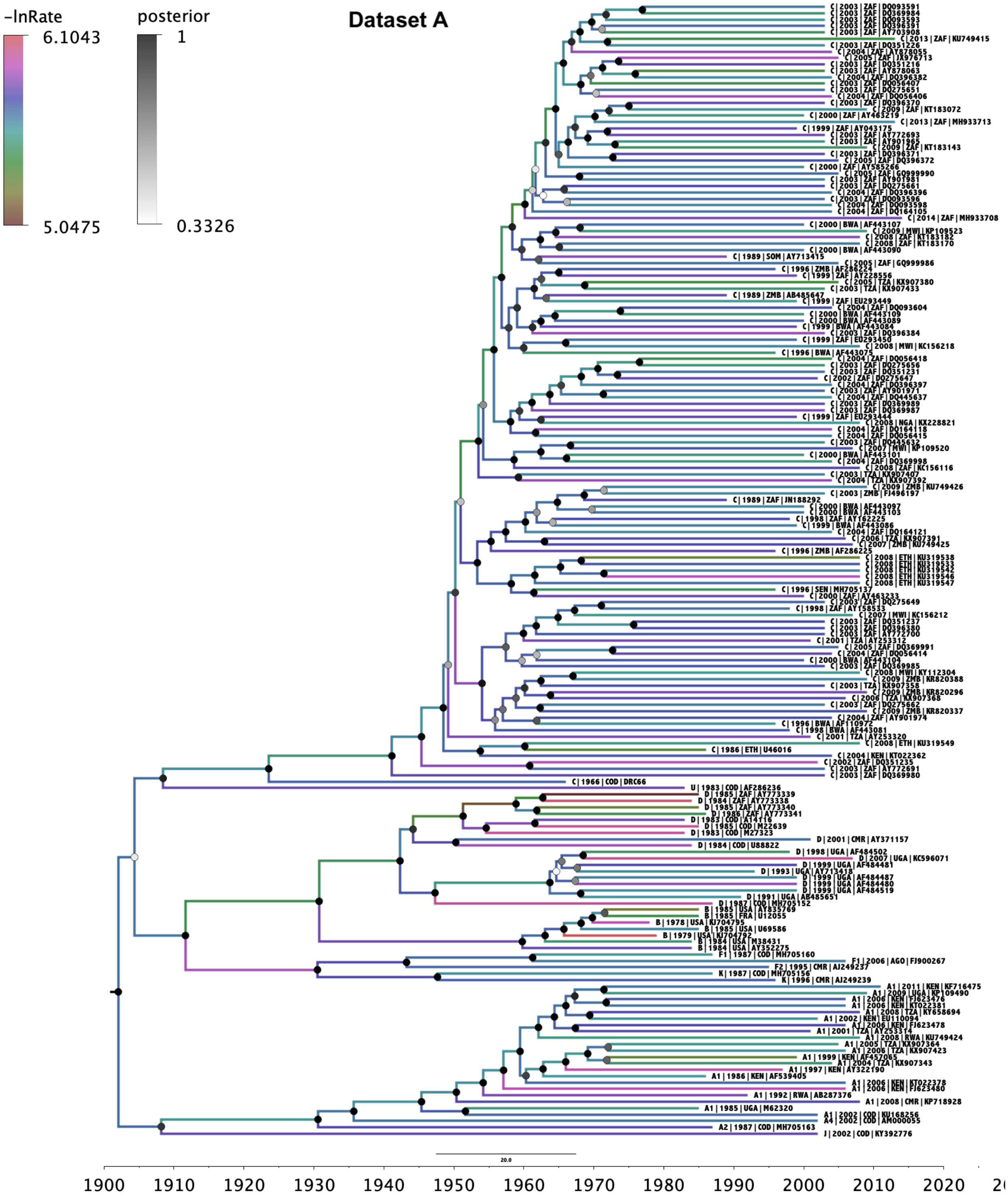
Dataset A
-lnRate
6.1043
5.0475
posterior
1
0.3326
20.0
1900
1910
1920
1930
1940
1950
1960
1970
1980
1990
2000
2010
2020

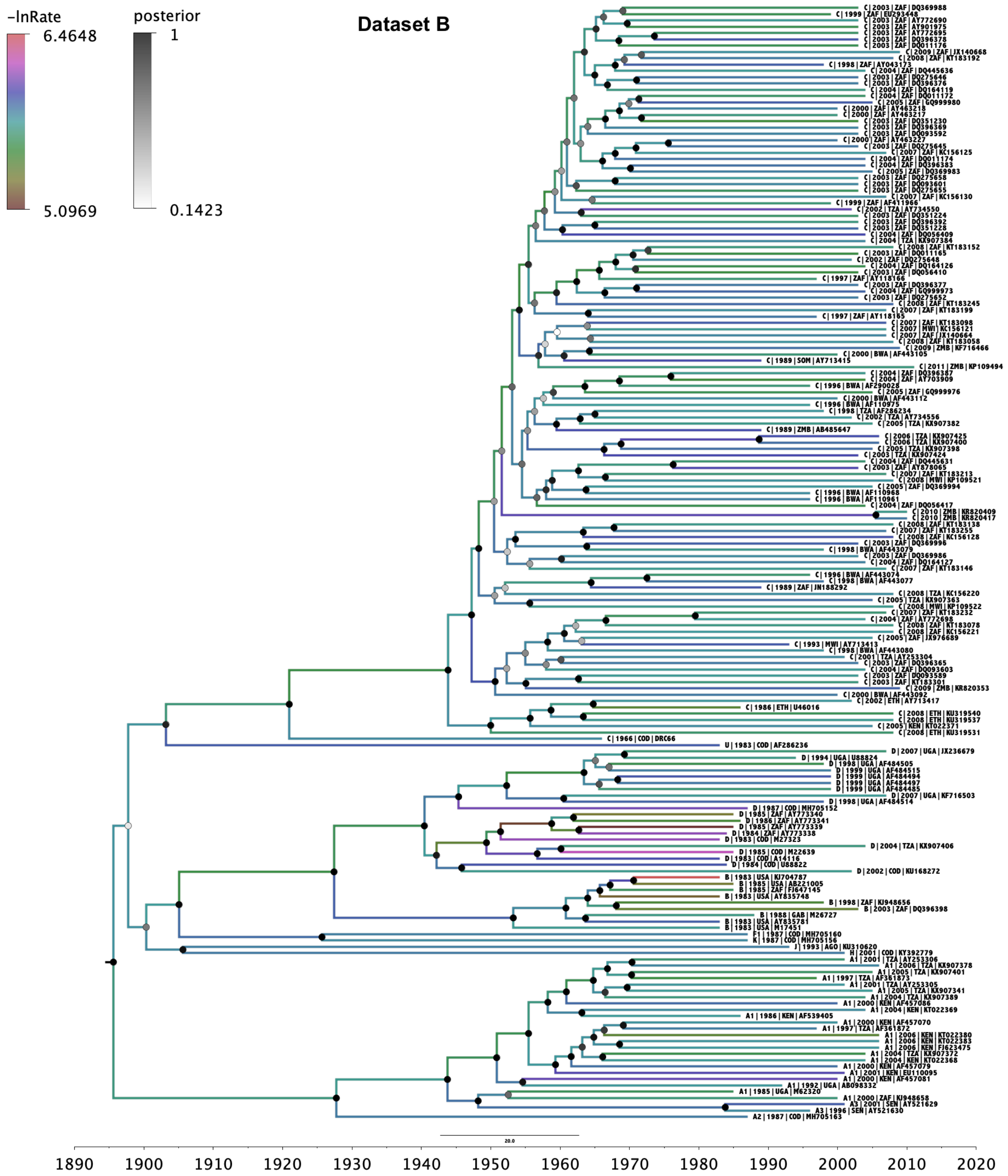
-lnRate
6.4648
5.0969
posterior
1
0.1423
Dataset B
1890
1900
1910
1920
1930
1940
1950
1960
1970
1980
1990
2000
2010
2020

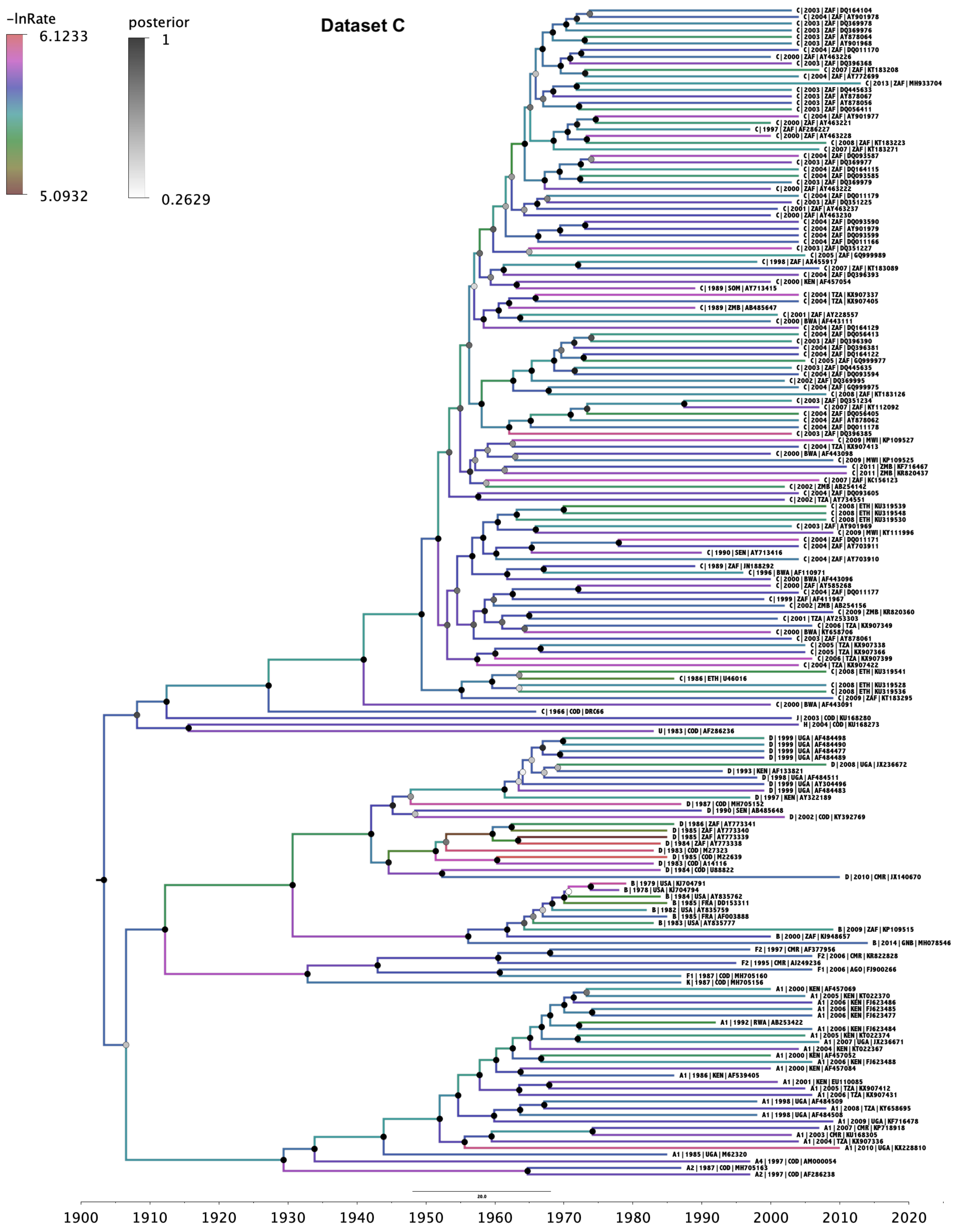
Dataset C
-lnRate
6.1233
5.0932
posterior
1
0.2629
20.0
1900
1910
1920
1930
1940
1950
1960
1970
1980
1990
2000
2010
2020

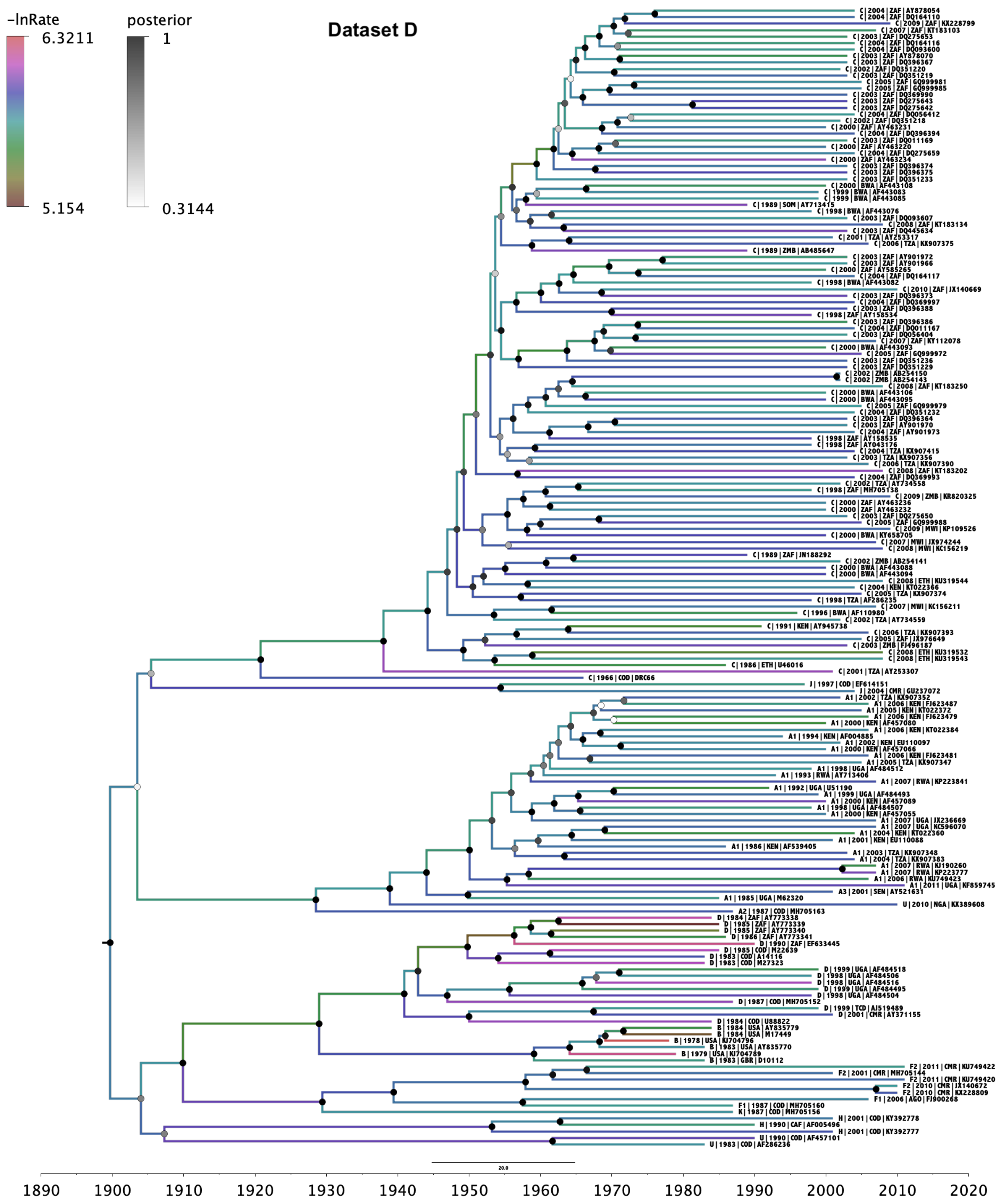
-lnRate
6.3211
5.154
posterior
1
0.3144
Dataset D
20.0
1890
1900
1910
1920
1930
1940
1950
1960
1970
1980
1990
2000
2010
2020

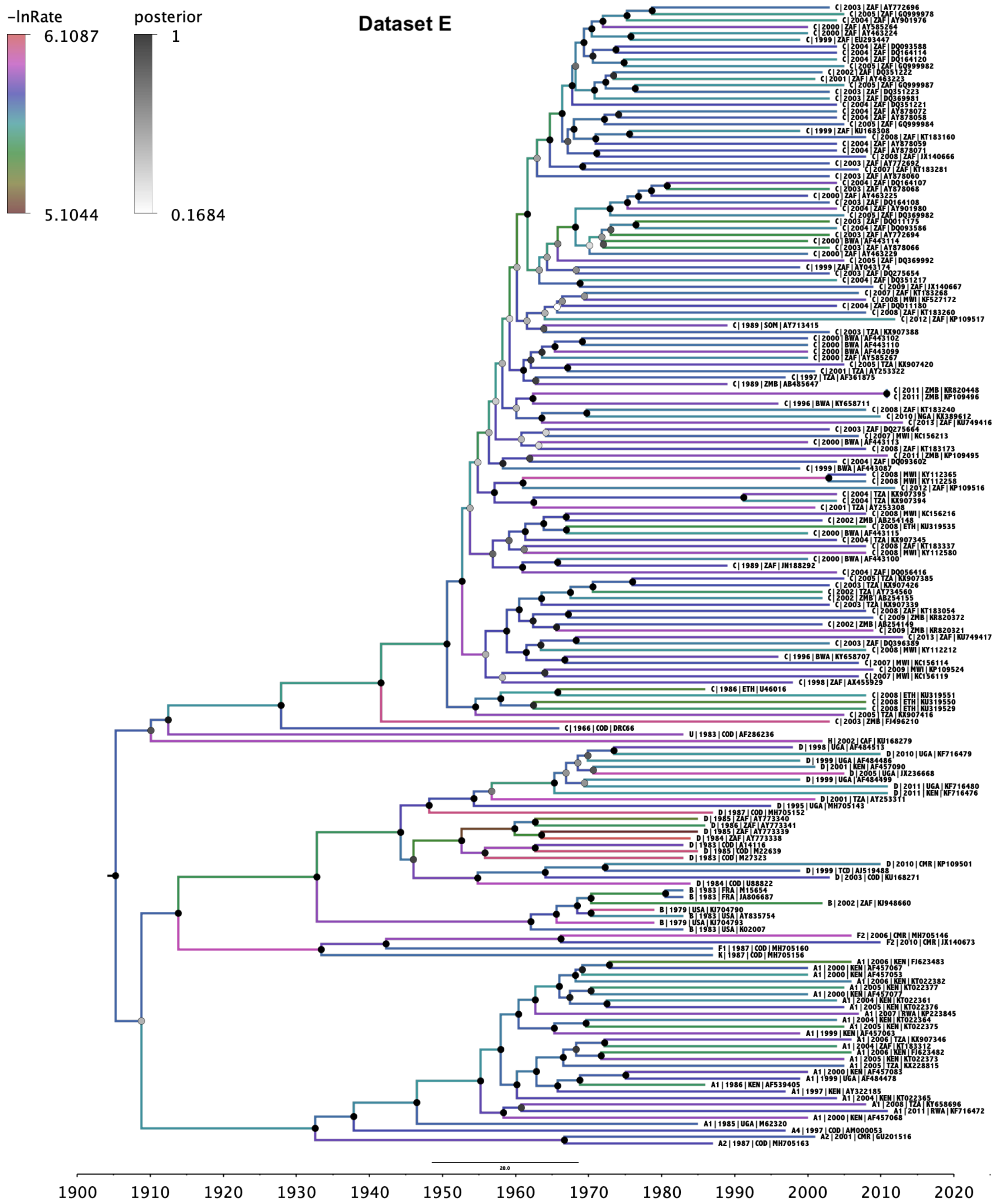

**Fig. S2.** Time scaled phylogenetic BEAST trees of subsampled datasets A-E, where sampling date of DRC66 was included in model. Branches are color coded by the log of the estimated evolutionary rate for that branch, drawn from a lognormal distribution using the uncorrelated relaxed clock model (18). Node labels are coded on a grey scale by the posterior probability of monophyly of the corresponding clade. Subtype, year, country code of sampling and GenBank accession number, are given at each tip label (for DRC66 sample it states DRC66 instead of accession number).

**Table S4**. List of all primers used in jackhammer RT-PCRs for the genome characterization of DRC66.

| Primername | Primer sequence (5'-3') | Target | Purpose | Output |
|---|---|---|---|---|
| HE10F | CAATATCASCACAARCA | env | First sweep of PCRs general for HIV-1 group M | Did not yield sequence data |
| HE10R | GGTACTAYATCAAGTYT | env | First sweep of PCRs general for HIV-1 group M | Did not yield sequence data |
| HE11F | TGCAYTTTTTCGTARAC | env | First sweep of PCRs general for HIV-1 group M | Did not yield sequence data |
| HE11R | CTGTGTAATGACTGAKSTGT | env | First sweep of PCRs general for HIV-1 group M | Did not yield sequence data |
| HE12F | ACASATCAGTCMTTACACAG | env | First sweep of PCRs general for HIV-1 group M | Did not yield sequence data |
| HE12R | AAACCAGCCGGGGYAC | env | First sweep of PCRs general for HIV-1 group M | Did not yield sequence data |
| HE13F | CATTTGAGCCAATTCCCAT | env | First sweep of PCRs general for HIV-1 group M | Did not yield sequence data |
| HE13R | GTCCTKTTCCATTGAACKT | env | First sweep of PCRs general for HIV-1 group M | Did not yield sequence data |
| HE14F | GGTTTTGCRATTCTAAARTG | env | First sweep of PCRs general for HIV-1 group M | Yielded sequence data |
| HE14R | CATGTGTACATTGTACTGTG | env | First sweep of PCRs general for HIV-1 group M | Yielded sequence data |
| HE15F | AAGGACCATGYAMAAATGTC | env | First sweep of PCRs general for HIV-1 group M | Yielded sequence data |
| HE15R | CAGCARTTGAGTTGAYAC | env | First sweep of PCRs general for HIV-1 group M | Yielded sequence data |
| HE19F | TTCWCGRACAATGCT | env | First sweep of PCRs general for HIV-1 group M | Did not yield sequence data |
| HE19R | GTTGTTGGGTCTTRTAC | env | First sweep of PCRs general for HIV-1 group M | Did not yield sequence data |
| HE1F | GTGCWTCAGATGCTARAGC | env | First sweep of PCRs general for HIV-1 group M | Did not yield sequence data |
| HE1R | GGGGTCTGTGGGTACAC | env | First sweep of PCRs general for HIV-1 group M | Did not yield sequence data |
| HE20F | CAGCTGAACACATCTGT | env | First sweep of PCRs general for HIV-1 group M | Did not yield sequence data |
| HE20R | ATGCTCTCCCTGGYCCT | env | First sweep of PCRs general for HIV-1 group M | Did not yield sequence data |
| HE21F | AGACCCAACAACAATACAAG | env | First sweep of PCRs general for HIV-1 group M | Yielded sequence data |
| HE21R | CTCCTATTATKTCTCCTGTTG | env | First sweep of PCRs general for HIV-1 group M | Yielded sequence data |
| HE22F | ATATAGGRCCAGGGAGAG | env | First sweep of PCRs general for HIV-1 group M | Did not yield sequence data |
| HE22R | GTTACAATRTGCTTKTC | env | First sweep of PCRs general for HIV-1 group M | Did not yield sequence data |
| HE23F | RCAACAGGAGAMATAATAGG | env | First sweep of PCRs general for HIV-1 group M | Did not yield sequence data |
| HE23R | GTGTYATTCCATTYTGC | env | First sweep of PCRs general for HIV-1 group M | Did not yield sequence data |
| HE24F | GTACAGCARAATGGAATRAC | env | First sweep of PCRs general for HIV-1 group M | Did not yield sequence data |
| HE24R | CCTGAGGAKTGATTAAAGAYT | env | First sweep of PCRs general for HIV-1 group M | Did not yield sequence data |
| HE25F | GTGGAGGRGAATTTTTCTAC | env | First sweep of PCRs general for HIV-1 group M | Did not yield sequence data |
| HE25R | GACCCTTCARYATTCCAAG | env | First sweep of PCRs general for HIV-1 group M | Did not yield sequence data |
| HE26F | TTGGAGTRYTGAAGGGTCA | env | First sweep of PCRs general for HIV-1 group M | Did not yield sequence data |
| HE26R | TCCTACTTYCTGCCACMTG | env | First sweep of PCRs general for HIV-1 group M | Did not yield sequence data |
| HE27F | CACMCTCCCATGYAGA | env | First sweep of PCRs general for HIV-1 group M | Did not yield sequence data |
| HE27R | GAGGRGCATACATTGCTT | env | First sweep of PCRs general for HIV-1 group M | Did not yield sequence data |
| HE28F | GTGGCAGAAAGTAGGAA | env | First sweep of PCRs general for HIV-1 group M | Did not yield sequence data |
| HE28R | GTTAATAGYAKCCCTGT | env | First sweep of PCRs general for HIV-1 group M | Did not yield sequence data |
| HE29F | ATGTATGCYCCTCCCATC | env | First sweep of PCRs general for HIV-1 group M | Did not yield sequence data |
| HE29R | GTTGCTATYACCACCATCYC | env | First sweep of PCRs general for HIV-1 group M | Did not yield sequence data |
| HE2F | CCTGTGTACCCACAGAC | env | First sweep of PCRs general for HIV-1 group M | Did not yield sequence data |
| HE2R | CATGCATCTGKTCTACCATG | env | First sweep of PCRs general for HIV-1 group M | Did not yield sequence data |
| HE30F | CAGGGMTRCTATTAACAAG | env | First sweep of PCRs general for HIV-1 group M | Did not yield sequence data |
| HE30R | CTTCTCCAATTGTCCMTCA | env | First sweep of PCRs general for HIV-1 group M | Did not yield sequence data |
| HE31F | AGACCTGGAGGAGGARA | env | First sweep of PCRs general for HIV-1 group M | Did not yield sequence data |
| HE31R | GGTGCTACYCCTAWTGGT | env | First sweep of PCRs general for HIV-1 group M | Did not yield sequence data |
| HE32F | KGGACAATTGGAGAAGTGA | env | First sweep of PCRs general for HIV-1 group M | Did not yield sequence data |
| HE32R | TCTGCACCACTCTTCTCT | env | First sweep of PCRs general for HIV-1 group M | Did not yield sequence data |
| HE33F | GCACCCACYARGGCAA | env | First sweep of PCRs general for HIV-1 group M | Did not yield sequence data |
| HE33R | TCCCAAGAMCCCAAGGAA | env | First sweep of PCRs general for HIV-1 group M | Did not yield sequence data |
| HE34F | GGTGCAGAGAGAAAAAAGAG | env | First sweep of PCRs general for HIV-1 group M | Yielded sequence data |
| HE34R | CTTCCTGCTGCTCCCAA | env | First sweep of PCRs general for HIV-1 group M | Yielded sequence data |
| HE35F | TAGGAGCTDTGTTCCTTGG | env | First sweep of PCRs general for HIV-1 group M | Did not yield sequence data |
| HE35R | CTGTACCGTCAGCGTYA | env | First sweep of PCRs general for HIV-1 group M | Did not yield sequence data |
| HE36F | CAGCAGGAAGCACYATGG | env | First sweep of PCRs general for HIV-1 group M | Yielded sequence data |
| HE36R | CTGYTGCACTATACCAGA | env | First sweep of PCRs general for HIV-1 group M | Yielded sequence data |
| HE37F | GACGGTACAGRCCAGA | env | First sweep of PCRs general for HIV-1 group M | Did not yield sequence data |
| HE37R | GCCTCAATAGCYCTCAG | env | First sweep of PCRs general for HIV-1 group M | Did not yield sequence data |
| HE38F | RTCTGGTATAGTGCARCA | env | First sweep of PCRs general for HIV-1 group M | Did not yield sequence data |
| HE38R | AGACTGTGAGTTGCAACAG | env | First sweep of PCRs general for HIV-1 group M | Did not yield sequence data |
| HE39F | CAATTTGCTGAGRGCTA | env | First sweep of PCRs general for HIV-1 group M | Did not yield sequence data |
| HE39R | GAYTCTTGCCTGGAGYT | env | First sweep of PCRs general for HIV-1 group M | Did not yield sequence data |
| HE3F | CAGACCCYARCCCACA | env | First sweep of PCRs general for HIV-1 group M | Yielded sequence data |
| HE3R | TCCTCATGCATYTGKTC | env | First sweep of PCRs general for HIV-1 group M | Yielded sequence data |
| HE40F | CTCACAGTCTGGGGCAT | env | First sweep of PCRs general for HIV-1 group M | Did not yield sequence data |
| HE40R | AAATYCCYAGGAGCTGT | env | First sweep of PCRs general for HIV-1 group M | Did not yield sequence data |
| HE41F | TGGCTGTGGAAAGATACCT | env | First sweep of PCRs general for HIV-1 group M | Did not yield sequence data |
| HE41R | GYAGTGGTGCARATGAGT | env | First sweep of PCRs general for HIV-1 group M | Did not yield sequence data |
| HE42F | AGCTCCTRGGRATTTGG | env | First sweep of PCRs general for HIV-1 group M | Did not yield sequence data |
| HE42R | CTCCAACTAGYATTCCAAG | env | First sweep of PCRs general for HIV-1 group M | Did not yield sequence data |
| HE43F | CTCATYTGCACCACTRCT | env | First sweep of PCRs general for HIV-1 group M | Did not yield sequence data |
| HE43R | TCCCACTSCATCCAGGT | env | First sweep of PCRs general for HIV-1 group M | Did not yield sequence data |
| HE44F | CCTTGGAATGCTAGTTGG | env | First sweep of PCRs general for HIV-1 group M | Did not yield sequence data |
| HE44R | TCTCTKTCCCACTSCATC | env | First sweep of PCRs general for HIV-1 group M | Yielded sequence data |
| HE45F | TTGGAATMACAYGACCT | env | First sweep of PCRs general for HIV-1 group M | Did not yield sequence data |
| HE45R | CTTGCTGGTTYTGYGAT | env | First sweep of PCRs general for HIV-1 group M | Did not yield sequence data |

| | | | | |
|---|---|---|---|---|
| HE46F | GGATGSAGTGGGAMAGA | env | First sweep of PCRs general for HIV-1 group M | Did not yield sequence data |
| HE46R | TCTTGYTGGTTYTGCGAT | env | First sweep of PCRs general for HIV-1 group M | Did not yield sequence data |
| HE47F | TGAASAATCGCARAACC | env | First sweep of PCRs general for HIV-1 group M | Did not yield sequence data |
| HE47R | CCAAYTCCACAAACTTG | env | First sweep of PCRs general for HIV-1 group M | Did not yield sequence data |
| HE48F | GATAARTGGGCAAGTTTGTG | env | First sweep of PCRs general for HIV-1 group M | Did not yield sequence data |
| HE48R | ACCTAYYAAGCCTCCTAC | env | First sweep of PCRs general for HIV-1 group M | Did not yield sequence data |
| HE49F | ACARAYTGGCTGTGGTA | env | First sweep of PCRs general for HIV-1 group M | Did not yield sequence data |
| HE49R | CCTGCCTAACTCTATTYAC | env | First sweep of PCRs general for HIV-1 group M | Did not yield sequence data |
| HE4F | GACATGGTAGAMCARATGC | env | First sweep of PCRs general for HIV-1 group M | Did not yield sequence data |
| HE4R | ACTRACACAGAGTGGGGTT | env | First sweep of PCRs general for HIV-1 group M | Did not yield sequence data |
| HE50F | TAGGAGGCTTRRTAGGTT | env | First sweep of PCRs general for HIV-1 group M | Did not yield sequence data |
| HE50R | AGGYGGGTCTGAAAYGA | env | First sweep of PCRs general for HIV-1 group M | Did not yield sequence data |
| HE51F | AGTTAGGCAGGGATAYTCAC | env | First sweep of PCRs general for HIV-1 group M | Did not yield sequence data |
| HE51R | KATTCCTTCGGGCCTGTC | env | First sweep of PCRs general for HIV-1 group M | Did not yield sequence data |
| HE52F | TATCRTTTCAGACCCRCCT | env | First sweep of PCRs general for HIV-1 group M | Did not yield sequence data |
| HE52R | CTGTCTCTCTCTCCACCT | env | First sweep of PCRs general for HIV-1 group M | Did not yield sequence data |
| HE53F | GACAGGCCCGAAGGAA | env | First sweep of PCRs general for HIV-1 group M | Did not yield sequence data |
| HE53R | TAAGTGCTAAGAATCCRTYC | env | First sweep of PCRs general for HIV-1 group M | Did not yield sequence data |
| HE54F | GAGAGACAGAKACAGATCC | env | First sweep of PCRs general for HIV-1 group M | Did not yield sequence data |
| HE54R | CGCAGRTCGWCCCAGA | env | First sweep of PCRs general for HIV-1 group M | Did not yield sequence data |
| HE55F | CGATTAGTGRAYGGATTCT | env | First sweep of PCRs general for HIV-1 group M | Did not yield sequence data |
| HE55R | AGCGGTGGTARCTGAAG | env | First sweep of PCRs general for HIV-1 group M | Did not yield sequence data |
| HE56F | GAYCTGCGGARCCTGT | env | First sweep of PCRs general for HIV-1 group M | Did not yield sequence data |
| HE56R | CCAGAAGTTCCACARTCCT | env | First sweep of PCRs general for HIV-1 group M | Did not yield sequence data |
| HE57F | CKCTTGAGAGACTTACTCT | env | First sweep of PCRs general for HIV-1 group M | Did not yield sequence data |
| HE57R | CCAATATTTGAGGRYTTCC | env | First sweep of PCRs general for HIV-1 group M | Did not yield sequence data |
| HE58F | GAGGAYTGTGGAACTTCTG | env | First sweep of PCRs general for HIV-1 group M | Did not yield sequence data |
| HE58R | TCTTTAGTTCCTGRMTCCA | env | First sweep of PCRs general for HIV-1 group M | Did not yield sequence data |
| HE59F | GCCCTCAAATATTGKTGGA | env | First sweep of PCRs general for HIV-1 group M | Did not yield sequence data |
| HE59R | GAGCAAGCTAAYAGCRCT | env | First sweep of PCRs general for HIV-1 group M | Did not yield sequence data |
| HE5F | AGCCTAAAGCCATGTGT | env | First sweep of PCRs general for HIV-1 group M | Did not yield sequence data |
| HE5R | TCCCGCTACTACTAKTGGT | env | First sweep of PCRs general for HIV-1 group M | Did not yield sequence data |
| HE60F | ATTGGAKYCAGGAGCTA | env | First sweep of PCRs general for HIV-1 group M | Did not yield sequence data |
| HE60R | TGTCCCCYCAGCTACTG | env | First sweep of PCRs general for HIV-1 group M | Did not yield sequence data |
| HE61F | GCTTGCTCRAYGCCACA | env | First sweep of PCRs general for HIV-1 group M | Did not yield sequence data |
| HE61R | TAGGTATGTGGAGAAHAGC | env | First sweep of PCRs general for HIV-1 group M | Did not yield sequence data |
| HE6F | TTAACCCCACTCTGTGT | env | First sweep of PCRs general for HIV-1 group M | Did not yield sequence data |
| HE6R | CCATTATCATTCTCCCGCTA | env | First sweep of PCRs general for HIV-1 group M | Did not yield sequence data |
| HE7F | CAMTAGTAGTAGCTGGGAGA | env | First sweep of PCRs general for HIV-1 group M | Did not yield sequence data |
| HE7R | CCTCTTATGYTTGTGSTGA | env | First sweep of PCRs general for HIV-1 group M | Did not yield sequence data |
| HE8F | GCGGGAGAATGATAATGGA | env | First sweep of PCRs general for HIV-1 group M | Did not yield sequence data |
| HE8R | TCTTTCTGCAYCTTAYC | env | First sweep of PCRs general for HIV-1 group M | Did not yield sequence data |
| HE9F | GCTCTTTCAATATCACCACA | env | First sweep of PCRs general for HIV-1 group M | Did not yield sequence data |
| HE9R | CAAGGTTACGAAAAARYGCA | env | First sweep of PCRs general for HIV-1 group M | Did not yield sequence data |
| HZ10F | GGAMACAGCAGYCAGA | gag | First sweep of PCRs general for HIV-1 group M | Did not yield sequence data |
| HZ10R | GATATKGCCTGATGTACCAT | gag | First sweep of PCRs general for HIV-1 group M | Did not yield sequence data |
| HZ11F | AGAACMTCCAGGGGCAA | gag | First sweep of PCRs general for HIV-1 group M | Did not yield sequence data |
| HZ11R | CTGAAAGCCTTYTCTTCTAC | gag | First sweep of PCRs general for HIV-1 group M | Did not yield sequence data |
| HZ12F | CATCAGGCMATATCACCTA | gag | First sweep of PCRs general for HIV-1 group M | Did not yield sequence data |
| HZ12R | GGTATYACTTCTGGGCTGA | gag | First sweep of PCRs general for HIV-1 group M | Did not yield sequence data |
| HZ13F | GTAGAAGARAAGGCTTTCAG | gag | First sweep of PCRs general for HIV-1 group M | Did not yield sequence data |
| HZ13R | TTGTGGGGTGGCTCCT | gag | First sweep of PCRs general for HIV-1 group M | Did not yield sequence data |
| HZ14F | GCCCAGAAGTRATACCCAT | gag | First sweep of PCRs general for HIV-1 group M | Yielded sequence data |
| HZ14R | CCCCCACTGTGTTTAGC | gag | First sweep of PCRs general for HIV-1 group M | Yielded sequence data |
| HZ16F | ACACAGTGGGGGGACA | gag | First sweep of PCRs general for HIV-1 group M | Yielded sequence data |
| HZ16R | CTGGATGCAMTCTATCCCAT | gag | First sweep of PCRs general for HIV-1 group M | Yielded sequence data |
| HZ17F | AGARACCATCAATGAGGAAG | gag | First sweep of PCRs general for HIV-1 group M | Yielded sequence data |
| HZ17R | TCTCTCATYTGGCCTGGT | gag | First sweep of PCRs general for HIV-1 group M | Yielded sequence data |
| HZ18F | GGATAGAKTGCATCCAGTG | gag | First sweep of PCRs general for HIV-1 group M | Did not yield sequence data |
| HZ18R | CTATGTCACTTCCCCTTGG | gag | First sweep of PCRs general for HIV-1 group M | Did not yield sequence data |
| HZ19F | GTGCATGCAGGGCCTA | gag | First sweep of PCRs general for HIV-1 group M | Yielded sequence data |
| HZ19R | TTGTTCCTGAAGGGTACTAG | gag | First sweep of PCRs general for HIV-1 group M | Yielded sequence data |
| HZ1f | GAGAGCGTCAGTATTAAGC | gag | First sweep of PCRs general for HIV-1 group M | Yielded sequence data |
| HZ1r | TTCTTTCCYCCTGGCCTT | gag | First sweep of PCRs general for HIV-1 group M | Yielded sequence data |
| HZ20F | GAACCAAGGGGAAGTGAC | gag | First sweep of PCRs general for HIV-1 group M | Yielded sequence data |
| HZ20R | CCTACTGGGATAGGTGGAT | gag | First sweep of PCRs general for HIV-1 group M | Yielded sequence data |
| HZ21F | GAAGTGACATAGCAGGAAC | gag | First sweep of PCRs general for HIV-1 group M | Yielded sequence data |
| HZ21R | ATTTCTCCTACTGGGATAGG | gag | First sweep of PCRs general for HIV-1 group M | Yielded sequence data |
| HZ22F | AATCCACCTATCCCAGT | gag | First sweep of PCRs general for HIV-1 group M | Did not yield sequence data |
| HZ22R | AATGCTGGWAGGRCTA | gag | First sweep of PCRs general for HIV-1 group M | Did not yield sequence data |
| HZ23F | ATGTATAGYCCTACCAGCA | gag | First sweep of PCRs general for HIV-1 group M | Did not yield sequence data |
| HZ23R | TAGAACCGRTCTACATAGTC | gag | First sweep of PCRs general for HIV-1 group M | Did not yield sequence data |
| HZ24F | GGACATAARACAAGGACCAA | gag | First sweep of PCRs general for HIV-1 group M | Yielded sequence data |
| HZ24R | CTTGYTCGGCTCTTAGAGT | gag | First sweep of PCRs general for HIV-1 group M | Yielded sequence data |
| HZ25F | RGAACCCTTTAGAGACT | gag | First sweep of PCRs general for HIV-1 group M | Did not yield sequence data |
| HZ25R | GGTTTCTGTCATCCA | gag | First sweep of PCRs general for HIV-1 group M | Did not yield sequence data |

| | | | | |
|---|---|---|---|---|
| HZ26F | RCAAGCTTCACAGGAKGT | gag | First sweep of PCRs general for HIV-1 group M | Did not yield sequence data |
| HZ26R | CCAATGCTTTYAARATAGTC | gag | First sweep of PCRs general for HIV-1 group M | Did not yield sequence data |
| HZ27F | GATGACAGAAACCTTGTTGG | gag | First sweep of PCRs general for HIV-1 group M | Yielded sequence data |
| HZ27R | CTTCTARTGTAGCYGCTGGT | gag | First sweep of PCRs general for HIV-1 group M | Yielded sequence data |
| HZ29F | RTTGGGACCAGCRGCT | gag | First sweep of PCRs general for HIV-1 group M | Did not yield sequence data |
| HZ29R | GCTTCMGCCAAAACTCTTG | gag | First sweep of PCRs general for HIV-1 group M | Did not yield sequence data |
| HZ30F | TGATGACAGCATGYCAG | gag | First sweep of PCRs general for HIV-1 group M | Did not yield sequence data |
| HZ30R | GTTACTTGGCTCATTGC | gag | First sweep of PCRs general for HIV-1 group M | Did not yield sequence data |
| HZ31F | CGGCCATAARGCAAGAGT | gag | First sweep of PCRs general for HIV-1 group M | Did not yield sequence data |
| HZ31R | CTARAATTGCCTYTCTGCA | gag | First sweep of PCRs general for HIV-1 group M | Did not yield sequence data |
| HZ33F | ATGCAGARAGGCAATTYTAG | gag | First sweep of PCRs general for HIV-1 group M | Did not yield sequence data |
| HZ33R | AGGGGCYCTGCAATTTYTG | gag | First sweep of PCRs general for HIV-1 group M | Did not yield sequence data |
| HZ34F | TGGCAARGAAGGGCACA | gag | First sweep of PCRs general for HIV-1 group M | Yielded sequence data |
| HZ34R | CCTTTCCACATTTCCAACAG | gag | First sweep of PCRs general for HIV-1 group M | Yielded sequence data |
| HZ35F | ATTGCAGRGCYCCTA | gag | First sweep of PCRs general for HIV-1 group M | Did not yield sequence data |
| HZ35R | TTAGCCTGTCTCTCA | gag | First sweep of PCRs general for HIV-1 group M | Did not yield sequence data |
| HZ36F | GGAAATGTGGAAAGGAAGG | gag | First sweep of PCRs general for HIV-1 group M | Did not yield sequence data |
| HZ36R | CTTGTRGGARGGCCAGA | gag | First sweep of PCRs general for HIV-1 group M | Did not yield sequence data |
| HZ37F | TGTACTGAGAGACAGGCTA | gag | First sweep of PCRs general for HIV-1 group M | Did not yield sequence data |
| HZ37R | GTTGGTTCTRGTCTGYTCTG | gag | First sweep of PCRs general for HIV-1 group M | Did not yield sequence data |
| HZ38F | TGGCCTTCCYACAAGG | gag | First sweep of PCRs general for HIV-1 group M | Did not yield sequence data |
| HZ38R | CCARACCTGAAGMTCTCTTC | gag | First sweep of PCRs general for HIV-1 group M | Did not yield sequence data |
| HZ39F | CCAGAGCAGACYAGARCCAA | gag | First sweep of PCRs general for HIV-1 group M | Did not yield sequence data |
| HZ39R | TCTGAGRGGGAGYTGTTG | gag | First sweep of PCRs general for HIV-1 group M | Did not yield sequence data |
| HZ40F | GAGCTTCAGGTYTGGGGWA | gag | First sweep of PCRs general for HIV-1 group M | Did not yield sequence data |
| HZ40R | GATCTGAGGGAAGYTAARGG | gag | First sweep of PCRs general for HIV-1 group M | Did not yield sequence data |
| HZ41F | ARCAGGAGCCGATAGAC | gag | First sweep of PCRs general for HIV-1 group M | Yielded sequence data |
| HZ41R | GAGGGGTCGYTGCCAA | gag | First sweep of PCRs general for HIV-1 group M | Yielded sequence data |
| HZ5F | CTGGCCTRTTAGAAACATC | gag | First sweep of PCRs general for HIV-1 group M | Yielded sequence data |
| HZ5R | GTTCTTCTGATCCTGTCTGA | gag | First sweep of PCRs general for HIV-1 group M | Yielded sequence data |
| HY10F | GTCTCCATAGAATGGAGGAA | vif-vpr-vpu | First sweep of PCRs general for HIV-1 group M | Did not yield sequence data |
| HY10R | CAGRTGAATTAGTYGGTCT | vif-vpr-vpu | First sweep of PCRs general for HIV-1 group M | Did not yield sequence data |
| HY11F | ACCCTGRCCTAGCAGA | vif-vpr-vpu | First sweep of PCRs general for HIV-1 group M | Yielded sequence data |
| HY11R | AGGGCTAACTMTAYGTCCTA | vif-vpr-vpu | First sweep of PCRs general for HIV-1 group M | Yielded sequence data |
| HY12F | ACCRACTAATTCATCTGYAT | vif-vpr-vpu | First sweep of PCRs general for HIV-1 group M | Did not yield sequence data |
| HY12R | TTCACAMCTAGGRCTAACTA | vif-vpr-vpu | First sweep of PCRs general for HIV-1 group M | Did not yield sequence data |
| HY13F | AGGACATAKAGTTAGYCCTA | vif-vpr-vpu | First sweep of PCRs general for HIV-1 group M | Yielded sequence data |
| HY13R | CCAAGTAYTGTAGAGATCCT | vif-vpr-vpu | First sweep of PCRs general for HIV-1 group M | Yielded sequence data |
| HY14F | CAAGCAGGACATAAYAAGG | vif-vpr-vpu | First sweep of PCRs general for HIV-1 group M | Yielded sequence data |
| HY14R | GTGGCTTTRTCYTTTTTGG | vif-vpr-vpu | First sweep of PCRs general for HIV-1 group M | Yielded sequence data |
| HY15F | CAAGGTAGGATCTCTACART | vif-vpr-vpu | First sweep of PCRs general for HIV-1 group M | Did not yield sequence data |
| HY15R | GTTTCCYAACACTRGGCAA | vif-vpr-vpu | First sweep of PCRs general for HIV-1 group M | Did not yield sequence data |
| HY16F | AAARGAYAAAGCCACCT | vif-vpr-vpu | First sweep of PCRs general for HIV-1 group M | Did not yield sequence data |
| HY16R | YTCCCTCTGTGGYCCT | vif-vpr-vpu | First sweep of PCRs general for HIV-1 group M | Did not yield sequence data |
| HY17F | AGAGGATAGATGGAACAAGC | vif-vpr-vpu | First sweep of PCRs general for HIV-1 group M | Yielded sequence data |
| HY17R | GCTTCRYTCTTAAGCTCCT | vif-vpr-vpu | First sweep of PCRs general for HIV-1 group M | Yielded sequence data |
| HY18F | CCACAGAGGGARCCAT | vif-vpr-vpu | First sweep of PCRs general for HIV-1 group M | Yielded sequence data |
| HY18R | GGAAARTGTCTRACAGCTTC | vif-vpr-vpu | First sweep of PCRs general for HIV-1 group M | Yielded sequence data |
| HY19F | RCCATACAATGAATGGACAC | vif-vpr-vpu | First sweep of PCRs general for HIV-1 group M | Did not yield sequence data |
| HY19R | GCCATGAAGCCATAYYCT | vif-vpr-vpu | First sweep of PCRs general for HIV-1 group M | Did not yield sequence data |
| HY20F | GGAGCTTAAGARYGAAGCT | vif-vpr-vpu | First sweep of PCRs general for HIV-1 group M | Did not yield sequence data |
| HY20R | CCTTCCCARGTATCYCCAT | vif-vpr-vpu | First sweep of PCRs general for HIV-1 group M | Did not yield sequence data |
| HY21F | GAGTATGGCTYCATRGCTT | vif-vpr-vpu | First sweep of PCRs general for HIV-1 group M | Did not yield sequence data |
| HY21R | GYATGGCYTCCACTCCT | vif-vpr-vpu | First sweep of PCRs general for HIV-1 group M | Did not yield sequence data |
| HY22F | CTTATGGAGATACYTGGGMA | vif-vpr-vpu | First sweep of PCRs general for HIV-1 group M | Did not yield sequence data |
| HY22R | GTCGRCACCCAATTCTGA | vif-vpr-vpu | First sweep of PCRs general for HIV-1 group M | Yielded sequence data |
| HY23F | GAGTGGAAGCCATRMTA | vif-vpr-vpu | First sweep of PCRs general for HIV-1 group M | Did not yield sequence data |
| HY23R | TATYCTGCTATGTCGRCAC | vif-vpr-vpu | First sweep of PCRs general for HIV-1 group M | Did not yield sequence data |
| HY24F | TYTRCAACAACTGCTGT | vif-vpr-vpu | First sweep of PCRs general for HIV-1 group M | Did not yield sequence data |
| HY24R | CTGTGGARTAATGCCTATYC | vif-vpr-vpu | First sweep of PCRs general for HIV-1 group M | Did not yield sequence data |
| HY25F | TCAGAATTGGGTGYCGA | vif-vpr-vpu | First sweep of PCRs general for HIV-1 group M | Did not yield sequence data |
| HY25R | CTAGGATCTACTGGMTCCAT | vif-vpr-vpu | First sweep of PCRs general for HIV-1 group M | Did not yield sequence data |
| HY26F | GGCATTATTCMACAGAGRAG | vif-vpr-vpu | First sweep of PCRs general for HIV-1 group M | Did not yield sequence data |
| HY26R | GACTYCCTGGATGCTTCC | vif-vpr-vpu | First sweep of PCRs general for HIV-1 group M | Did not yield sequence data |
| HY27F | GAAATGGAKCCAGTAGATCC | vif-vpr-vpu | First sweep of PCRs general for HIV-1 group M | Did not yield sequence data |
| HY27R | TTGGTACAAGCAGTYYTAGG | vif-vpr-vpu | First sweep of PCRs general for HIV-1 group M | Did not yield sequence data |
| HY28F | CCTGGAAGCATCCAGGA | vif-vpr-vpu | First sweep of PCRs general for HIV-1 group M | Did not yield sequence data |
| HY28R | CTTGGCAATGAARGCARC | vif-vpr-vpu | First sweep of PCRs general for HIV-1 group M | Did not yield sequence data |
| HY29F | CCTARRACTGCTTGTACCA | vif-vpr-vpu | First sweep of PCRs general for HIV-1 group M | Did not yield sequence data |
| HY29R | TAGGAGATGCCTAAGSCTT | vif-vpr-vpu | First sweep of PCRs general for HIV-1 group M | Did not yield sequence data |
| HY30F | GYTGCYTTCATTGCCAAG | vif-vpr-vpu | First sweep of PCRs general for HIV-1 group M | Did not yield sequence data |
| HY30R | CTGTCTCCGCTTCTTCC | vif-vpr-vpu | First sweep of PCRs general for HIV-1 group M | Did not yield sequence data |
| HY31F | AAGSCTTAGGCATCTCCT | vif-vpr-vpu | First sweep of PCRs general for HIV-1 group M | Did not yield sequence data |
| HY31R | GCTTGATGAGTCTKACWGTC | vif-vpr-vpu | First sweep of PCRs general for HIV-1 group M | Did not yield sequence data |
| HY37F | AKAGAAARAGCAGAAGACAG | vif-vpr-vpu | First sweep of PCRs general for HIV-1 group M | Yielded sequence data |
| HY37R | AGRTGCCCCMTCTCCA | vif-vpr-vpu | First sweep of PCRs general for HIV-1 group M | Yielded sequence data |

| | | | | |
|---|---|---|---|---|
| HY38F | GTGAAGGRGAYCAGGAAG | vif-vpr-vpu | First sweep of PCRs general for HIV-1 group M | Did not yield sequence data |
| HY38R | GTTCTGYAGCASTACAGATC | vif-vpr-vpu | First sweep of PCRs general for HIV-1 group M | Did not yield sequence data |
| HY39F | GGCAYCATGCTCCTTG | vif-vpr-vpu | First sweep of PCRs general for HIV-1 group M | Did not yield sequence data |
| HY39R | ACCCCATAATAKACTGTGAC | vif-vpr-vpu | First sweep of PCRs general for HIV-1 group M | Did not yield sequence data |
| HY3F | GGATGAGGATTARAACATGG | vif-vpr-vpu | First sweep of PCRs general for HIV-1 group M | Yielded sequence data |
| HY3R | GGGKGYTTTCATAGTGATG | vif-vpr-vpu | First sweep of PCRs general for HIV-1 group M | Yielded sequence data |
| HY40F | GCTCCTTGGGATRTTRATG | vif-vpr-vpu | First sweep of PCRs general for HIV-1 group M | Did not yield sequence data |
| HY40R | GKTTGCTTCTTTCCAYACAG | vif-vpr-vpu | First sweep of PCRs general for HIV-1 group M | Did not yield sequence data |
| HY41F | CACAGTMTATTATGGGGTAC | vif-vpr-vpu | First sweep of PCRs general for HIV-1 group M | Yielded sequence data |
| HY41R | GCTYTAGCATCTGAWGCA | vif-vpr-vpu | First sweep of PCRs general for HIV-1 group M | Yielded sequence data |
| HY42F | GGAAAGAAGCAAMCACCA | vif-vpr-vpu | First sweep of PCRs general for HIV-1 group M | Did not yield sequence data |
| HY42R | GGCCCAAACATTATGYACC | vif-vpr-vpu | First sweep of PCRs general for HIV-1 group M | Did not yield sequence data |
| HY4F | CAAAGAAAGCTAARGGATGG | vif-vpr-vpu | First sweep of PCRs general for HIV-1 group M | Did not yield sequence data |
| HY4R | TACTTCTGAACTTAYTYTTG | vif-vpr-vpu | First sweep of PCRs general for HIV-1 group M | Did not yield sequence data |
| HY5F | RGGATGGKTTTATAGACATC | vif-vpr-vpu | First sweep of PCRs general for HIV-1 group M | Yielded sequence data |
| HY5R | CATCCCCTARTGGGAYGTG | vif-vpr-vpu | First sweep of PCRs general for HIV-1 group M | Yielded sequence data |
| HY6F | TGAAARCMCCCATCCAA | vif-vpr-vpu | First sweep of PCRs general for HIV-1 group M | Did not yield sequence data |
| HY6R | CCTGTATGCAGAMCCCAA | vif-vpr-vpu | First sweep of PCRs general for HIV-1 group M | Did not yield sequence data |
| HY7F | CTAGRGGATGCTARATTGG | vif-vpr-vpu | First sweep of PCRs general for HIV-1 group M | Did not yield sequence data |
| HY7R | CYTGRCCCAAATGCCAGT | vif-vpr-vpu | First sweep of PCRs general for HIV-1 group M | Did not yield sequence data |
| HY8F | TCTGCATACAGGAGAAAGAG | vif-vpr-vpu | First sweep of PCRs general for HIV-1 group M | Did not yield sequence data |
| HY8R | AGGGTCTACTTGTGTGCT | vif-vpr-vpu | First sweep of PCRs general for HIV-1 group M | Did not yield sequence data |
| HY9F | CATTTGGGYCARGGAGTCT | vif-vpr-vpu | First sweep of PCRs general for HIV-1 group M | Did not yield sequence data |
| HY9R | GCTAGGYCAGGGTCTAC | vif-vpr-vpu | First sweep of PCRs general for HIV-1 group M | Did not yield sequence data |
| KE10F | GCAATACCTCARCCRT | env | Second sweep of PCRs general for subtype C | Did not yield sequence data |
| KE10R | CATAACCAGCTGGRGYAC | env | Second sweep of PCRs general for subtype C | Did not yield sequence data |
| KE11F | ACAAGCYTGTCCAAARG | env | Second sweep of PCRs general for subtype C | Yielded sequence data |
| KE11R | TAGAATYGCATAACCAGCT | env | Second sweep of PCRs general for subtype C | Yielded sequence data |
| KE12F | CCAAAGGTCWCTTTTGAYCC | env | Second sweep of PCRs general for subtype C | Yielded sequence data |
| KE12R | CCTGTYCCATTGAATGYCT | env | Second sweep of PCRs general for subtype C | Yielded sequence data |
| KE13F | TTATTGTRCYCCAGCTGGT | env | Second sweep of PCRs general for subtype C | Yielded sequence data |
| KE13R | TAKTGCATGKTCCTGT | env | Second sweep of PCRs general for subtype C | Yielded sequence data |
| KE14F | AAGACATTCAATGGRWCAG | env | Second sweep of PCRs general for subtype C | Yielded sequence data |
| KE14R | ACCACTRGCTTRATTCCAT | env | Second sweep of PCRs general for subtype C | Yielded sequence data |
| KE15F | AAGGACCATGCAMKAATGTC | env | Second sweep of PCRs general for subtype C | Yielded sequence data |
| KE15R | CAGCARCTGAGTTGAYAC | env | Second sweep of PCRs general for subtype C | Yielded sequence data |
| KE16F | CAGTACAATGTACACATGGA | env | Second sweep of PCRs general for subtype C | Yielded sequence data |
| KE16R | TCTTCTGCTAGRCTRCCAT | env | Second sweep of PCRs general for subtype C | Yielded sequence data |
| KE17F | TATCAACTCAGYTRCTGT | env | Second sweep of PCRs general for subtype C | Yielded sequence data |
| KE17R | GTTCGTYAKATTTTYAGATC | env | Second sweep of PCRs general for subtype C | Yielded sequence data |
| KE18F | AATGGYAGYCTAGCAGAAG | env | Second sweep of PCRs general for subtype C | Did not yield sequence data |
| KE18R | CAGATTCRTTMAGMTGTAC | env | Second sweep of PCRs general for subtype C | Did not yield sequence data |
| KE19F | TGTACAAGGCCYRGCA | env | Second sweep of PCRs general for subtype C | Yielded sequence data |
| KE19R | CCTATTATRTCWCCTGTTGC | env | Second sweep of PCRs general for subtype C | Yielded sequence data |
| KE1F | CTATTYTGTGCRTCAGATGC | env | Second sweep of PCRs general for subtype C | Yielded sequence data |
| KE1R | GGTACACAGGCATGKRT | env | Second sweep of PCRs general for subtype C | Yielded sequence data |
| KE20F | CCAGGACAAACATTYTWTGC | env | Second sweep of PCRs general for subtype C | Did not yield sequence data |
| KE20R | TACCTACCTCTTKTAARGTT | env | Second sweep of PCRs general for subtype C | Did not yield sequence data |
| KE21F | AAACTTTACAAGAGGTARRT | env | Second sweep of PCRs general for subtype C | Did not yield sequence data |
| KE21R | TTCYARGTCCCCTCCTGA | env | Second sweep of PCRs general for subtype C | Did not yield sequence data |
| KE22F | GGGGACYTRGAAATTACAAC | env | Second sweep of PCRs general for subtype C | Did not yield sequence data |
| KE22R | CCTCCTGTACTATTAAACMRT | env | Second sweep of PCRs general for subtype C | Did not yield sequence data |
| KE23F | ATCACACTCCMATGCAGA | env | Second sweep of PCRs general for subtype C | Yielded sequence data |
| KE23R | GGGAGGRGCATAYATTGC | env | Second sweep of PCRs general for subtype C | Yielded sequence data |
| KE24F | AAATATGTGGCAGGRRGTAG | env | Second sweep of PCRs general for subtype C | Yielded sequence data |
| KE24R | GTGTCAATAKTAGTCCTGTG | env | Second sweep of PCRs general for subtype C | Yielded sequence data |
| KE25F | CCCTCCCAYTGMAGGAA | env | Second sweep of PCRs general for subtype C | Yielded sequence data |
| KE25R | CTCCTCCATCMCKTGTCA | env | Second sweep of PCRs general for subtype C | Yielded sequence data |
| KE26F | TCACAGGACTAMTATTGACA | env | Second sweep of PCRs general for subtype C | Did not yield sequence data |
| KE26R | TCCYCCTCCAGGTCTG | env | Second sweep of PCRs general for subtype C | Did not yield sequence data |
| KE27F | TACTATTRACAMGTGATGGA | env | Second sweep of PCRs general for subtype C | Yielded sequence data |
| KE27R | ATATYTCCYCCTCCAGGT | env | Second sweep of PCRs general for subtype C | Yielded sequence data |
| KE28F | ATGARAYATTCAGACCTGGA | env | Second sweep of PCRs general for subtype C | Yielded sequence data |
| KE28R | GTKGGTGCTAYTCCYAATGG | env | Second sweep of PCRs general for subtype C | Yielded sequence data |
| KE29F | TGAGRGACAATTGGAGA | env | Second sweep of PCRs general for subtype C | Yielded sequence data |
| KE29R | TTGCCYTRGTGGGTGCT | env | Second sweep of PCRs general for subtype C | Yielded sequence data |
| KE2F | GTGCATAATGTYTGGGCYA | env | Second sweep of PCRs general for subtype C | Did not yield sequence data |
| KE2R | TCCACAWCYATYTCTTG | env | Second sweep of PCRs general for subtype C | Did not yield sequence data |
| KE30F | TTAARCCATTRGGRATAGC | env | Second sweep of PCRs general for subtype C | Did not yield sequence data |
| KE30R | GAACAYAGCTCCTAKTC | env | Second sweep of PCRs general for subtype C | Did not yield sequence data |
| KE31F | AARAGRAGAGTGGTGGAGA | env | Second sweep of PCRs general for subtype C | Yielded sequence data |
| KE31R | CGCCCATAGTGCTTCC | env | Second sweep of PCRs general for subtype C | Yielded sequence data |
| KE32F | CGWTCCTTGGGTTCTTRG | env | Second sweep of PCRs general for subtype C | Yielded sequence data |
| KE32R | RCAATTGTCTRGCCTGT | env | Second sweep of PCRs general for subtype C | Yielded sequence data |
| KE33F | GCGCAGCRTCAATAACG | env | Second sweep of PCRs general for subtype C | Yielded sequence data |
| KE33R | TCTATRGCTYTCAGCAA | env | Second sweep of PCRs general for subtype C | Yielded sequence data |

| | | | | |
|---|---|---|---|---|
| KE34F | YTGGYATAGTGCAACAGCA | env | Second sweep of PCRs general for subtype C | Yielded sequence data |
| KE34R | CAGACYGTGAGTTKCAA | env | Second sweep of PCRs general for subtype C | Yielded sequence data |
| KE35F | ATAGAGGCKTAACARCA | env | Second sweep of PCRs general for subtype C | Did not yield sequence data |
| KE35R | CTTTCTATAGCCARGAYTCT | env | Second sweep of PCRs general for subtype C | Did not yield sequence data |
| KE36F | TGGGGCATWAARCAGCT | env | Second sweep of PCRs general for subtype C | Yielded sequence data |
| KE36R | AAATYCCTAGGAGCTGTTG | env | Second sweep of PCRs general for subtype C | Yielded sequence data |
| KE37F | CCTGGCTRTRGAAAGATACC | env | Second sweep of PCRs general for subtype C | Yielded sequence data |
| KE37R | CAGTGGTGCARATGWGT | env | Second sweep of PCRs general for subtype C | Yielded sequence data |
| KE38F | CRGGATCAACAGCTCCTAG | env | Second sweep of PCRs general for subtype C | Yielded sequence data |
| KE38R | CTCCAACTAWTRTTCCAAGG | env | Second sweep of PCRs general for subtype C | Yielded sequence data |
| KE39F | AACWCATYTGCACCACT | env | Second sweep of PCRs general for subtype C | Yielded sequence data |
| KE39R | GCATCCARGTCATGTTWTYC | env | Second sweep of PCRs general for subtype C | Yielded sequence data |
| KE3F | TACCYACMGACCCCAAC | env | Second sweep of PCRs general for subtype C | Yielded sequence data |
| KE3R | CATGCATCTGMTCYACCA | env | Second sweep of PCRs general for subtype C | Yielded sequence data |
| KE40F | GCCTTGGAACAWTASTTGG | env | Second sweep of PCRs general for subtype C | Yielded sequence data |
| KE40R | CCTGTATATTGCACCTGTG | env | Second sweep of PCRs general for subtype C | Did not yield sequence data |
| KE41F | GGGAWAACATGACYTGGATG | env | Second sweep of PCRs general for subtype C | Did not yield sequence data |
| KE41R | TTCCTGCTGGWTYTGYG | env | Second sweep of PCRs general for subtype C | Did not yield sequence data |
| KE42F | GGGATAGAGARATTARYAA | env | Second sweep of PCRs general for subtype C | Did not yield sequence data |
| KE42R | CCAAYTGTCCAATKCYA | env | Second sweep of PCRs general for subtype C | Did not yield sequence data |
| KE43F | TACTAGMATTGGACARKTGG | env | Second sweep of PCRs general for subtype C | Did not yield sequence data |
| KE43R | AAACCTATYAARCCYCCTAC | env | Second sweep of PCRs general for subtype C | Did not yield sequence data |
| KE44F | GATAGTAGGRGGYTTRATAGG | env | Second sweep of PCRs general for subtype C | Yielded sequence data |
| KE44R | ACGACARAGGTGARTATCC | env | Second sweep of PCRs general for subtype C | Yielded sequence data |
| KE45F | RAATAGAGTTAGGCAGGGAT | env | Second sweep of PCRs general for subtype C | Yielded sequence data |
| KE45R | CACCTTCTTCTTCKATTCYT | env | Second sweep of PCRs general for subtype C | Yielded sequence data |
| KE46F | TCGTTTCAGACCCTYAYCC | env | Second sweep of PCRs general for subtype C | Did not yield sequence data |
| KE46R | ATCTGTYTCTGTCTYGCTCT | env | Second sweep of PCRs general for subtype C | Did not yield sequence data |
| KE47F | GAGGAATMGAAGRAGAAGG | env | Second sweep of PCRs general for subtype C | Did not yield sequence data |
| KE47R | AGRTCGTCCCAGRCRA | env | Second sweep of PCRs general for subtype C | Did not yield sequence data |
| KE48F | GTGGAGAGCRAGRCAGAG | env | Second sweep of PCRs general for subtype C | Yielded sequence data |
| KE48R | GTAGAKGAARAKGCACAG | env | Second sweep of PCRs general for subtype C | Yielded sequence data |
| KE49F | GACTYGYCTGGGACGA | env | Second sweep of PCRs general for subtype C | Did not yield sequence data |
| KE49R | CTCCGTASCAGAAKYTCCA | env | Second sweep of PCRs general for subtype C | Did not yield sequence data |
| KE4F | GGAAAAATGAYATGGTRGA | env | Second sweep of PCRs general for subtype C | Yielded sequence data |
| KE4R | AGYGGGGTYAACTTTACAC | env | Second sweep of PCRs general for subtype C | Yielded sequence data |
| KE50F | CTCTACCACCRATTGAGRG | env | Second sweep of PCRs general for subtype C | Yielded sequence data |
| KE50R | CAGATAYTTRAGRGCTTCC | env | Second sweep of PCRs general for subtype C | Yielded sequence data |
| KE51F | GGGCRGTGGARMTTCTG | env | Second sweep of PCRs general for subtype C | Yielded sequence data |
| KE51R | CARACCCCAATAYTGYA | env | Second sweep of PCRs general for subtype C | Yielded sequence data |
| KE52F | AGGTCTTSTRCARTATTGG | env | Second sweep of PCRs general for subtype C | Yielded sequence data |
| KE52R | CYCAGCTACTRCTATKGCT | env | Second sweep of PCRs general for subtype C | Yielded sequence data |
| KE53F | GGCCTGGAACTAAAAAWKAG | env | Second sweep of PCRs general for subtype C | Yielded sequence data |
| KE53R | AATCCTATCTGTYCCYYCAG | env | Second sweep of PCRs general for subtype C | Yielded sequence data |
| KE54F | CCATAGCMATAGYAGTAGC | env | Second sweep of PCRs general for subtype C | Yielded sequence data |
| KE54R | CCCTGTCTTATTCKTCTRGG | env | Second sweep of PCRs general for subtype C | Yielded sequence data |
| KE55F | AATAGCAGTAGCTGARGGRA | env | Second sweep of PCRs general for subtype C | Yielded sequence data |
| KE55R | CCCTGTCTTATTCKTSTRGGT | env | Second sweep of PCRs general for subtype C | Yielded sequence data |
| KE56F | GTAGAGCTATCYAYMACATACC | env | Second sweep of PCRs general for subtype C | Yielded sequence data |
| KE56Fb | TTAGAGTTATCCTCAACATACC | env | Second sweep of PCRs general for subtype C | Yielded sequence data |
| KE56R | CCMCCCATTTTATWGYAA | env | Second sweep of PCRs general for subtype C | Yielded sequence data |
| KE57F | ACAMGRATAAGACAGGGYTT | env | Second sweep of PCRs general for subtype C | Did not yield sequence data |
| KE57R | CCATCCAACTATRCTRCTT | env | Second sweep of PCRs general for subtype C | Yielded sequence data |
| KE58F | ATGGGKGGYAARTGGT | env | Second sweep of PCRs general for subtype C | Did not yield sequence data |
| KE58R | CTGCTGCKGGMTCAGT | env | Second sweep of PCRs general for subtype C | Did not yield sequence data |
| KE59F | AGCAGYATAGTKGGATG | env | Second sweep of PCRs general for subtype C | Did not yield sequence data |
| KE59R | CCCCATGTTTWTCYARGTCT | env | Second sweep of PCRs general for subtype C | Did not yield sequence data |
| KE5F | ACCAAAGCCTAAAGCCAT | env | Second sweep of PCRs general for subtype C | Did not yield sequence data |
| KE5R | CTGGTAACATTAGCATYWC | env | Second sweep of PCRs general for subtype C | Did not yield sequence data |
| KE60F | AACTGAKCCMGCRGCA | env | Second sweep of PCRs general for subtype C | Did not yield sequence data |
| KE60R | GCTGTRTTGCTRSTTG | env | Second sweep of PCRs general for subtype C | Did not yield sequence data |
| KE61F | AGTRGGARCAGCRTCTC | env | Second sweep of PCRs general for subtype C | Did not yield sequence data |
| KE61R | AGCCAGGCACARKCA | env | Second sweep of PCRs general for subtype C | Did not yield sequence data |
| KE62F | CTTACAASYAGCAAYACAG | env | Second sweep of PCRs general for subtype C | Did not yield sequence data |
| KE62R | CTACCTCYTCYTCSTCCTCT | env | Second sweep of PCRs general for subtype C | Did not yield sequence data |
| KE63F | CYTGTGCCTGGCTRGA | env | Second sweep of PCRs general for subtype C | Did not yield sequence data |
| KE63R | GTCATTGGTCTTAAMGGYAC | env | Second sweep of PCRs general for subtype C | Did not yield sequence data |
| KE64F | GTRGGYTTTCCAGTCARA | env | Second sweep of PCRs general for subtype C | Did not yield sequence data |
| KE64R | CWTCCAGTCCCCCCTT | env | Second sweep of PCRs general for subtype C | Did not yield sequence data |
| KE65F | GGTRCCKTTAAGACCAATGAC | env | Second sweep of PCRs general for subtype C | Did not yield sequence data |
| KE65R | CTTGGAGYAAATTARCCCWTCC | env | Second sweep of PCRs general for subtype C | Did not yield sequence data |
| KE66F | TTAARAGAAAAGGGGGGACT | env | Second sweep of PCRs general for subtype C | Did not yield sequence data |
| KE66R | GTAGCCTTGTGTGTKRTAGAC | env | Second sweep of PCRs general for subtype C | Did not yield sequence data |
| KE66Rb | GTAGCCTTGTGTGTTATAGAC | env | Second sweep of PCRs general for subtype C | Did not yield sequence data |
| KE67F | GAARAGRCAAGAKATCCTTG | env | Second sweep of PCRs general for subtype C | Did not yield sequence data |
| KE67R | CCTGGYCCYGGTGTGT | env | Second sweep of PCRs general for subtype C | Did not yield sequence data |

| | | | | |
|---|---|---|---|---|
| KE6F | GACCCCRCTCTGTGTCA | env | Second sweep of PCRs general for subtype C | Did not yield sequence data |
| KE6R | TTTCTCCMTTCATGGT | env | Second sweep of PCRs general for subtype C | Did not yield sequence data |
| KE7F | TTGCTCTTTCAATRYAACC | env | Second sweep of PCRs general for subtype C | Did not yield sequence data |
| KE7R | CTCCTTAAKTGGTACTAYATC | env | Second sweep of PCRs general for subtype C | Did not yield sequence data |
| KE8F | GCAACCACAGAAMTAAGRG | env | Second sweep of PCRs general for subtype C | Did not yield sequence data |
| KE8R | ACTCACTAGAGYTGYT | env | Second sweep of PCRs general for subtype C | Did not yield sequence data |
| KE9F | GTAGTACCAMTTAAGGAGA | env | Second sweep of PCRs general for subtype C | Did not yield sequence data |
| KE9R | CTTTGGACARGCYTGTG | env | Second sweep of PCRs general for subtype C | Did not yield sequence data |
| KG10F | ARAGACACCAARGAAGC | gag | Second sweep of PCRs general for subtype C | Yielded sequence data |
| KG10R | CCTGCTGTGYTTTTTKC | gag | Second sweep of PCRs general for subtype C | Yielded sequence data |
| KG11F | GCCTTAGACAARATARAGGA | gag | Second sweep of PCRs general for subtype C | Did not yield sequence data |
| KG11R | TTTCCTYYGTCAGCYGCT | gag | Second sweep of PCRs general for subtype C | Did not yield sequence data |
| KG12F | AAACACAGCAGGCARRA | gag | Second sweep of PCRs general for subtype C | Did not yield sequence data |
| KG12R | TTGCCCCTGKARATTCTG | gag | Second sweep of PCRs general for subtype C | Did not yield sequence data |
| KG13F | CAGAATATMCARGGGCAA | gag | Second sweep of PCRs general for subtype C | Did not yield sequence data |
| KG13R | GGCTRAAAGCCTTYTCCT | gag | Second sweep of PCRs general for subtype C | Did not yield sequence data |
| KG14F | ATCAGCCCATRTCASCTAG | gag | Second sweep of PCRs general for subtype C | Did not yield sequence data |
| KG14R | GTATTACCTCTGGRCTRA | gag | Second sweep of PCRs general for subtype C | Did not yield sequence data |
| KG15F | GTAGAGGAAAAGGCYTTYAG | gag | Second sweep of PCRs general for subtype C | Did not yield sequence data |
| KG15R | TTGTGGGRTGGCTCCT | gag | Second sweep of PCRs general for subtype C | Did not yield sequence data |
| KG16F | CCCAGARGTAATACCCATG | gag | Second sweep of PCRs general for subtype C | Yielded sequence data |
| KG16R | CCCCCACTGTRTTTARCAT | gag | Second sweep of PCRs general for subtype C | Yielded sequence data |
| KG17F | AAGGAGCCAYCCCACA | gag | Second sweep of PCRs general for subtype C | Yielded sequence data |
| KG17R | CATYTGCATRGCTGCTTG | gag | Second sweep of PCRs general for subtype C | Yielded sequence data |
| KG18F | TAAAYACAGTRGGGGGACA | gag | Second sweep of PCRs general for subtype C | Did not yield sequence data |
| KG18R | ATTCTGCAGCYTCYTCA | gag | Second sweep of PCRs general for subtype C | Did not yield sequence data |
| KG19F | CATCAAGCAGCYATGCA | gag | Second sweep of PCRs general for subtype C | Did not yield sequence data |
| KG19R | CATGCACTGGATGTARYCT | gag | Second sweep of PCRs general for subtype C | Did not yield sequence data |
| KG20F | AATGARGARGCTGCAGA | gag | Second sweep of PCRs general for subtype C | Did not yield sequence data |
| KG20R | ACTTCCCCTTGGYTCTC | gag | Second sweep of PCRs general for subtype C | Did not yield sequence data |
| KG21F | GCACCAGGYCARATGAG | gag | Second sweep of PCRs general for subtype C | Did not yield sequence data |
| KG21R | GGGTTRCCTGTCAYCCAT | gag | Second sweep of PCRs general for subtype C | Did not yield sequence data |
| KG22F | AGGAACWACTAGTAMCCTTC | gag | Second sweep of PCRs general for subtype C | Yielded sequence data |
| KG22R | GCCCCAGRATTATCCAYC | gag | Second sweep of PCRs general for subtype C | Yielded sequence data |
| KG23F | GTTCCAGTRGGAGAMATC | gag | Second sweep of PCRs general for subtype C | Yielded sequence data |
| KG23R | GTCCAAAATGCTRAYAG | gag | Second sweep of PCRs general for subtype C | Yielded sequence data |
| KG24F | AGAATGTATAGCCCTRTYAG | gag | Second sweep of PCRs general for subtype C | Yielded sequence data |
| KG24R | AACCKGTCTACATAGTCTCT | gag | Second sweep of PCRs general for subtype C | Yielded sequence data |
| KG25F | CAAGGACCAAARGAACYCT | gag | Second sweep of PCRs general for subtype C | Yielded sequence data |
| KG25R | GTGTAGMTTGTTCRGCTCT | gag | Second sweep of PCRs general for subtype C | Yielded sequence data |
| KG26F | GAGAYTATGTAGACMGGTTC | gag | Second sweep of PCRs general for subtype C | Yielded sequence data |
| KG26R | AACAAGGTKTCKGTCATCC | gag | Second sweep of PCRs general for subtype C | Yielded sequence data |
| KG27F | GCTGAACAAGCTWCACARG | gag | Second sweep of PCRs general for subtype C | Did not yield sequence data |
| KG27R | GGTYTTACAATCTGGGTTYG | gag | Second sweep of PCRs general for subtype C | Yielded sequence data |
| KG28F | GAMACCTTGTTGGTYCAA | gag | Second sweep of PCRs general for subtype C | Did not yield sequence data |
| KG28R | CCCTGRCATGCTGTCATC | gag | Second sweep of PCRs general for subtype C | Yielded sequence data |
| KG29F | CCAGGGGCTWCAYTAGAAG | gag | Second sweep of PCRs general for subtype C | Yielded sequence data |
| KG29R | GCCTCAGCCAAMACYCT | gag | Second sweep of PCRs general for subtype C | Yielded sequence data |
| KG2F | GGGGGARAATTAGATGMATG | gag | Second sweep of PCRs general for subtype C | Yielded sequence data |
| KG2R | TCYAGCTCCYTGCTTGC | gag | Second sweep of PCRs general for subtype C | Yielded sequence data |
| KG30F | GACCTRGCCAYAAAGC | gag | Second sweep of PCRs general for subtype C | Did not yield sequence data |
| KG30R | AGGGCCYTTAAAATTGC | gag | Second sweep of PCRs general for subtype C | Did not yield sequence data |
| KG31F | CTGAGGCAATGAGYCAA | gag | Second sweep of PCRs general for subtype C | Did not yield sequence data |
| KG31R | CCCTTCCYTGCCACAG | gag | Second sweep of PCRs general for subtype C | Did not yield sequence data |
| KG32F | GTTTYAACTGTGGCARGG | gag | Second sweep of PCRs general for subtype C | Did not yield sequence data |
| KG32R | TGGTGTCCTTCYTTTCCA | gag | Second sweep of PCRs general for subtype C | Did not yield sequence data |
| KG33F | GGCYCCTAGGAAAARGGG | gag | Second sweep of PCRs general for subtype C | Yielded sequence data |
| KG33R | CCTGTCTYTCRGTACAGT | gag | Second sweep of PCRs general for subtype C | Yielded sequence data |
| KG34F | ATGTGGAAARGAAGGACAC | gag | Second sweep of PCRs general for subtype C | Yielded sequence data |
| KG34R | TTGGAAGGCCAAAKTYTC | gag | Second sweep of PCRs general for subtype C | Yielded sequence data |
| KG35F | GTACTGAGRGRCAGGCTA | gag | Second sweep of PCRs general for subtype C | Yielded sequence data |
| KG35R | CTGCTCTGAAGRAAATTYC | gag | Second sweep of PCRs general for subtype C | Yielded sequence data |
| KG36F | CCTTCCMACAAGGGRAGG | gag | Second sweep of PCRs general for subtype C | Did not yield sequence data |
| KG36R | TCTCTGGTGGGGCTGT | gag | Second sweep of PCRs general for subtype C | Yielded sequence data |
| KG37F | CCTTCAGARCAGRCCAGA | gag | Second sweep of PCRs general for subtype C | Yielded sequence data |
| KG37R | TTCCTCGAACYTGAAGCT | gag | Second sweep of PCRs general for subtype C | Did not yield sequence data |
| KG38F | CAGRCCAGAGCCAACAG | gag | Second sweep of PCRs general for subtype C | Yielded sequence data |
| KG38R | TTGTCCYTCGRCTCCT | gag | Second sweep of PCRs general for subtype C | Yielded sequence data |
| KG39F | CAACYCCCSCTCCGAAG | gag | Second sweep of PCRs general for subtype C | Yielded sequence data |
| KG39R | GTCGCTGCCAAAGAGTG | gag | Second sweep of PCRs general for subtype C | Yielded sequence data |
| KG3F | GTTAAGGCCAGGRGGRA | gag | Second sweep of PCRs general for subtype C | Yielded sequence data |
| KG3R | GGTCAGGGTYAASTGCAA | gag | Second sweep of PCRs general for subtype C | Did not yield sequence data |
| KG4F | AACACYTAGTMTGGGCAA | gag | Second sweep of PCRs general for subtype C | Yielded sequence data |
| KG4R | GTTTACAGCCTKCTGMTG | gag | Second sweep of PCRs general for subtype C | Did not yield sequence data |
| KG5F | GGGAGCTGGAAARATTYGC | gag | Second sweep of PCRs general for subtype C | Yielded sequence data |
| KG5R | GAAGAGMTGGTTGTARCT | gag | Second sweep of PCRs general for subtype C | Yielded sequence data |

| | | | | |
|---|---|---|---|---|
| KG6F | GATACATCAGMAGGMTGT | gag | Second sweep of PCRs general for subtype C | Yielded sequence data |
| KG6R | AGTTCCTCTGWTCCTGTYT | gag | Second sweep of PCRs general for subtype C | Yielded sequence data |
| KG7F | AGCTACAACCAKCTMTYC | gag | Second sweep of PCRs general for subtype C | Did not yield sequence data |
| KG7R | CAATAGAGRGTTGCYACTG | gag | Second sweep of PCRs general for subtype C | Did not yield sequence data |
| KG8F | CAGACAGGAACARARGAAC | gag | Second sweep of PCRs general for subtype C | Did not yield sequence data |
| KG8R | TAARGCTTCYTTGGTGTCT | gag | Second sweep of PCRs general for subtype C | Did not yield sequence data |
| KG9F | GTAGCARCYCTCTATTGTG | gag | Second sweep of PCRs general for subtype C | Yielded sequence data |
| KG9R | CTATCTTRTCTAARGCTTCC | gag | Second sweep of PCRs general for subtype C | Yielded sequence data |
| DP10F | TGTGGAAAMARGGCTATAGG | pol | Second sweep of PCRs general for subtype C | Yielded sequence data |
| DP10R | CTGAGTCAACAKATTYCTTC | pol | Second sweep of PCRs general for subtype C | Yielded sequence data |
| DP11F | GGACCYACRCCTGTCAAC | pol | Second sweep of PCRs general for subtype C | Did not yield sequence data |
| DP11R | GTTTCAATRGGACTAATKGG | pol | Second sweep of PCRs general for subtype C | Did not yield sequence data |
| DP12F | GAATMTGTTGACTCAGMTTG | pol | Second sweep of PCRs general for subtype C | Did not yield sequence data |
| DP12R | CCATCCATYCCTGGCTT | pol | Second sweep of PCRs general for subtype C | Did not yield sequence data |
| DP13F | TAGTCCYATTGAAACTG | pol | Second sweep of PCRs general for subtype C | Did not yield sequence data |
| DP13R | TTYTCTTCTGTCAATGG | pol | Second sweep of PCRs general for subtype C | Did not yield sequence data |
| DP14F | ATGGATGGCCCAAARGT | pol | Second sweep of PCRs general for subtype C | Yielded sequence data |
| DP14R | CCTTCTTTCTCCATTTCWTC | pol | Second sweep of PCRs general for subtype C | Did not yield sequence data |
| DP15F | GGCCATTGACAGAAGARA | pol | Second sweep of PCRs general for subtype C | Yielded sequence data |
| DP15R | ATATGGATTYTCAGGCCCAA | pol | Second sweep of PCRs general for subtype C | Yielded sequence data |
| DP16F | ACAAAAATTGGGCCTGA | pol | Second sweep of PCRs general for subtype C | Yielded sequence data |
| DP16R | ACTTTCTCCACTTRGTACTG | pol | Second sweep of PCRs general for subtype C | Yielded sequence data |
| DP17F | AAYACTCCARTATTTGC | pol | Second sweep of PCRs general for subtype C | Yielded sequence data |
| DP17R | AGTTCYCTGAARTCTAC | pol | Second sweep of PCRs general for subtype C | Did not yield sequence data |
| DP18F | AGGACAGTACYAARTGGAGA | pol | Second sweep of PCRs general for subtype C | Yielded sequence data |
| DP18R | AACTTCCCARAARTCTTGAG | pol | Second sweep of PCRs general for subtype C | Yielded sequence data |
| DP19F | GTTAGTAGAYTTCAGRGAAC | pol | Second sweep of PCRs general for subtype C | Yielded sequence data |
| DP19R | CTGGRTGYGGTATTCCTA | pol | Second sweep of PCRs general for subtype C | Yielded sequence data |
| DP1F | TAGGGAAGMTYTGGCCTTC | pol | Second sweep of PCRs general for subtype C | Did not yield sequence data |
| DP1R | TGGGGCTGTYGGCTCT | pol | Second sweep of PCRs general for subtype C | Did not yield sequence data |
| DP20F | TCAAGACTTYTGGGARGTTC | pol | Second sweep of PCRs general for subtype C | Yielded sequence data |
| DP20R | CATCYAGTACTGTYACTGA | pol | Second sweep of PCRs general for subtype C | Yielded sequence data |
| DP21F | CACAYCCAGCRGGGTT | pol | Second sweep of PCRs general for subtype C | Yielded sequence data |
| DP21R | GCAGTATAYTTCCTGAARTC | pol | Second sweep of PCRs general for subtype C | Did not yield sequence data |
| DP22F | GGGGATGCATATTTYTCAGT | pol | Second sweep of PCRs general for subtype C | Yielded sequence data |
| DP22R | TACTAGGTATGGTRAATGCA | pol | Second sweep of PCRs general for subtype C | Yielded sequence data |
| DP23F | GGATGTGGGKGATGCA | pol | Second sweep of PCRs general for subtype C | Did not yield sequence data |
| DP23R | CTAGGTATGGTRAATGC | pol | Second sweep of PCRs general for subtype C | Did not yield sequence data |
| DP24F | AGATGARGAYTTCAGGA | pol | Second sweep of PCRs general for subtype C | Yielded sequence data |
| DP24R | ATCCCTGTGGAAGYACA | pol | Second sweep of PCRs general for subtype C | Did not yield sequence data |
| DP25F | CTGCATTYACCATACCT | pol | Second sweep of PCRs general for subtype C | Did not yield sequence data |
| DP25R | TGMTGGTGATCCTTTCC | pol | Second sweep of PCRs general for subtype C | Yielded sequence data |
| DP26F | GTRCTTCCACAGGGWTG | pol | Second sweep of PCRs general for subtype C | Did not yield sequence data |
| DP26R | TCTTGCYCTAAARGGCTCT | pol | Second sweep of PCRs general for subtype C | Yielded sequence data |
| DP27F | CCCTTTAGRGCAMRAAATCC | pol | Second sweep of PCRs general for subtype C | Yielded sequence data |
| DP27R | TGCTCTATGYTGYCCTA | pol | Second sweep of PCRs general for subtype C | Yielded sequence data |
| DP28F | TAGGGCAACATAGRRCA | pol | Second sweep of PCRs general for subtype C | Did not yield sequence data |
| DP28R | TGTTTCTTRTCYGGTGT | pol | Second sweep of PCRs general for subtype C | Did not yield sequence data |
| DP29F | CATCTGTTAARGTGGGGAT | pol | Second sweep of PCRs general for subtype C | Did not yield sequence data |
| DP29R | TCAGGATGGAGTTCATAMC | pol | Second sweep of PCRs general for subtype C | Did not yield sequence data |
| DP2F | GGGAATTTYCTTCAGARCAG | pol | Second sweep of PCRs general for subtype C | Did not yield sequence data |
| DP2R | CTTTCGGCTCCKGYTTC | pol | Second sweep of PCRs general for subtype C | Did not yield sequence data |
| DP30F | GAACCYCCATTTCTTTGGA | pol | Second sweep of PCRs general for subtype C | Yielded sequence data |
| DP30R | TGGCARCTKTATAGGCTGT | pol | Second sweep of PCRs general for subtype C | Yielded sequence data |
| DP31F | GTATGAACTCCATCCTGAYA | pol | Second sweep of PCRs general for subtype C | Did not yield sequence data |
| DP31R | TCATTGACAGTCCAGCT | pol | Second sweep of PCRs general for subtype C | Did not yield sequence data |
| DP32F | AGAGYTGCCAGAAAARG | pol | Second sweep of PCRs general for subtype C | Yielded sequence data |
| DP32R | CCTGRTAAATCTGRCT | pol | Second sweep of PCRs general for subtype C | Did not yield sequence data |
| DP33F | TAAAYTGGGCMAGYCAG | pol | Second sweep of PCRs general for subtype C | Did not yield sequence data |
| DP33R | TGCTTTGGYCCCCCTA | pol | Second sweep of PCRs general for subtype C | Did not yield sequence data |
| DP34F | MAGTCAGATTTACCCAGGA | pol | Second sweep of PCRs general for subtype C | Yielded sequence data |
| DP34R | GCTTCTTCWGTTAGTGGTAC | pol | Second sweep of PCRs general for subtype C | Yielded sequence data |
| DP35F | TAGGGGGRCCAAAGCA | pol | Second sweep of PCRs general for subtype C | Yielded sequence data |
| DP35R | TTCYCTGTTYTCTGCCAA | pol | Second sweep of PCRs general for subtype C | Yielded sequence data |
| DP36F | GTACCACTAACTGAAGAAGC | pol | Second sweep of PCRs general for subtype C | Did not yield sequence data |
| DP36R | CYCCATGTACTGGTTCTYT | pol | Second sweep of PCRs general for subtype C | Yielded sequence data |
| DP37F | TTGGCAGARAACAGRGAA | pol | Second sweep of PCRs general for subtype C | Did not yield sequence data |
| DP37R | GCTAYYAAGTCTTTTGATGG | pol | Second sweep of PCRs general for subtype C | Did not yield sequence data |
| DP38F | CTAARAGAACCAGTACATGG | pol | Second sweep of PCRs general for subtype C | Yielded sequence data |
| DP38R | CCCYTGTTTCTGTATTTCWG | pol | Second sweep of PCRs general for subtype C | Yielded sequence data |
| DP39F | CCCATCAAAAGACTTGRTAG | pol | Second sweep of PCRs general for subtype C | Did not yield sequence data |
| DP39R | CTTYCCTGTTTTCAGRT | pol | Second sweep of PCRs general for subtype C | Did not yield sequence data |
| DP3F | CRAACYTGAAGCTCTCTG | pol | Second sweep of PCRs general for subtype C | Did not yield sequence data |
| DP3R | CTGCCAAAGAGTGATYTRAG | pol | Second sweep of PCRs general for subtype C | Did not yield sequence data |
| DP40F | GGCAKGACCAATGGACA | pol | Second sweep of PCRs general for subtype C | Did not yield sequence data |
| DP40R | GGCAGTCCTCRTTYTTGC | pol | Second sweep of PCRs general for subtype C | Did not yield sequence data |

| | | | | |
|---|---|---|---|---|
| DP41F | AYCTGAAAACAGGRAAG | pol | Second sweep of PCRs general for subtype C | Did not yield sequence data |
| DP41R | CACWGCCTCTGTTAAYTG | pol | Second sweep of PCRs general for subtype C | Yielded sequence data |
| DP42F | GARGRCTGCCCACACT | pol | Second sweep of PCRs general for subtype C | Did not yield sequence data |
| DP42R | TAGGAGTCTTTCCCCAT | pol | Second sweep of PCRs general for subtype C | Did not yield sequence data |
| DP43F | GTTARCAGAGGCWGTGCA | pol | Second sweep of PCRs general for subtype C | Did not yield sequence data |
| DP43R | TCCACCATGTYTCCCATG | pol | Second sweep of PCRs general for subtype C | Did not yield sequence data |
| DP44F | ATGGGGAAAGAYTCCTA | pol | Second sweep of PCRs general for subtype C | Did not yield sequence data |
| DP44R | CAATAGTCTGYCCACCATG | pol | Second sweep of PCRs general for subtype C | Did not yield sequence data |
| DP45F | AGAAACATGGGARACATGG | pol | Second sweep of PCRs general for subtype C | Yielded sequence data |
| DP45R | GGAGGGGTATTRACAAAYTC | pol | Second sweep of PCRs general for subtype C | Yielded sequence data |
| DP46F | CACCTGGATTCCTGARTG | pol | Second sweep of PCRs general for subtype C | Yielded sequence data |
| DP46R | CTTTCTCYARCTGGTACCAT | pol | Second sweep of PCRs general for subtype C | Yielded sequence data |
| DP47F | GTTAATACCCCTCCYYT | pol | Second sweep of PCRs general for subtype C | Yielded sequence data |
| DP47R | CTATTAGCTGCYCCATCYAC | pol | Second sweep of PCRs general for subtype C | Yielded sequence data |
| DP48F | GAWCCCATAAYAGGAGCAGA | pol | Second sweep of PCRs general for subtype C | Yielded sequence data |
| DP48R | GTYAGTAACATAYCCTGCTT | pol | Second sweep of PCRs general for subtype C | Did not yield sequence data |
| DP49F | TAGATGGRGCAGCYAATAGG | pol | Second sweep of PCRs general for subtype C | Yielded sequence data |
| DP49R | TCTGCCTTCCTCTRTYAG | pol | Second sweep of PCRs general for subtype C | Yielded sequence data |
| DP4F | AGCAGGAGCCGAWAGAC | pol | Second sweep of PCRs general for subtype C | Yielded sequence data |
| DP4R | CCCCCTACTYTTATTGWGAC | pol | Second sweep of PCRs general for subtype C | Yielded sequence data |
| DP50F | GCAGGRTATGTTACTRA | pol | Second sweep of PCRs general for subtype C | Yielded sequence data |
| DP50R | GCTAGMTRAATTGCTTG | pol | Second sweep of PCRs general for subtype C | Did not yield sequence data |
| DP51F | CAGAAGACTGARTTRCAAGC | pol | Second sweep of PCRs general for subtype C | Yielded sequence data |
| DP51R | CTAATGCATAYTGTGARTCTG | pol | Second sweep of PCRs general for subtype C | Yielded sequence data |
| DP52F | GCARGATTCAGGAYCAGAAG | pol | Second sweep of PCRs general for subtype C | Yielded sequence data |
| DP52R | GTTGTGCTTGAATGATYCC | pol | Second sweep of PCRs general for subtype C | Yielded sequence data |
| DP53F | TAACAGAYTCACARTATGCA | pol | Second sweep of PCRs general for subtype C | Yielded sequence data |
| DP53R | GRTTRACTAACTCTGATTCAC | pol | Second sweep of PCRs general for subtype C | Yielded sequence data |
| DP54F | GAATCATTCARGCACARCC | pol | Second sweep of PCRs general for subtype C | Did not yield sequence data |
| DP54R | GACAGGTAGACYYTTTCCT | pol | Second sweep of PCRs general for subtype C | Did not yield sequence data |
| DP55F | CAGAGTTAGTYAAYCAA | pol | Second sweep of PCRs general for subtype C | Did not yield sequence data |
| DP55R | GTGCTGGTACCCATGAC | pol | Second sweep of PCRs general for subtype C | Did not yield sequence data |
| DP56F | AAGGAAARRGTCTACCTGTC | pol | Second sweep of PCRs general for subtype C | Yielded sequence data |
| DP56R | AYAGCACTTTCCTGATYCC | pol | Second sweep of PCRs general for subtype C | Yielded sequence data |
| DP57F | CCAGCACAYAAAGGRATTGG | pol | Second sweep of PCRs general for subtype C | Yielded sequence data |
| DP57R | CCTGRGCYTTATCTATTCCA | pol | Second sweep of PCRs general for subtype C | Yielded sequence data |
| DP58F | GGRATCAGGAAAGTGCTRT | pol | Second sweep of PCRs general for subtype C | Yielded sequence data |
| DP58R | GCTCTCCAATTGYTRTGA | pol | Second sweep of PCRs general for subtype C | Yielded sequence data |
| DP59F | RGCTCARGAAGAGCATG | pol | Second sweep of PCRs general for subtype C | Did not yield sequence data |
| DP59R | TTTGCTACTAYGGGTGGYAG | pol | Second sweep of PCRs general for subtype C | Did not yield sequence data |
| DP5F | CCCTYARATCACTCTTTGG | pol | Second sweep of PCRs general for subtype C | Did not yield sequence data |
| DP5R | TGTCTAAGAGAGCYTCYTT | pol | Second sweep of PCRs general for subtype C | Did not yield sequence data |
| DP60F | ATTGGAGARCAATGGCTAG | pol | Second sweep of PCRs general for subtype C | Yielded sequence data |
| DP60R | GCTGACAYTKATCACAGCT | pol | Second sweep of PCRs general for subtype C | Yielded sequence data |
| DP61F | TCTRCCACCCRTAGTAGCAA | pol | Second sweep of PCRs general for subtype C | Yielded sequence data |
| DP61R | CCATGYATGGCTTCYCCTT | pol | Second sweep of PCRs general for subtype C | Yielded sequence data |
| DP62F | GCTGTGATMARTGTCAGCTA | pol | Second sweep of PCRs general for subtype C | Yielded sequence data |
| DP62R | TTGCCATATYCCTGGACTRC | pol | Second sweep of PCRs general for subtype C | Yielded sequence data |
| DP63F | AAGGRGAAGCCATRCATGG | pol | Second sweep of PCRs general for subtype C | Yielded sequence data |
| DP63R | CATGRACTGCTACCAGRA | pol | Second sweep of PCRs general for subtype C | Yielded sequence data |
| DP64F | GYAGTCCAGGRATATGGCAA | pol | Second sweep of PCRs general for subtype C | Did not yield sequence data |
| DP64R | CTGCTTCYATRTARCCACT | pol | Second sweep of PCRs general for subtype C | Did not yield sequence data |
| DP65F | GGAAAAATYATYCTGGTAGC | pol | Second sweep of PCRs general for subtype C | Yielded sequence data |
| DP65R | TCCTGTTTCTGCTGGRAT | pol | Second sweep of PCRs general for subtype C | Yielded sequence data |
| DP66F | CAGTGGYTACATRGAAGCA | pol | Second sweep of PCRs general for subtype C | Yielded sequence data |
| DP66R | CTGGCCATCTTCCTGCTA | pol | Second sweep of PCRs general for subtype C | Yielded sequence data |
| DP67F | CCAGCAGAAACAGGRCA | pol | Second sweep of PCRs general for subtype C | Did not yield sequence data |
| DP67R | RCTRCCATTGTCTGT | pol | Second sweep of PCRs general for subtype C | Did not yield sequence data |
| DP68F | GCAGGAAGATGGCCAGT | pol | Second sweep of PCRs general for subtype C | Yielded sequence data |
| DP68R | TAYCYGCCCACCAACAG | pol | Second sweep of PCRs general for subtype C | Yielded sequence data |
| DP69F | GYAATTTCACCRGTGCT | pol | Second sweep of PCRs general for subtype C | Yielded sequence data |
| DP69R | CTGACTTTGGGGATTGTAG | pol | Second sweep of PCRs general for subtype C | Yielded sequence data |
| DP6F | GCGACCCCTYGTCWCA | pol | Second sweep of PCRs general for subtype C | Did not yield sequence data |
| DP6R | GTATCATCWGCTCCTGTRT | pol | Second sweep of PCRs general for subtype C | Did not yield sequence data |
| DP70F | TAAGGCMGCCTGYTGGT | pol | Second sweep of PCRs general for subtype C | Yielded sequence data |
| DP70R | GATTCYACTACTCCYTGACT | pol | Second sweep of PCRs general for subtype C | Yielded sequence data |
| DP71F | CTGGSATTCCCTACAATCC | pol | Second sweep of PCRs general for subtype C | Yielded sequence data |
| DP71R | TTGATCTCTTACCTGYCCTA | pol | Second sweep of PCRs general for subtype C | Yielded sequence data |
| DP72F | AGGGAGTAGTAGARTCYATG | pol | Second sweep of PCRs general for subtype C | Yielded sequence data |
| DP72R | TAAGRTGYTCAGCTTGATC | pol | Second sweep of PCRs general for subtype C | Yielded sequence data |
| DP73F | AATYATAGGRCAGGTAAGAG | pol | Second sweep of PCRs general for subtype C | Did not yield sequence data |
| DP73R | CACTRTAYCCCCCAATCC | pol | Second sweep of PCRs general for subtype C | Did not yield sequence data |
| DP74F | GGCAGTATTYATYCACA | pol | Second sweep of PCRs general for subtype C | Did not yield sequence data |
| DP74R | CTTTGTATGYCTGTTGC | pol | Second sweep of PCRs general for subtype C | Did not yield sequence data |
| DP75F | GCAGGGGAAAGAATARTAGAC | pol | Second sweep of PCRs general for subtype C | Did not yield sequence data |
| DP75R | GTCYCTGTAATAAACCCGRA | pol | Second sweep of PCRs general for subtype C | Did not yield sequence data |

| | | | | |
|---|---|---|---|---|
| DP76F | TYCGGGTTTATTACAGRGAC | pol | Second sweep of PCRs general for subtype C | Yielded sequence data |
| DP76R | CYCCTTCACCTTTCCAGA | pol | Second sweep of PCRs general for subtype C | Yielded sequence data |
| DP77F | CCCYATTTGGAAAGGACCA | pol | Second sweep of PCRs general for subtype C | Did not yield sequence data |
| DP77R | CCTTGGTACTACYTTTATGTC | pol | Second sweep of PCRs general for subtype C | Yielded sequence data |
| DP78F | GAAGGRGCAGTAGTMATACAAG | pol | Second sweep of PCRs general for subtype C | Yielded sequence data |
| DP78R | CCTGCCATCTGTTTTCCA | pol | Second sweep of PCRs general for subtype C | Yielded sequence data |
| DP79F | AAGGTAGTACCAAGRAGRAA | pol | Second sweep of PCRs general for subtype C | Yielded sequence data |
| DP79R | CTGTYTACYTGCCACACA | pol | Second sweep of PCRs general for subtype C | Yielded sequence data |
| DP7F | GTCARATAAARGARGCTCTC | pol | Second sweep of PCRs general for subtype C | Did not yield sequence data |
| DP7R | TTCCTGGCAAAYTYA | pol | Second sweep of PCRs general for subtype C | Did not yield sequence data |
| DP8F | CAGGRGCWGATGATACAGT | pol | Second sweep of PCRs general for subtype C | Did not yield sequence data |
| DP8R | AACCTCCAATTCCYCCT | pol | Second sweep of PCRs general for subtype C | Did not yield sequence data |
| DP9F | ATGATAGGRGGAATTGG | pol | Second sweep of PCRs general for subtype C | Did not yield sequence data |
| DP9R | ACTGTACCTATAGCCYT | pol | Second sweep of PCRs general for subtype C | Did not yield sequence data |
| KP79F | AAGGTAGTACCAAGRAGRAA | vif-vpr-vpu | Second sweep of PCRs general for subtype C | Did not yield sequence data |
| KP79R | CTGTYTACYTGCCACACA | vif-vpr-vpu | Second sweep of PCRs general for subtype C | Did not yield sequence data |
| KV10F | TTTKCAGAMTCTGCCAT | vif-vpr-vpu | Second sweep of PCRs general for subtype C | Yielded sequence data |
| KV10R | CTGCTTGATAGTCACACCTA | vif-vpr-vpu | Second sweep of PCRs general for subtype C | Yielded sequence data |
| KV11F | CTGCCATAAGRMAAGCCAT | vif-vpr-vpu | Second sweep of PCRs general for subtype C | Yielded sequence data |
| KV11R | GCCAARTATTGTAGAGATCC | vif-vpr-vpu | Second sweep of PCRs general for subtype C | Yielded sequence data |
| KV12F | GTGACTATCAAGCAGGACA | vif-vpr-vpu | Second sweep of PCRs general for subtype C | Yielded sequence data |
| KV12R | GGCTTTATYAATGCTGTYAG | vif-vpr-vpu | Second sweep of PCRs general for subtype C | Yielded sequence data |
| KV13F | GCAGGACATAAYAAGGTAGG | vif-vpr-vpu | Second sweep of PCRs general for subtype C | Yielded sequence data |
| KV13R | CTAGGCAGAGGTGRCTT | vif-vpr-vpu | Second sweep of PCRs general for subtype C | Yielded sequence data |
| KV14F | TAYTTGGCACTRACAGCA | vif-vpr-vpu | Second sweep of PCRs general for subtype C | Did not yield sequence data |
| KV14R | GGCTTGTTCCATCTATCYTC | vif-vpr-vpu | Second sweep of PCRs general for subtype C | Yielded sequence data |
| KV15F | CTCTGCCTAGTGTTARRAA | vif-vpr-vpu | Second sweep of PCRs general for subtype C | Yielded sequence data |
| KV15R | TCATTGWATGGYTCCCTCTG | vif-vpr-vpu | Second sweep of PCRs general for subtype C | Yielded sequence data |
| KV16F | CAAGCCYCAGAAGAYCAG | vif-vpr-vpu | Second sweep of PCRs general for subtype C | Yielded sequence data |
| KV16R | AGCTTCCTGCTTRAGYTC | vif-vpr-vpu | Second sweep of PCRs general for subtype C | Yielded sequence data |
| KV17F | GGGAACCATWYAATGAATGG | vif-vpr-vpu | Second sweep of PCRs general for subtype C | Yielded sequence data |
| KV17R | CTAGGGAARTGTCTRACAG | vif-vpr-vpu | Second sweep of PCRs general for subtype C | Yielded sequence data |
| KV18F | TAGAGGAACTYAAGCAKGA | vif-vpr-vpu | Second sweep of PCRs general for subtype C | Did not yield sequence data |
| KV18R | CTGYCCAAGTATCCCCAT | vif-vpr-vpu | Second sweep of PCRs general for subtype C | Did not yield sequence data |
| KV19F | GACAYTTYCCTAGACCATG | vif-vpr-vpu | Second sweep of PCRs general for subtype C | Did not yield sequence data |
| KV19R | GCTTCRACTCCTGYCCA | vif-vpr-vpu | Second sweep of PCRs general for subtype C | Did not yield sequence data |
| KV1F | CAGGTGCTGATTGTGTG | vif-vpr-vpu | Second sweep of PCRs general for subtype C | Did not yield sequence data |
| KV1R | AACCATCCMYTAGCTCT | vif-vpr-vpu | Second sweep of PCRs general for subtype C | Did not yield sequence data |
| KV20F | ATATGGGGATACTTGGRCRG | vif-vpr-vpu | Second sweep of PCRs general for subtype C | Yielded sequence data |
| KV20R | GWTGGCACCCAATTCTGA | vif-vpr-vpu | Second sweep of PCRs general for subtype C | Yielded sequence data |
| KV21F | AYTGCAACAACTACTGT | vif-vpr-vpu | Second sweep of PCRs general for subtype C | Did not yield sequence data |
| KV21R | CCATTTCTTGCTCTTCTYTG | vif-vpr-vpu | Second sweep of PCRs general for subtype C | Did not yield sequence data |
| KV22F | TGCCAWCATAGCAGAATAGG | vif-vpr-vpu | Second sweep of PCRs general for subtype C | Yielded sequence data |
| KV22R | AGGGCTCTARGTTAGGATC | vif-vpr-vpu | Second sweep of PCRs general for subtype C | Yielded sequence data |
| KV23F | GCAAGAAATGGAKCCART | vif-vpr-vpu | Second sweep of PCRs general for subtype C | Yielded sequence data |
| KV23R | GTTRCAAGCAGTYTTAGG | vif-vpr-vpu | Second sweep of PCRs general for subtype C | Did not yield sequence data |
| KV24F | CCTGGAAYCAYCCAGGA | vif-vpr-vpu | Second sweep of PCRs general for subtype C | Did not yield sequence data |
| KV24R | GCCTTTTKTCTGAAAGC | vif-vpr-vpu | Second sweep of PCRs general for subtype C | Did not yield sequence data |
| KV25F | GCCTAARACTGCTTGYAA | vif-vpr-vpu | Second sweep of PCRs general for subtype C | Yielded sequence data |
| KV25R | CTGCCATAGGARATGCCTA | vif-vpr-vpu | Second sweep of PCRs general for subtype C | Yielded sequence data |
| KV26F | CAGAMAAAAGGCTTAGG | vif-vpr-vpu | Second sweep of PCRs general for subtype C | Did not yield sequence data |
| KV26R | CTCRCTGYTTGGAGGA | vif-vpr-vpu | Second sweep of PCRs general for subtype C | Did not yield sequence data |
| KV27F | CGAAGCGCTCCTCMAA | vif-vpr-vpu | Second sweep of PCRs general for subtype C | Did not yield sequence data |
| KV27R | CTCCTACTCCTARTGCTRT | vif-vpr-vpu | Second sweep of PCRs general for subtype C | Did not yield sequence data |
| KV28F | GAGRGATTTAACAGCAYTAG | vif-vpr-vpu | Second sweep of PCRs general for subtype C | Did not yield sequence data |
| KV28R | AYRCTATGGTCCACACA | vif-vpr-vpu | Second sweep of PCRs general for subtype C | Did not yield sequence data |
| KV29F | TGTGTGGACCATAGYRT | vif-vpr-vpu | Second sweep of PCRs general for subtype C | Did not yield sequence data |
| KV29R | GTCTTCTGCTCTTTCYCTA | vif-vpr-vpu | Second sweep of PCRs general for subtype C | Did not yield sequence data |
| KV2F | ATGAGGATYARAACATGGA | vif-vpr-vpu | Second sweep of PCRs general for subtype C | Did not yield sequence data |
| KV2R | ACTTTTGGATKTCTRCT | vif-vpr-vpu | Second sweep of PCRs general for subtype C | Did not yield sequence data |
| KV30F | TWAGRGAAAGAGCAGAAGAC | vif-vpr-vpu | Second sweep of PCRs general for subtype C | Did not yield sequence data |
| KV30R | AAGCCTAAGAYKCCCCAT | vif-vpr-vpu | Second sweep of PCRs general for subtype C | Did not yield sequence data |
| KV31F | TGAWGGRGATACWGAGGAA | vif-vpr-vpu | Second sweep of PCRs general for subtype C | Yielded sequence data |
| KV31R | ACAAGTTYCCYACCACA | vif-vpr-vpu | Second sweep of PCRs general for subtype C | Yielded sequence data |
| KV32F | GMATCTTAGGCTTTTGGATG | vif-vpr-vpu | Second sweep of PCRs general for subtype C | Yielded sequence data |
| KV32R | AGGTACYCCATAATARACTG | vif-vpr-vpu | Second sweep of PCRs general for subtype C | Yielded sequence data |
| KV33F | TTGTAATGTGGTRGGRAAC | vif-vpr-vpu | Second sweep of PCRs general for subtype C | Did not yield sequence data |
| KV33R | TGACGCACARAATAGAGTRG | vif-vpr-vpu | Second sweep of PCRs general for subtype C | Yielded sequence data |
| KV34F | GTYTATTATGGRGTACCTGT | vif-vpr-vpu | Second sweep of PCRs general for subtype C | Did not yield sequence data |
| KV34R | TCCYTCTCATATGCTYTAGC | vif-vpr-vpu | Second sweep of PCRs general for subtype C | Yielded sequence data |
| KV35F | CTACTCTATTYTGTGCRTCA | vif-vpr-vpu | Second sweep of PCRs general for subtype C | Yielded sequence data |
| KV35R | TACACAGGCATGTGTRGC | vif-vpr-vpu | Second sweep of PCRs general for subtype C | Yielded sequence data |
| KV3F | CAAAAAGAGCTARKGGATGG | vif-vpr-vpu | Second sweep of PCRs general for subtype C | Yielded sequence data |
| KV3R | TGGGATGTGTACTTCTGAAC | vif-vpr-vpu | Second sweep of PCRs general for subtype C | Yielded sequence data |
| KV4F | GAAAGYAGAMATCCAAA | vif-vpr-vpu | Second sweep of PCRs general for subtype C | Did not yield sequence data |
| KV4R | CCTGTTTGCARMCCCCAA | vif-vpr-vpu | Second sweep of PCRs general for subtype C | Yielded sequence data |

| | | | | |
|---|---|---|---|---|
| KV5F | GAAGTACACATCCCATTAGG | vif-vpr-vpu | Second sweep of PCRs general for subtype C | Yielded sequence data |
| KV5R | GCCAATCYCTTTCTCCT | vif-vpr-vpu | Second sweep of PCRs general for subtype C | Yielded sequence data |
| KV6F | CCCATTAGGGGAKGCYAG | vif-vpr-vpu | Second sweep of PCRs general for subtype C | Yielded sequence data |
| KV6R | GAGCAYCCATKACCCAA | vif-vpr-vpu | Second sweep of PCRs general for subtype C | Did not yield sequence data |
| KV7F | GGGTTTGCAAACAGGAGA | vif-vpr-vpu | Second sweep of PCRs general for subtype C | Yielded sequence data |
| KV7R | CCAGGRTCTACYTGTGTG | vif-vpr-vpu | Second sweep of PCRs general for subtype C | Yielded sequence data |
| KV8F | GGAGATTGAGGARATAYAGC | vif-vpr-vpu | Second sweep of PCRs general for subtype C | Did not yield sequence data |
| KV8R | CTTATGGCAGAKTCTGMA | vif-vpr-vpu | Second sweep of PCRs general for subtype C | Did not yield sequence data |
| KV9F | ATCCTGGCCTGGCAGA | vif-vpr-vpu | Second sweep of PCRs general for subtype C | Yielded sequence data |
| KV9R | GGCTTKYCTTATGGCAGA | vif-vpr-vpu | Second sweep of PCRs general for subtype C | Yielded sequence data |
| 66KE10R | CATAACCAGCTGGTGCAC | env | Specific primerwalking for DRC66 | Did not yield sequence data |
| 66KE12F | CCAAAGGTCACCTTTGATCC | env | Specific primerwalking for DRC66 | Did not yield sequence data |
| 66KE13F | TTATTGTGCACCAGCTGGT | env | Specific primerwalking for DRC66 | Yielded sequence data |
| 66KE13R | TCGTGCATGRTCCTGT | env | Specific primerwalking for DRC66 | Yielded sequence data |
| 66KE14F | GAGACATTCAATGGAACAG | env | Specific primerwalking for DRC66 | Yielded sequence data |
| 66KE17F | TATCAACTCAGCTGCTGT | env | Specific primerwalking for DRC66 | Yielded sequence data |
| 66KE18F | AATGGTAGTCTAGTAGAAA | env | Specific primerwalking for DRC66 | Did not yield sequence data |
| 66KE19F | TGTACAAGACCCAACA | env | Specific primerwalking for DRC66 | Yielded sequence data |
| 66KE20F | CCAGGACAAACATTYTATGC | env | Specific primerwalking for DRC66 | Did not yield sequence data |
| 66KE24F | AAGGATGTGGCAGGGGGTAG | env | Specific primerwalking for DRC66 | Yielded sequence data |
| 66KE25F | CCCTCCCATTAAAGGAA | env | Specific primerwalking for DRC66 | Did not yield sequence data |
| 66KE28F | ATGAGACCTTCAGACCTGGA | env | Specific primerwalking for DRC66 | Yielded sequence data |
| 66KE28R | GTKGGTGCTACTCCTAGTGG | env | Specific primerwalking for DRC66 | Yielded sequence data |
| 66KE2F | GTGCATAATGTYTGGGCTA | env | Specific primerwalking for DRC66 | Did not yield sequence data |
| 66KE2R | TCCATGACTATCTCTTG | env | Specific primerwalking for DRC66 | Did not yield sequence data |
| 66KE30F | TTAAACCACTAGGAGTAGC | env | Specific primerwalking for DRC66 | Did not yield sequence data |
| 66KE30R | GAACACAGCTCCTATTC | env | Specific primerwalking for DRC66 | Did not yield sequence data |
| 66KE31R | CGCCCATAGTRCTTCC | env | Specific primerwalking for DRC66 | Yielded sequence data |
| 66KE32F | TGTTCCTTGGGTTCTTRG | env | Specific primerwalking for DRC66 | Yielded sequence data |
| 66KE32R | ATAATTGTCTAGCCTGT | env | Specific primerwalking for DRC66 | Yielded sequence data |
| 66KE33F | GCGCGGCGTCAATAACG | env | Specific primerwalking for DRC66 | Yielded sequence data |
| 66KE33R | TCTATGGCTTTCAGCAA | env | Specific primerwalking for DRC66 | Yielded sequence data |
| 66KE35F | ATAGAGGCGTAACAACA | env | Specific primerwalking for DRC66 | Did not yield sequence data |
| 66KE35R | CTTTCCATAGCCAGGACTCT | env | Specific primerwalking for DRC66 | Did not yield sequence data |
| 66KE38R | CTCCAACTAATGTTCCAGGG | env | Specific primerwalking for DRC66 | Yielded sequence data |
| 66KE39R | GCATCCAGGTCATGTTATTC | env | Specific primerwalking for DRC66 | Yielded sequence data |
| 66KE40F | GCCCTGGAACATTAGTTGG | env | Specific primerwalking for DRC66 | Yielded sequence data |
| 66KE41F | GGARTAAYATGACCTGGATG | env | Specific primerwalking for DRC66 | Did not yield sequence data |
| 66KE42F | GGGATCGAGAGATTARYAA | env | Specific primerwalking for DRC66 | Did not yield sequence data |
| 66KE42Fb | GGGATAGAGAGGTTAGCAA | env | Specific primerwalking for DRC66 | Did not yield sequence data |
| 66KE43R | AAGCCTATYAAACCCCCTAC | env | Specific primerwalking for DRC66 | Did not yield sequence data |
| 66KE45F | GAATAGAGTTAGGYAGGGAT | env | Specific primerwalking for DRC66 | Yielded sequence data |
| 66KE46R | ATCTTCCTCTGCCTCGCTCT | env | Specific primerwalking for DRC66 | Did not yield sequence data |
| 66KE47R | AGGTCGTCCCAGGCAA | env | Specific primerwalking for DRC66 | Did not yield sequence data |
| 66KE49F | CGCTTGCCTGGGACGA | env | Specific primerwalking for DRC66 | Did not yield sequence data |
| 66KE49R | CTCTGTCCCAGAATTTCCA | env | Specific primerwalking for DRC66 | Did not yield sequence data |
| 66KE49Rb | CTCTATCCCAGAAGTTCCA | env | Specific primerwalking for DRC66 | Did not yield sequence data |
| 66KE50R | CAGATATTTAAGGGCTTCC | env | Specific primerwalking for DRC66 | Yielded sequence data |
| 66KE51R | CTGACCCCAATATTGCA | env | Specific primerwalking for DRC66 | Yielded sequence data |
| 66KE53F | GGTCAGGAGCTAAAGAAGAG | env | Specific primerwalking for DRC66 | Yielded sequence data |
| 66KE54F | CTATAGCCATAGCAGTAGC | env | Specific primerwalking for DRC66 | Yielded sequence data |
| 66KE57F | AGAAGAATAAGACAGGGCCT | env | Specific primerwalking for DRC66 | Did not yield sequence data |
| 66KE58F | ATGGGGGGCAAGTGGT | env | Specific primerwalking for DRC66 | Did not yield sequence data |
| 66KE5F | ATCAAAGCTTAARGCCAT | env | Specific primerwalking for DRC66 | Did not yield sequence data |
| 66KE65R | CTTAGAGTAAATTAGCCCATCC | env | Specific primerwalking for DRC66 | Did not yield sequence data |
| 66KE9R | CTTTGGACAGGCTTGTG | env | Specific primerwalking for DRC66 | Did not yield sequence data |
| CH114F | ACCTAGAAGAATAAGACAGG | env | Specific primerwalking for DRC66 | Yielded sequence data |
| CH114R | CCATCCAACTATACTRCTYTT | env | Specific primerwalking for DRC66 | Yielded sequence data |
| CH115F | GAAAGAGTTTTGCWATAAAAT | env | Specific primerwalking for DRC66 | Did not yield sequence data |
| CH115R | CATTCTTTCTCTTACMKCAG | env | Specific primerwalking for DRC66 | Did not yield sequence data |
| CH43F | CCACAGAAATAAGAGATARG | env | Specific primerwalking for DRC66 | Did not yield sequence data |
| CH43R | TCACTAGAGTTGCTATTGCT | env | Specific primerwalking for DRC66 | Did not yield sequence data |
| CH44F | GAGAGAAGTGCATAATGT | env | Specific primerwalking for DRC66 | Yielded sequence data |
| CH44R | GACTATCTCTTGTGGGT | env | Specific primerwalking for DRC66 | Yielded sequence data |
| CH45F | ATAATCAGTCTATGGGATCA | env | Specific primerwalking for DRC66 | Did not yield sequence data |
| CH45R | CTCTACATKTTAAAGTGACA | env | Specific primerwalking for DRC66 | Did not yield sequence data |
| CH46F | CACTCTGTGTCACTTTAAC | env | Specific primerwalking for DRC66 | Did not yield sequence data |
| CH46R | TTATTTCTCCCTTCATGGTA | env | Specific primerwalking for DRC66 | Did not yield sequence data |
| CH47F | CAGTAATGGAAGTATTACCTA | env | Specific primerwalking for DRC66 | Did not yield sequence data |
| CH47R | TTATTTCTGTGGTTRCATT | env | Specific primerwalking for DRC66 | Did not yield sequence data |
| CH48F | AAAAATTGCTCTTTCAATGC | env | Specific primerwalking for DRC66 | Did not yield sequence data |
| CH48R | AGTGGTACTAYATCAAGTYT | env | Specific primerwalking for DRC66 | Did not yield sequence data |
| CH49F | ATATTAATAAATTGYAATACCTC | env | Specific primerwalking for DRC66 | Yielded sequence data |
| CH49R | ATAGGAATTGGATCAAAGGTG | env | Specific primerwalking for DRC66 | Yielded sequence data |
| CH50F | ACTCAGCTGCTGTTAAA | env | Specific primerwalking for DRC66 | Yielded sequence data |
| CH50R | ATTGTYTGTCAGATTTTSAG | env | Specific primerwalking for DRC66 | Yielded sequence data |

| | | | | |
|---|---|---|---|---|
| CH51F | CTGACARACAATGYCAAAAC | env | Specific primerwalking for DRC66 | Did not yield sequence data |
| CH51R | ATACTTTGTCTTGTATTATTGC | env | Specific primerwalking for DRC66 | Did not yield sequence data |
| CH51Rb | ATACTTTGTCTTGTATTATTGT | env | Specific primerwalking for DRC66 | Did not yield sequence data |
| CH52F | CACCTGTARAAATTGWGTG | env | Specific primerwalking for DRC66 | Did not yield sequence data |
| CH52R | TGTTTGTCCTGGTCCTATAC | env | Specific primerwalking for DRC66 | Did not yield sequence data |
| CH53F | ATACGTATAGGACCAGGA | env | Specific primerwalking for DRC66 | Yielded sequence data |
| CH53R | CTAATGTTACAATGTGCYTG | env | Specific primerwalking for DRC66 | Yielded sequence data |
| CH54F | TAGGAAAYATAAGACARGC | env | Specific primerwalking for DRC66 | Did not yield sequence data |
| CH54R | CAGGGAAGTGTTCTYYTAAT | env | Specific primerwalking for DRC66 | Did not yield sequence data |
| CH55F | GAACACTTCCCTRATAAAA | env | Specific primerwalking for DRC66 | Did not yield sequence data |
| CH55R | GTTAAARCTATGTGTTGTAAT | env | Specific primerwalking for DRC66 | Did not yield sequence data |
| CH56F | TCCCTRATAAAACAATAARAT | env | Specific primerwalking for DRC66 | Did not yield sequence data |
| CH56R | CCTCTACAATTAAARCTRTG | env | Specific primerwalking for DRC66 | Did not yield sequence data |
| CH57F | ACCTAGAAATTACAACACAT | env | Specific primerwalking for DRC66 | Did not yield sequence data |
| CH57R | TGTACTATTAAACARTYTTGA | env | Specific primerwalking for DRC66 | Did not yield sequence data |
| CH58F | GGAAATATAACATGTAAATCAA | env | Specific primerwalking for DRC66 | Yielded sequence data |
| CH58R | CCTGTACTATTTGCACC | env | Specific primerwalking for DRC66 | Yielded sequence data |
| CH59F | AGAAATTAAACCACTAGGAG | env | Specific primerwalking for DRC66 | Yielded sequence data |
| CH59R | TATTCCCACTGCTCTTTT | env | Specific primerwalking for DRC66 | Yielded sequence data |
| CH74F | AATAGGAGCTGTGTTCC | env | Specific primerwalking for DRC66 | Did not yield sequence data |
| CH74R | CAATAATTGTCTAGCCTGTA | env | Specific primerwalking for DRC66 | Did not yield sequence data |
| CH75F | TCAATAACGCTGATGGTA | env | Specific primerwalking for DRC66 | Yielded sequence data |
| CH75R | CTATGGCTTTCAGCAAAT | env | Specific primerwalking for DRC66 | Yielded sequence data |
| CH76F | CATAGAGGCGTAACAACATT | env | Specific primerwalking for DRC66 | Did not yield sequence data |
| CH76R | TTAGGTATCTTTCCATAGCC | env | Specific primerwalking for DRC66 | Did not yield sequence data |
| CH89F | ATAAYATGACCTGGATGC | env | Specific primerwalking for DRC66 | Yielded sequence data |
| CH89R | TTTTGCGATTYTTCAAGYAA | env | Specific primerwalking for DRC66 | Yielded sequence data |
| CH90F | RATTTACTAGMATTGGACA | env | Specific primerwalking for DRC66 | Did not yield sequence data |
| CH90R | TATATACCAYAGCCAKTTTG | env | Specific primerwalking for DRC66 | Did not yield sequence data |
| CH90Rb | TATATACCACAGCCARTTTG | env | Specific primerwalking for DRC66 | Did not yield sequence data |
| CH91F | TTGGTTTRACATAACAAAMTG | env | Specific primerwalking for DRC66 | Did not yield sequence data |
| CH91Fb | TTGGTTTRACATARCAAAYTG | env | Specific primerwalking for DRC66 | Did not yield sequence data |
| CH91R | AAAGCACARCAAAAACTA | env | Specific primerwalking for DRC66 | Did not yield sequence data |
| CH92F | AGTGAATAGAGTTAGGYA | env | Specific primerwalking for DRC66 | Did not yield sequence data |
| CH92R | ACCTTCTTCTTCGATTCYT | env | Specific primerwalking for DRC66 | Did not yield sequence data |
| CH93F | CAGACCCTTATCCCAAMC | env | Specific primerwalking for DRC66 | Did not yield sequence data |
| CH93R | CGCTCACTAATCGAATG | env | Specific primerwalking for DRC66 | Did not yield sequence data |
| CH94F | GATTCTTAGCGCTTGC | env | Specific primerwalking for DRC66 | Did not yield sequence data |
| CH94R | CAATCAATATGAAGTYTCTC | env | Specific primerwalking for DRC66 | Did not yield sequence data |
| CH95F | CKCTACCACCRATTGAGA | env | Specific primerwalking for DRC66 | Did not yield sequence data |
| CH95R | AGATATTTAAGGGCTTCCCA | env | Specific primerwalking for DRC66 | Did not yield sequence data |
| CH96F | TTGCAATACATCAARAYTG | env | Specific primerwalking for DRC66 | Did not yield sequence data |
| CH96R | TTTGTCTTATTCTGCATKGG | env | Specific primerwalking for DRC66 | Did not yield sequence data |
| CH97F | TTATARACTTGATRTAGTACC | env | Specific primerwalking for DRC66 | Yielded sequence data |
| CH97R | ACAGGCTTGTGTAACG | env | Specific primerwalking for DRC66 | Yielded sequence data |
| TI10F | TCTGAAAATCTRACARACAATGYCA | env | Specific primerwalking for DRC66 | Did not yield sequence data |
| TI10R | TGTATTGTTGYYGGGTCTTGT | env | Specific primerwalking for DRC66 | Did not yield sequence data |
| TI11F | TGCAACAGGAGAGATAATAGGAG | env | Specific primerwalking for DRC66 | Did not yield sequence data |
| TI11R | CCCGTTGTAAAGTTCYRTTCC | env | Specific primerwalking for DRC66 | Did not yield sequence data |
| TI12F | AAATTAGCARARCACTTCCCT | env | Specific primerwalking for DRC66 | Did not yield sequence data |
| TI12R | TAATYTCTAGGTCCCCTCCYG | env | Specific primerwalking for DRC66 | Did not yield sequence data |
| TI13F | ATCGAYCCTCRGGAGGGG | env | Specific primerwalking for DRC66 | Yielded sequence data |
| TI13R | TTGCAATAGAAAAATTCTCCTCTAC | env | Specific primerwalking for DRC66 | Yielded sequence data |
| TI14F | TTGCAATACATCAARRCTGTTT | env | Specific primerwalking for DRC66 | Did not yield sequence data |
| TI14R | CTGCATGGGAGTGTGATKKT | env | Specific primerwalking for DRC66 | Did not yield sequence data |
| TI15F | GAAAGCCATAGAGGCGTAACA | env | Specific primerwalking for DRC66 | Did not yield sequence data |
| TI15R | TCTTTCCATAGCCAGGACTCT | env | Specific primerwalking for DRC66 | Did not yield sequence data |
| TI16F | CAGTGGGATAGAGAGGTTAGCA | env | Specific primerwalking for DRC66 | Did not yield sequence data |
| TI16R | TTTCYTCCTGCTGGTTYTGC | env | Specific primerwalking for DRC66 | Did not yield sequence data |
| TI7F | ATAATCAGTCTATGGGATCAAAGC | env | Specific primerwalking for DRC66 | Yielded sequence data |
| TI7R | ATTCTARAGTGACACAGAGTGKGG | env | Specific primerwalking for DRC66 | Yielded sequence data |
| TI8F | GGATCAAAGCTTAAAGCCATGTGT | env | Specific primerwalking for DRC66 | Did not yield sequence data |
| TI8R | CGAGCTAACATTAACCAAGGTAACA | env | Specific primerwalking for DRC66 | Did not yield sequence data |
| TI9F | CAATGCAACYACAGAAMTAARAGAT | env | Specific primerwalking for DRC66 | Did not yield sequence data |
| TI9R | CCTCTCACTCTCATTCCCC | env | Specific primerwalking for DRC66 | Did not yield sequence data |
| 66CH12F | GACAAAGGAAARGTCAGTC | gag | Specific primerwalking for DRC66 | Did not yield sequence data |
| 66CH12R | CCGCATTYAAAGTTCTAG | gag | Specific primerwalking for DRC66 | Did not yield sequence data |
| 66CH77R | TGCACTATAGGATARTTYTG | gag | Specific primerwalking for DRC66 | Did not yield sequence data |
| 66CH85F | GGTCAGTCAAAATTATCCTA | gag | Specific primerwalking for DRC66 | Did not yield sequence data |
| 66CH85R | TCTATTACTTTTACCCAYGC | gag | Specific primerwalking for DRC66 | Did not yield sequence data |
| 66CH86F | GTACACCAGSCCWTATCAC | gag | Specific primerwalking for DRC66 | Did not yield sequence data |
| 66KG11F | GCTTTAGAAAAGATGGAAGA | gag | Specific primerwalking for DRC66 | Did not yield sequence data |
| 66KG15R | TTGTGGGATGRCTCCT | gag | Specific primerwalking for DRC66 | Did not yield sequence data |
| 66KG18F | TAAACACAGTGGGGGGAYA | gag | Specific primerwalking for DRC66 | Did not yield sequence data |
| 66KG21R | GRGTTGCTTGTCAYCCAT | gag | Specific primerwalking for DRC66 | Did not yield sequence data |
| 66KG22F | AGGAACTACTAGTAACCTTC | gag | Specific primerwalking for DRC66 | Yielded sequence data |

| | | | | |
|---|---|---|---|---|
| 66KG25F | CAAGGGCCAAAAGAACYCT | gag | Specific primerwalking for DRC66 | Yielded sequence data |
| 66KG25R | GTGTGACTTGCTCAGCTCT | gag | Specific primerwalking for DRC66 | Yielded sequence data |
| 66KG27F | GCTGAGCAAGTCACACAGG | gag | Specific primerwalking for DRC66 | Yielded sequence data |
| 66KG2F | GGGGGGAAATTAGATGCATG | gag | Specific primerwalking for DRC66 | Yielded sequence data |
| 66KG31F | CTGAGGCGATGAGCCAA | gag | Specific primerwalking for DRC66 | Did not yield sequence data |
| 66KG34R | TGAGAAGGCCAAAGTTTC | gag | Specific primerwalking for DRC66 | Yielded sequence data |
| 66KG38R | TGGTCTTTCGGCTCCT | gag | Specific primerwalking for DRC66 | Yielded sequence data |
| 66KG3R | GGCCAGGGTTAAGTGCAA | gag | Specific primerwalking for DRC66 | Yielded sequence data |
| 66KG5F | GGGAGCTGGAAAGATTTGC | gag | Specific primerwalking for DRC66 | Yielded sequence data |
| 66KG5R | GAAGAGCTGGTTGTAGCT | gag | Specific primerwalking for DRC66 | Yielded sequence data |
| 66KG6F | GAAACAACAGAAGGCTGT | gag | Specific primerwalking for DRC66 | Yielded sequence data |
| 66KG7F | AGCTACAAYCAGCTYTTC | gag | Specific primerwalking for DRC66 | Did not yield sequence data |
| CH10F | CATCAAAGGATAGCGGTAC | gag | Specific primerwalking for DRC66 | Yielded sequence data |
| CH10R | CCTGCTGTGYTTTTTGC | gag | Specific primerwalking for DRC66 | Yielded sequence data |
| CH11F | AAARCACAGCAGGCARA | gag | Specific primerwalking for DRC66 | Did not yield sequence data |
| CH11R | GGTGTACCATTTGCCCYT | gag | Specific primerwalking for DRC66 | Did not yield sequence data |
| CH12F | GACAAAGGAAARGTCAGYC | gag | Specific primerwalking for DRC66 | Did not yield sequence data |
| CH12R | CCGCATTCAARGTTCTRG | gag | Specific primerwalking for DRC66 | Did not yield sequence data |
| CH13F | CTCCAAGGGCAAATGGTA | gag | Specific primerwalking for DRC66 | Did not yield sequence data |
| CH13R | TGGGTATTACCTCTGGGYT | gag | Specific primerwalking for DRC66 | Did not yield sequence data |
| CH14F | CGKGGGTAAAAGTARTAGAG | gag | Specific primerwalking for DRC66 | Yielded sequence data |
| CH14R | TGGGATGRCTCCTTCTGA | gag | Specific primerwalking for DRC66 | Yielded sequence data |
| CH15F | CCTATTCCAGTGGGAGACA | gag | Specific primerwalking for DRC66 | Did not yield sequence data |
| CH15R | GGCCCTTGTTTTATGTCCA | gag | Specific primerwalking for DRC66 | Did not yield sequence data |
| CH16F | TGGTYCAAAATGCGAACC | gag | Specific primerwalking for DRC66 | Yielded sequence data |
| CH16R | CCCTGACATGCTGTCATC | gag | Specific primerwalking for DRC66 | Yielded sequence data |
| CH17F | ACAGCATGTCAGGGAGT | gag | Specific primerwalking for DRC66 | Yielded sequence data |
| CH17R | GTTTGCTTGGCTCATCG | gag | Specific primerwalking for DRC66 | Yielded sequence data |
| CH18F | AGGCGATGAGCCAAGC | gag | Specific primerwalking for DRC66 | Yielded sequence data |
| CH18R | CTTCCTTGCCACAGTTG | gag | Specific primerwalking for DRC66 | Yielded sequence data |
| CH19F | TCAACTGTGGCAAGGAAG | gag | Specific primerwalking for DRC66 | Did not yield sequence data |
| CH19R | GGTGTCCTTCTTTTCCACA | gag | Specific primerwalking for DRC66 | Did not yield sequence data |
| CH77F | AGAAAAGATGGAAGAAGCA | gag | Specific primerwalking for DRC66 | Did not yield sequence data |
| CH77R | TGCACTATAGGRTAATTTTG | gag | Specific primerwalking for DRC66 | Did not yield sequence data |
| CH85F | GGTCAGTCAAAATTAYCCYA | gag | Specific primerwalking for DRC66 | Did not yield sequence data |
| CH85R | TCTAYTACTTTTACCCAKGC | gag | Specific primerwalking for DRC66 | Did not yield sequence data |
| CH86F | GTACACCAGSCCMTRTCAC | gag | Specific primerwalking for DRC66 | Did not yield sequence data |
| CH86R | GAGTATTACTTCTGAGTTGA | gag | Specific primerwalking for DRC66 | Did not yield sequence data |
| CH87F | ATAAAGCAAGRGTTTTGG | gag | Specific primerwalking for DRC66 | Did not yield sequence data |
| CH87R | ATAATTCTTTTAGGGCCTTT | gag | Specific primerwalking for DRC66 | Did not yield sequence data |
| CH88F | TTAAAGGCCCTAAAAGAAT | gag | Specific primerwalking for DRC66 | Did not yield sequence data |
| CH88R | CTGCAATTTCTGGCTAT | gag | Specific primerwalking for DRC66 | Did not yield sequence data |
| CH98F | TATAAAAGATGGATAATCCTG | gag | Specific primerwalking for DRC66 | Did not yield sequence data |
| CH98R | TTTGGCCCTTGTTTTATG | gag | Specific primerwalking for DRC66 | Did not yield sequence data |
| CH9F | CYAGCTACAACCAGCTCT | gag | Specific primerwalking for DRC66 | Yielded sequence data |
| CH9R | GTACCGCTATCCTTTGATG | gag | Specific primerwalking for DRC66 | Yielded sequence data |
| 66CH25R | ACCCCATCCAAAGGAATGA | pol | Specific primerwalking for DRC66 | Did not yield sequence data |
| 66CH28F | TTGGCAGAGAAYAGRGAA | pol | Specific primerwalking for DRC66 | Did not yield sequence data |
| 66CH31F | GACTCATCAAAAGACTTAA | pol | Specific primerwalking for DRC66 | Did not yield sequence data |
| 66CH41R | ATTGCTGCCATTGTCTG | pol | Specific primerwalking for DRC66 | Did not yield sequence data |
| 66CH83R | TTTGTAAAATTTGTTTTTGT | pol | Specific primerwalking for DRC66 | Did not yield sequence data |
| 66DP35F | TAGGGGGAYCAAAGCA | pol | Specific primerwalking for DRC66 | Yielded sequence data |
| 66KP11F | GGACCTACACCTGTCAAC | pol | Specific primerwalking for DRC66 | Did not yield sequence data |
| 66KP16R | ATTTTCTCCAYTTRGTACTG | pol | Specific primerwalking for DRC66 | Yielded sequence data |
| 66KP17F | AATACTCTAGTATTTGC | pol | Specific primerwalking for DRC66 | Yielded sequence data |
| 66KP20F | TCAAGACTTCTGGGAAGTCC | pol | Specific primerwalking for DRC66 | Did not yield sequence data |
| 66KP20R | CATCTAGTACTGTTACTGA | pol | Specific primerwalking for DRC66 | Yielded sequence data |
| 66KP21R | GCAGTATAYTTCCTGAARCC | pol | Specific primerwalking for DRC66 | Yielded sequence data |
| 66KP24F | AGATGAAGGYTTCAGGA | pol | Specific primerwalking for DRC66 | Did not yield sequence data |
| 66KP25R | TGACGGTGATCCTTTCC | pol | Specific primerwalking for DRC66 | Did not yield sequence data |
| 66KP29R | TCARGATGGAGTTCATACC | pol | Specific primerwalking for DRC66 | Did not yield sequence data |
| 66KP2F | GGGAATTTCCTTCAGARYAG | pol | Specific primerwalking for DRC66 | Did not yield sequence data |
| 66KP2Fb | GGGAATTTCCTTCAGAGCAG | pol | Specific primerwalking for DRC66 | Did not yield sequence data |
| 66KP2R | CTTTCGGCTCCTGYTTC | pol | Specific primerwalking for DRC66 | Did not yield sequence data |
| 66KP31R | TCATTGACAGCCCAGCT | pol | Specific primerwalking for DRC66 | Did not yield sequence data |
| 66KP32R | CCTGGATAAATCTGACT | pol | Specific primerwalking for DRC66 | Yielded sequence data |
| 66KP33F | TAAAYTGGGTAAGTCAG | pol | Specific primerwalking for DRC66 | Did not yield sequence data |
| 66KP33FB | TAAATTGGGYAAGTCAG | pol | Specific primerwalking for DRC66 | Did not yield sequence data |
| 66KP33R | TGCTTTGRTCCCCCTA | pol | Specific primerwalking for DRC66 | Did not yield sequence data |
| 66KP34F | AAGTCAGATTTATCCAGGG | pol | Specific primerwalking for DRC66 | Did not yield sequence data |
| 66KP35R | GTACTGGTTCTCTGCCAA | pol | Specific primerwalking for DRC66 | Yielded sequence data |
| 66KP37R | GCTATTAAGTCTTTTGATGA | pol | Specific primerwalking for DRC66 | Did not yield sequence data |
| 66KP39F | CYCATCAAAAGACTTAATAG | pol | Specific primerwalking for DRC66 | Did not yield sequence data |
| 66KP43R | TCCACCATGTCTCCCATA | pol | Specific primerwalking for DRC66 | Did not yield sequence data |
| 66KP47F | GTYAATACCCCTCCTYT | pol | Specific primerwalking for DRC66 | Yielded sequence data |
| 66KP56R | ATAGTACTCTCCTGATTCC | pol | Specific primerwalking for DRC66 | Yielded sequence data |

| | | | | |
|---|---|---|---|---|
| 66KP58F | GGAATCAGGAGAGTACTAT | pol | Specific primerwalking for DRC66 | Did not yield sequence data |
| 66KP59F | GGCTCAAGAAGAACATG | pol | Specific primerwalking for DRC66 | Yielded sequence data |
| 66KP59R | TTTGCTACTATGGGTGGCAG | pol | Specific primerwalking for DRC66 | Did not yield sequence data |
| 66KP5F | CCCTYAAATCACTCTTTGG | pol | Specific primerwalking for DRC66 | Did not yield sequence data |
| 66KP64R | CTGCTTCCATGTAGCCACT | pol | Specific primerwalking for DRC66 | Did not yield sequence data |
| 66KP68R | TACCTGCCCACCARCAG | pol | Specific primerwalking for DRC66 | Did not yield sequence data |
| 66KP6F | GCGACCCCTTGTCWCA | pol | Specific primerwalking for DRC66 | Did not yield sequence data |
| 66KP71F | TTGGAATTCCCTACAATYC | pol | Specific primerwalking for DRC66 | Yielded sequence data |
| 66KP75R | GTCTCTGTAATAAACCCAAA | pol | Specific primerwalking for DRC66 | Did not yield sequence data |
| 66KP77F | CCCAATTTGGAAAGGACCA | pol | Specific primerwalking for DRC66 | Yielded sequence data |
| 66KP77R | TCTTGRCACTACTTTTATTTC | pol | Specific primerwalking for DRC66 | Yielded sequence data |
| 66KP7R | TTCCTGGTAAATTTA | pol | Specific primerwalking for DRC66 | Did not yield sequence data |
| CH100F | AAGGATTATGGAAAACAGAT | pol | Specific primerwalking for DRC66 | Did not yield sequence data |
| CH100R | CATACATATGRTGTTTCACT | pol | Specific primerwalking for DRC66 | Did not yield sequence data |
| CH104F | TAGTATCACTGACTGAAGAA | pol | Specific primerwalking for DRC66 | Yielded sequence data |
| CH104R | AGTCATAATATACCCCATGT | pol | Specific primerwalking for DRC66 | Yielded sequence data |
| CH111F | CCTTAACTTCCCTYAAATC | pol | Specific primerwalking for DRC66 | Yielded sequence data |
| CH111R | CCTGTATCTAAGAGAGCYT | pol | Specific primerwalking for DRC66 | Yielded sequence data |
| CH112F | CCCTTGTCACAATAARARTA | pol | Specific primerwalking for DRC66 | Did not yield sequence data |
| CH112R | TTCTAATACTGTATCATCTGC | pol | Specific primerwalking for DRC66 | Did not yield sequence data |
| CH113F | TCAAAGTAAGRCARTATGAT | pol | Specific primerwalking for DRC66 | Did not yield sequence data |
| CH113R | ATTATGTTGACAGGTGTAG | pol | Specific primerwalking for DRC66 | Did not yield sequence data |
| CH20F | TTGGGCCTGAAAATCCA | pol | Specific primerwalking for DRC66 | Yielded sequence data |
| CH20R | CTCCACTTAGTACTGTC | pol | Specific primerwalking for DRC66 | Yielded sequence data |
| CH21F | TTCAAGACTTCTGGGAAGTC | pol | Specific primerwalking for DRC66 | Yielded sequence data |
| CH21R | TCCCCCACATCTAGTACTG | pol | Specific primerwalking for DRC66 | Yielded sequence data |
| CH23F | ACAATGAGAYACCAGGGAT | pol | Specific primerwalking for DRC66 | Yielded sequence data |
| CH23R | GGAATATTGACGGTGATCCT | pol | Specific primerwalking for DRC66 | Yielded sequence data |
| CH24F | TGGAAAGGATCACCGTCA | pol | Specific primerwalking for DRC66 | Yielded sequence data |
| CH24R | CTGGATTTTKTGCYCTA | pol | Specific primerwalking for DRC66 | Yielded sequence data |
| CH25F | GAGGAGCTRAGARAACATC | pol | Specific primerwalking for DRC66 | Did not yield sequence data |
| CH25R | ACCCCATCCAAAGAAATGG | pol | Specific primerwalking for DRC66 | Did not yield sequence data |
| CH26F | CCAGAYAAGAARCATCAGA | pol | Specific primerwalking for DRC66 | Yielded sequence data |
| CH26R | GTATRGGCTGTACTATCCAT | pol | Specific primerwalking for DRC66 | Yielded sequence data |
| CH27F | GGTATGAACTCCATCCTGA | pol | Specific primerwalking for DRC66 | Did not yield sequence data |
| CH27R | ATTGACAGCCCAGCTGT | pol | Specific primerwalking for DRC66 | Did not yield sequence data |
| CH28F | TTGGCAGAGAACCAGTAC | pol | Specific primerwalking for DRC66 | Did not yield sequence data |
| CH28R | GCTATTAAGTCTTTTGATGA | pol | Specific primerwalking for DRC66 | Did not yield sequence data |
| CH29F | CTAARAGAACCAGTACATGG | pol | Specific primerwalking for DRC66 | Did not yield sequence data |
| CH29R | TGATATGTCCAYTGGTCMTG | pol | Specific primerwalking for DRC66 | Did not yield sequence data |
| CH30F | GGCTCAAGAAGAACATG | pol | Specific primerwalking for DRC66 | Yielded sequence data |
| CH30R | GCTACTATGGGTGGCAGA | pol | Specific primerwalking for DRC66 | Yielded sequence data |
| CH31F | GACCCATCAAAAGACTTGG | pol | Specific primerwalking for DRC66 | Did not yield sequence data |
| CH31R | CTTYCCTGTTTTCAGRT | pol | Specific primerwalking for DRC66 | Did not yield sequence data |
| CH32F | GGCAKGACCAATGGACA | pol | Specific primerwalking for DRC66 | Did not yield sequence data |
| CH32R | GGCAGTCCTCATTYTTGC | pol | Specific primerwalking for DRC66 | Did not yield sequence data |
| CH33F | CTGAAAACAGGRAAGTATGC | pol | Specific primerwalking for DRC66 | Did not yield sequence data |
| CH33R | CACTGCCTCTGYTAAYTGT | pol | Specific primerwalking for DRC66 | Did not yield sequence data |
| CH34F | GAGGACTGCCCACACT | pol | Specific primerwalking for DRC66 | Did not yield sequence data |
| CH34R | TAGGAGTCTTTCCCCAT | pol | Specific primerwalking for DRC66 | Did not yield sequence data |
| CH35F | TATGGGGAAAGACTCCT | pol | Specific primerwalking for DRC66 | Yielded sequence data |
| CH35R | GTGGCTTGCCARTAGTC | pol | Specific primerwalking for DRC66 | Yielded sequence data |
| CH36F | GTCAATACCCCTCCTCTAG | pol | Specific primerwalking for DRC66 | Yielded sequence data |
| CH36R | CTATTAGCTGCTCCATCTAC | pol | Specific primerwalking for DRC66 | Yielded sequence data |
| CH37F | GAACCCATAAYAGGAGCAGA | pol | Specific primerwalking for DRC66 | Yielded sequence data |
| CH37R | GTAACATACCCTGCTTTTCC | pol | Specific primerwalking for DRC66 | Yielded sequence data |
| CH38F | GCAGGGTATGTTACTGAYA | pol | Specific primerwalking for DRC66 | Yielded sequence data |
| CH38R | CAAAGCCAGCTGAATTGC | pol | Specific primerwalking for DRC66 | Yielded sequence data |
| CH39F | CAAGCACAACCAGATAAGAG | pol | Specific primerwalking for DRC66 | Yielded sequence data |
| CH39R | TGGTACCCATGACAGGTAG | pol | Specific primerwalking for DRC66 | Yielded sequence data |
| CH40F | TAGTCCAGGGATATGGCAA | pol | Specific primerwalking for DRC66 | Did not yield sequence data |
| CH40R | CCATGTAGCCACTGGCTA | pol | Specific primerwalking for DRC66 | Did not yield sequence data |
| CH41F | ACAGGACAGGAAACAGT | pol | Specific primerwalking for DRC66 | Did not yield sequence data |
| CH41R | ATTRCTRCCATTGTCTG | pol | Specific primerwalking for DRC66 | Did not yield sequence data |
| CH42F | AGGAAGATGGCCAGTCA | pol | Specific primerwalking for DRC66 | Did not yield sequence data |
| CH42R | ACCAACAGGCTGCCTT | pol | Specific primerwalking for DRC66 | Did not yield sequence data |
| CH60F | TGATACAGTATTAGAAGAAA | pol | Specific primerwalking for DRC66 | Yielded sequence data |
| CH60R | ATAAAACCTCCAATTCCYC | pol | Specific primerwalking for DRC66 | Yielded sequence data |
| CH61F | GAAAATGGAARCCAAAAATGA | pol | Specific primerwalking for DRC66 | Yielded sequence data |
| CH61R | GATATTTGATCATAYTGTCTTA | pol | Specific primerwalking for DRC66 | Yielded sequence data |
| CH62F | ACCAAAAATGATAGGRGGA | pol | Specific primerwalking for DRC66 | Did not yield sequence data |
| CH62R | TACTGTACCTATAGCYYT | pol | Specific primerwalking for DRC66 | Did not yield sequence data |
| CH63F | TACACCTGTCAACATAATTG | pol | Specific primerwalking for DRC66 | Did not yield sequence data |
| CH63R | ATTTTACTGGYACAGTTTCA | pol | Specific primerwalking for DRC66 | Did not yield sequence data |
| CH64F | CTAAATTTTCCMATTAGYCC | pol | Specific primerwalking for DRC66 | Did not yield sequence data |
| CH64R | TTTTCTTCTGTCAATGGC | pol | Specific primerwalking for DRC66 | Did not yield sequence data |

| | | | | |
|---|---|---|---|---|
| CH65F | AAGCATTAATGGAAATTTGT | pol | Specific primerwalking for DRC66 | Yielded sequence data |
| CH65R | AGTRTTRTATGGATTTTCAG | pol | Specific primerwalking for DRC66 | Yielded sequence data |
| CH78F | CAATACCCCAGTATTTGC | pol | Specific primerwalking for DRC66 | Did not yield sequence data |
| CH78R | TTGAGTTCTCTGAAATCTAC | pol | Specific primerwalking for DRC66 | Yielded sequence data |
| CH79F | AGTAGCATGACAAAAATCT | pol | Specific primerwalking for DRC66 | Did not yield sequence data |
| CH79R | TCCTACATACAAGTCATCC | pol | Specific primerwalking for DRC66 | Did not yield sequence data |
| CH80F | TTGTATACACTTAGAAGGGA | pol | Specific primerwalking for DRC66 | Did not yield sequence data |
| CH80R | TAACTTCTGCTTCCATGTAG | pol | Specific primerwalking for DRC66 | Did not yield sequence data |
| CH81F | TTAAAGAAAATCATAGGGCA | pol | Specific primerwalking for DRC66 | Did not yield sequence data |
| CH81R | AAATTGTGRATGAAWACTGC | pol | Specific primerwalking for DRC66 | Did not yield sequence data |
| CH82F | TAAGACAGCAKTACAAATG | pol | Specific primerwalking for DRC66 | Did not yield sequence data |
| CH82R | CTATTATTCTTTCCCCTGC | pol | Specific primerwalking for DRC66 | Did not yield sequence data |
| CH83F | GCAGGGGAAAGAATRATAG | pol | Specific primerwalking for DRC66 | Did not yield sequence data |
| CH83R | TTTTTRTAATTTGYTTTTGT | pol | Specific primerwalking for DRC66 | Did not yield sequence data |
| CH84F | ATAGCAACAGACATACAAAC | pol | Specific primerwalking for DRC66 | Yielded sequence data |
| CH84R | CTCTGCTGTCYCTGTAATA | pol | Specific primerwalking for DRC66 | Yielded sequence data |
| CH99F | CAGAGGCWGTGCARA | pol | Specific primerwalking for DRC66 | Did not yield sequence data |
| CH99R | TATTTCTTTTTGGATGGGC | pol | Specific primerwalking for DRC66 | Did not yield sequence data |
| TH10F | TGAAAACAGGRAARTATGCAARAAT | pol | Specific primerwalking for DRC66 | Yielded sequence data |
| TH10R | CACTGCCTCTGYTAAYTGTTTT | pol | Specific primerwalking for DRC66 | Yielded sequence data |
| TH11F | TGCAARAATGAGGACTGCCCAC | pol | Specific primerwalking for DRC66 | Did not yield sequence data |
| TH11R | CCATATTACTATGCTTTCCRTRGCT | pol | Specific primerwalking for DRC66 | Did not yield sequence data |
| TH12F | AGTTAACAGAGGCWGTRCARA | pol | Specific primerwalking for DRC66 | Yielded sequence data |
| TH12R | TTTCTTTTTGGATGGGCAGTTT | pol | Specific primerwalking for DRC66 | Yielded sequence data |
| TH13F | AAGAAAATCATAGGGCAGGTAAGA | pol | Specific primerwalking for DRC66 | Yielded sequence data |
| TH13R | TGAATACTGCCATTTGTACTGCTG | pol | Specific primerwalking for DRC66 | Yielded sequence data |
| TH14F | CTGAGCAYCTTAARACAGCAG | pol | Specific primerwalking for DRC66 | Did not yield sequence data |
| TH14R | TCTATTATTCTTTCCCCTGCACTRT | pol | Specific primerwalking for DRC66 | Did not yield sequence data |
| TH1F | ATCACTCTTTGGCAGCGACC | pol | Specific primerwalking for DRC66 | Did not yield sequence data |
| TH1R | GCTCCTGTRTCTAAGAGAGCC | pol | Specific primerwalking for DRC66 | Did not yield sequence data |
| TH2F | GGGGGYCAAATAAARGARGC | pol | Specific primerwalking for DRC66 | Did not yield sequence data |
| TH2R | TTTTTGGTTTCCATTTTCCTGGTA | pol | Specific primerwalking for DRC66 | Did not yield sequence data |
| TH3F | AGGACCTACACCTGTCAACAT | pol | Specific primerwalking for DRC66 | Did not yield sequence data |
| TH3R | ACTGGTAYAGTYTCAATRGGA | pol | Specific primerwalking for DRC66 | Did not yield sequence data |
| TH4F | GCTTGGATGCACAYTAAATTTTCC | pol | Specific primerwalking for DRC66 | Yielded sequence data |
| TH4R | GTTTAACYTTTGGGCCATCCA | pol | Specific primerwalking for DRC66 | Yielded sequence data |
| TH5F | TGGATGACTTGTATGTAGGATCTGA | pol | Specific primerwalking for DRC66 | Did not yield sequence data |
| TH5R | GGTGTGGTAAATCCCCACYTYA | pol | Specific primerwalking for DRC66 | Did not yield sequence data |
| TH6F | TAAAGTGGGGATTTACCACWCCA | pol | Specific primerwalking for DRC66 | Yielded sequence data |
| TH6R | GTTCATACCCCATCCAAAGGA | pol | Specific primerwalking for DRC66 | Yielded sequence data |
| TH7F | GGGGTATGAACTCCATCCTGATA | pol | Specific primerwalking for DRC66 | Did not yield sequence data |
| TH7R | TGTATATCATTGACAGCCCAGC | pol | Specific primerwalking for DRC66 | Did not yield sequence data |
| TH8F | ATAGCTGAAATACAGAAACARGGG | pol | Specific primerwalking for DRC66 | Did not yield sequence data |
| TH8R | CAGATTTTTGAATKGTTCYTGGT | pol | Specific primerwalking for DRC66 | Did not yield sequence data |
| TH9F | GAAACARGGGCAKGACCARTGGA | pol | Specific primerwalking for DRC66 | Did not yield sequence data |
| TH9R | ATTAGTGTGGGCAGTCCTCATTT | pol | Specific primerwalking for DRC66 | Did not yield sequence data |
| 66CH103R | GATGATCCTYACTGCTT | vif-vpr-vpu | Specific primerwalking for DRC66 | Did not yield sequence data |
| 66HY10R | CACRTRAATTAGTTGGTCT | vif-vpr-vpu | Specific primerwalking for DRC66 | Did not yield sequence data |
| 66HY12R | GTCACACCTAGGRCTAACTC | vif-vpr-vpu | Specific primerwalking for DRC66 | Did not yield sequence data |
| 66HY16F | AAARGAYAAAGTCACCT | vif-vpr-vpu | Specific primerwalking for DRC66 | Did not yield sequence data |
| 66HY16R | CTCCCTCTGTGGCCCC | vif-vpr-vpu | Specific primerwalking for DRC66 | Did not yield sequence data |
| 66HY19F | GCCATTTAATGAATGGACTC | vif-vpr-vpu | Specific primerwalking for DRC66 | Did not yield sequence data |
| 66HY20F | GGAGCTTAAGCATGAAGCT | vif-vpr-vpu | Specific primerwalking for DRC66 | Did not yield sequence data |
| 66HY21R | TYATAGCYTCAACTCCT | vif-vpr-vpu | Specific primerwalking for DRC66 | Did not yield sequence data |
| 66HY23F | GAGTTGAAGCTATRATA | vif-vpr-vpu | Specific primerwalking for DRC66 | Did not yield sequence data |
| 66HY24F | TYTRCAACAACTACTGT | vif-vpr-vpu | Specific primerwalking for DRC66 | Did not yield sequence data |
| 66HY2R | CTATAAAMCCATCCATTAGC | vif-vpr-vpu | Specific primerwalking for DRC66 | Did not yield sequence data |
| 66HY36R | CTTCACTCTCATTRCCACTG | vif-vpr-vpu | Specific primerwalking for DRC66 | Did not yield sequence data |
| 66HY38F | GTGAAGGGGATACAGAGG | vif-vpr-vpu | Specific primerwalking for DRC66 | Did not yield sequence data |
| 66HY40R | TTTTGCTTCTCTCCACACAG | vif-vpr-vpu | Specific primerwalking for DRC66 | Did not yield sequence data |
| 66HY42F | GGAGAGAAGCAAAAACTA | vif-vpr-vpu | Specific primerwalking for DRC66 | Yielded sequence data |
| 66HY42R | AGCCCARACATTATGCACT | vif-vpr-vpu | Specific primerwalking for DRC66 | Yielded sequence data |
| 66HY4F | CAAAAAGAGCTAATGGATGG | vif-vpr-vpu | Specific primerwalking for DRC66 | Did not yield sequence data |
| 66HY6F | TGAAARCACACATCCAA | vif-vpr-vpu | Specific primerwalking for DRC66 | Did not yield sequence data |
| 66KP79F | AAAGTAGTGCCAAGAAGGAA | vif-vpr-vpu | Specific primerwalking for DRC66 | Did not yield sequence data |
| 66KV10F | TTTTCAGAATCTGCCAT | vif-vpr-vpu | Specific primerwalking for DRC66 | Did not yield sequence data |
| 66KV11F | CTGCCATAAGGAAAGCCAT | vif-vpr-vpu | Specific primerwalking for DRC66 | Yielded sequence data |
| 66KV16F | CAAGCCCCAGAAGACCAG | vif-vpr-vpu | Specific primerwalking for DRC66 | Yielded sequence data |
| 66KV17F | GGGAGCCATTTAATGAATGG | vif-vpr-vpu | Specific primerwalking for DRC66 | Yielded sequence data |
| 66KV18F | TAGAGGAGYTTAAGCATGA | vif-vpr-vpu | Specific primerwalking for DRC66 | Did not yield sequence data |
| 66KV18R | CTGYCCAAGTATCTCCAT | vif-vpr-vpu | Specific primerwalking for DRC66 | Did not yield sequence data |
| 66KV19R | GCTTCAACTCCTGCCCA | vif-vpr-vpu | Specific primerwalking for DRC66 | Did not yield sequence data |
| 66KV1R | AACCATCCATTAGCTCT | vif-vpr-vpu | Specific primerwalking for DRC66 | Did not yield sequence data |
| 66KV24F | CCTGGAACCATCCGGGA | vif-vpr-vpu | Specific primerwalking for DRC66 | Did not yield sequence data |
| 66KV24R | GCCTTTGTTCAGAAAGC | vif-vpr-vpu | Specific primerwalking for DRC66 | Did not yield sequence data |
| 66KV26F | CTGAACAAAGGCTTAGG | vif-vpr-vpu | Specific primerwalking for DRC66 | Did not yield sequence data |

| | | | | |
|---|---|---|---|---|
| 66KV2R | ATTTTTGGATGTCTGCT | vif-vpr-vpu | Specific primerwalking for DRC66 | Did not yield sequence data |
| 66KV31F | TGAAGGGGATACAGAGGAA | vif-vpr-vpu | Specific primerwalking for DRC66 | Yielded sequence data |
| 66KV31R | ACAATTTTCCTATACCA | vif-vpr-vpu | Specific primerwalking for DRC66 | Yielded sequence data |
| 66KV33F | TTGTAATGGTATAGGAAAA | vif-vpr-vpu | Specific primerwalking for DRC66 | Did not yield sequence data |
| 66KV33R | TGATGCACAAAATAGAGTAG | vif-vpr-vpu | Specific primerwalking for DRC66 | Yielded sequence data |
| 66KV33Rb | TRATGCACAAAATARAGTAG | vif-vpr-vpu | Specific primerwalking for DRC66 | Yielded sequence data |
| 66KV34R | TCTCTCTCATATGCTTTAGC | vif-vpr-vpu | Specific primerwalking for DRC66 | Yielded sequence data |
| 66KV3F | CAAAAAGAGCTAATGGATGG | vif-vpr-vpu | Specific primerwalking for DRC66 | Yielded sequence data |
| 66KV4F | GAAAGCAGACATCCAAA | vif-vpr-vpu | Specific primerwalking for DRC66 | Did not yield sequence data |
| 66KV4R | CCTGTATACAAACCCCAA | vif-vpr-vpu | Specific primerwalking for DRC66 | Yielded sequence data |
| 66KV6F | CCCATTAGGGGATGCTAA | vif-vpr-vpu | Specific primerwalking for DRC66 | Yielded sequence data |
| 66KV6R | GAGACTCCATGACCCAG | vif-vpr-vpu | Specific primerwalking for DRC66 | Did not yield sequence data |
| 66KV7F | GGGTTTGTATACAGGAGA | vif-vpr-vpu | Specific primerwalking for DRC66 | Yielded sequence data |
| 66KV8F | GGAGAGGGAAAAGGTATAGC | vif-vpr-vpu | Specific primerwalking for DRC66 | Yielded sequence data |
| 66KV8R | CTTATGGCAGATTCTGAA | vif-vpr-vpu | Specific primerwalking for DRC66 | Did not yield sequence data |
| 66KV9R | GGCTTTCCTTATGGCAGA | vif-vpr-vpu | Specific primerwalking for DRC66 | Yielded sequence data |
| CH101F | AAAGAAAAATAGACTGGTTAAT | vif-vpr-vpu | Specific primerwalking for DRC66 | Yielded sequence data |
| CH101R | CCTCTGTATCCCCTTCA | vif-vpr-vpu | Specific primerwalking for DRC66 | Yielded sequence data |
| CH102F | AAGTGTTGCTTTCATTGT | vif-vpr-vpu | Specific primerwalking for DRC66 | Yielded sequence data |
| CH102R | CTGTCTCCGCTTCTTC | vif-vpr-vpu | Specific primerwalking for DRC66 | Yielded sequence data |
| CH103F | TTTCTGAACAAAGGCTTAGG | vif-vpr-vpu | Specific primerwalking for DRC66 | Did not yield sequence data |
| CH103R | GATGATCCTCACTGCYT | vif-vpr-vpu | Specific primerwalking for DRC66 | Did not yield sequence data |
| CH105F | AATAGTAGATGTAATGGTAGC | vif-vpr-vpu | Specific primerwalking for DRC66 | Yielded sequence data |
| CH105R | ATATATRCTATGGTCCACAC | vif-vpr-vpu | Specific primerwalking for DRC66 | Did not yield sequence data |
| CH106F | TGATAGTAGCACTAATYATAG | vif-vpr-vpu | Specific primerwalking for DRC66 | Did not yield sequence data |
| CH106R | AGTCTATTTTTCTTTGCCTTA | vif-vpr-vpu | Specific primerwalking for DRC66 | Yielded sequence data |
| CH107F | GTCAGACATTTTCCTAGA | vif-vpr-vpu | Specific primerwalking for DRC66 | Did not yield sequence data |
| CH107R | CTTATTATAGCTTCAACTCCT | vif-vpr-vpu | Specific primerwalking for DRC66 | Did not yield sequence data |
| CH108F | ATGGATTAGGACAAYATATC | vif-vpr-vpu | Specific primerwalking for DRC66 | Did not yield sequence data |
| CH108R | AGTTGTTGCAAAATTCTTAT | vif-vpr-vpu | Specific primerwalking for DRC66 | Did not yield sequence data |
| CH109F | ATTTCCTATGGCAGGAAG | vif-vpr-vpu | Specific primerwalking for DRC66 | Did not yield sequence data |
| CH109R | AGGATYTTGATGATCCTYAC | vif-vpr-vpu | Specific primerwalking for DRC66 | Did not yield sequence data |
| CH110F | CCAAGCAGTRAGGATCA | vif-vpr-vpu | Specific primerwalking for DRC66 | Did not yield sequence data |
| CH110R | TGCTACCATTACATCTACTA | vif-vpr-vpu | Specific primerwalking for DRC66 | Did not yield sequence data |
| CH1F | CACTATGAAAGCAGACATCC | vif-vpr-vpu | Specific primerwalking for DRC66 | Yielded sequence data |
| CH1R | CTCCTGTATACAAACCCCAA | vif-vpr-vpu | Specific primerwalking for DRC66 | Yielded sequence data |
| CH2F | ATCCCAYTAGGGGATGCT | vif-vpr-vpu | Specific primerwalking for DRC66 | Did not yield sequence data |
| CH2R | TCCATGACCCAGATACCA | vif-vpr-vpu | Specific primerwalking for DRC66 | Did not yield sequence data |
| CH3F | TATCTGGGTCATGGAGTCT | vif-vpr-vpu | Specific primerwalking for DRC66 | Did not yield sequence data |
| CH3R | TCTGCCAGGCCAGGRT | vif-vpr-vpu | Specific primerwalking for DRC66 | Did not yield sequence data |
| CH4F | GGAGAGGGAAAAGGTATAGC | vif-vpr-vpu | Specific primerwalking for DRC66 | Yielded sequence data |
| CH4R | GGCTTTCCTTATGGCAGA | vif-vpr-vpu | Specific primerwalking for DRC66 | Yielded sequence data |
| CH5F | TTGGCACTAACAGCA | vif-vpr-vpu | Specific primerwalking for DRC66 | Yielded sequence data |
| CH5R | GGCTTGTTCCATCTATCC | vif-vpr-vpu | Specific primerwalking for DRC66 | Yielded sequence data |
| CH66F | TTATGGAAAACAGATGGC | vif-vpr-vpu | Specific primerwalking for DRC66 | Yielded sequence data |
| CH66R | TTTCACTAAACTGTTCCAT | vif-vpr-vpu | Specific primerwalking for DRC66 | Yielded sequence data |
| CH67F | TGCAACAACTACTGTTTATT | vif-vpr-vpu | Specific primerwalking for DRC66 | Yielded sequence data |
| CH67R | CTACTGAATCCATTCCTTG | vif-vpr-vpu | Specific primerwalking for DRC66 | Yielded sequence data |
| CH68F | TAGATCCTAATCTAGAGCC | vif-vpr-vpu | Specific primerwalking for DRC66 | Yielded sequence data |
| CH68R | AATGAAAGCAACACTTTTTA | vif-vpr-vpu | Specific primerwalking for DRC66 | Yielded sequence data |
| CH69F | CTTTCTGAACAAAGGCTTA | vif-vpr-vpu | Specific primerwalking for DRC66 | Did not yield sequence data |
| CH69R | ATTTTGATGATCCTCACTG | vif-vpr-vpu | Specific primerwalking for DRC66 | Did not yield sequence data |
| CH6F | GGGAGCCATTTAATGAATGG | vif-vpr-vpu | Specific primerwalking for DRC66 | Yielded sequence data |
| CH6R | TGAAGCCATRGTCTAGGAA | vif-vpr-vpu | Specific primerwalking for DRC66 | Yielded sequence data |
| CH70F | CAGTGAGGATCATCAARATC | vif-vpr-vpu | Specific primerwalking for DRC66 | Did not yield sequence data |
| CH70R | TACTCCTAATGCTRTTAAAT | vif-vpr-vpu | Specific primerwalking for DRC66 | Did not yield sequence data |
| CH71F | AATCYTATATCAAAGCAGTAA | vif-vpr-vpu | Specific primerwalking for DRC66 | Yielded sequence data |
| CH71R | TACTATCAATGCTSCYACTC | vif-vpr-vpu | Specific primerwalking for DRC66 | Did not yield sequence data |
| CH72F | CAGCATTGATAGTAGCAC | vif-vpr-vpu | Specific primerwalking for DRC66 | Did not yield sequence data |
| CH72R | TTTCTTTGTYTTAMCAATTTC | vif-vpr-vpu | Specific primerwalking for DRC66 | Did not yield sequence data |
| CH72Rb | TTTCTTTGCCTTAACAGTTTC | vif-vpr-vpu | Specific primerwalking for DRC66 | Did not yield sequence data |
| CH73F | GGACCATAGYATATATAGAA | vif-vpr-vpu | Specific primerwalking for DRC66 | Yielded sequence data |
| CH73R | TTCTGCTCTTTCYCTAAT | vif-vpr-vpu | Specific primerwalking for DRC66 | Yielded sequence data |
| CH7F | GCATGAAGCTGTYAGACAC | vif-vpr-vpu | Specific primerwalking for DRC66 | Did not yield sequence data |
| CH7Fb | GCATGAAGYTGTCAGACAT | vif-vpr-vpu | Specific primerwalking for DRC66 | Did not yield sequence data |
| CH7R | TAGCTTCAACTCCTGYCCA | vif-vpr-vpu | Specific primerwalking for DRC66 | Did not yield sequence data |
| CH8F | GGCATCTTGCTCTTTTGG | vif-vpr-vpu | Specific primerwalking for DRC66 | Yielded sequence data |
| CH8R | TTGCTTCTCTCCACACA | vif-vpr-vpu | Specific primerwalking for DRC66 | Yielded sequence data |
| TI3F | AGCATGAAGYTGTCAGACATT | vif-vpr-vpu | Specific primerwalking for DRC66 | Yielded sequence data |
| TI3R | CCCAAGTRTCCCCATAGGTT | vif-vpr-vpu | Specific primerwalking for DRC66 | Yielded sequence data |
| TI4F | TTGGCTCCATRGMTTAGGACA | vif-vpr-vpu | Specific primerwalking for DRC66 | Yielded sequence data |
| TI4R | TATAGCTTCAACTCCTGYCCA | vif-vpr-vpu | Specific primerwalking for DRC66 | Yielded sequence data |
| TI5F | CGTGCTTTCTGAACAAAGGCT | vif-vpr-vpu | Specific primerwalking for DRC66 | Did not yield sequence data |
| TI5R | TCCTCACTGCTTKGAGGAGC | vif-vpr-vpu | Specific primerwalking for DRC66 | Did not yield sequence data |
| TI6F | AGGCATTTCCTATGGCAGGA | vif-vpr-vpu | Specific primerwalking for DRC66 | Yielded sequence data |
| TI6R | TTGATGATCCTYACTGCTTKGA | vif-vpr-vpu | Specific primerwalking for DRC66 | Yielded sequence data |
